# Combinatorial transcriptional regulation establishes subtype-appropriate synaptic properties in auditory neurons

**DOI:** 10.1101/2023.10.18.562788

**Authors:** Isle Bastille, Lucy Lee, Cynthia Moncada-Reid, Wei-Ming Yu, Austen Sitko, Andrea Yung, Lisa Goodrich

**Affiliations:** Department of Neurobiology, Harvard Medical School, Boston, MA; Department of Biology, Loyola University Chicago, Chicago, IL; Genentech, South San Francisco, CA

**Keywords:** combinatorial codes, transcriptional regulation, synaptic diversity, neuronal differentiation, neuronal identity, neuronal subtypes, auditory, hearing

## Abstract

Neurons develop diverse synapses that vary in content, morphology, and size. Although transcriptional regulators of neurotransmitter identity have been identified, it remains unclear how other synaptic features are patterned among neuronal subtypes. In the auditory system, glutamatergic synaptic properties vary across three subtypes of spiral ganglion neurons (SGNs) that collectively encode sound information. Here, we show that a combinatorial Maf transcription factor code establishes SGN subtype identity and shapes both shared and subtype-appropriate synaptic properties. We find that *c-Maf* and *Mafb* have independent and opposing effects on synaptic morphology and auditory function, while also acting redundantly to impart subtype identities and drive synaptic differentiation needed for normal auditory responses. Additionally, *c-Maf* and *Mafb* are expressed at different levels across subtypes and regulate subtype-appropriate gene expression in a dose-dependent manner. Thus, functional diversity of Maf family members enables flexible and robust control of gene expression needed to generate synaptic heterogeneity across neuronal subtypes.

## Introduction

Robust and accurate synaptic transmission is essential for perception and behavior. The task of relaying a huge range of sensory information is divided into sensory neuron subpopulations that each encode distinct features of incoming stimuli. To this end, sensory neuron synapses differ widely in their composition, size, and morphology, thereby enabling finely tuned differences in their signaling properties (Petitpré et al., 2022). This large range of synaptic features is in part dictated and maintained by the distinct transcriptional instructions found in each neuronal subtype. Although many synaptic proteins have been identified and characterized for their effects on signal transmission, it is unclear how groups of proteins are co-regulated to fine-tune synaptic properties among closely related neuronal subtypes.

In the periphery of the auditory system, spiral ganglion neurons (SGNs) relay complex sound information from mechanosensitive hair cells to the brain via highly specialized ribbon synapses (Shrestha and Goodrich, 2019). Approximately 5% of the total SGN population are Type II SGNs, which synapse with outer hair cells (OHCs) and are thought to relay information about noxious sound (Zhang and Coate, 2017). Meanwhile, Type I SGNs form synapses with inner hair cells (IHCs) to encode the frequency, intensity and timing of sounds (**Figure S1**).

Presynaptic densities, known as ribbons, localize to the basolateral pole of the IHC and cluster synaptic vesicles that release glutamate onto receptors in SGN postsynaptic densities (**Figure 1A**). Sound frequency is encoded tonotopically, with SGNs in the apex of the cochlea responding to lower frequency sounds and SGNs in the base responding to higher frequencies. Most SGNs receive input from one IHC and each IHC signals to approximately 10-20 SGNs depending on tonotopic location (Perkins and Morest, 1975; Ryugo, 1992; Spoendlin, 1985). At any one position, sound intensity is captured by an action potential rate code, with louder sounds eliciting more spikes. A single SGN is best tuned to a certain frequency and can only encode information about a small proportion of the range of sound intensities found in the environment. However, as a population, SGNs can tile a huge dynamic range of frequencies and intensities with high fidelity and speed.

**Figure 1.**
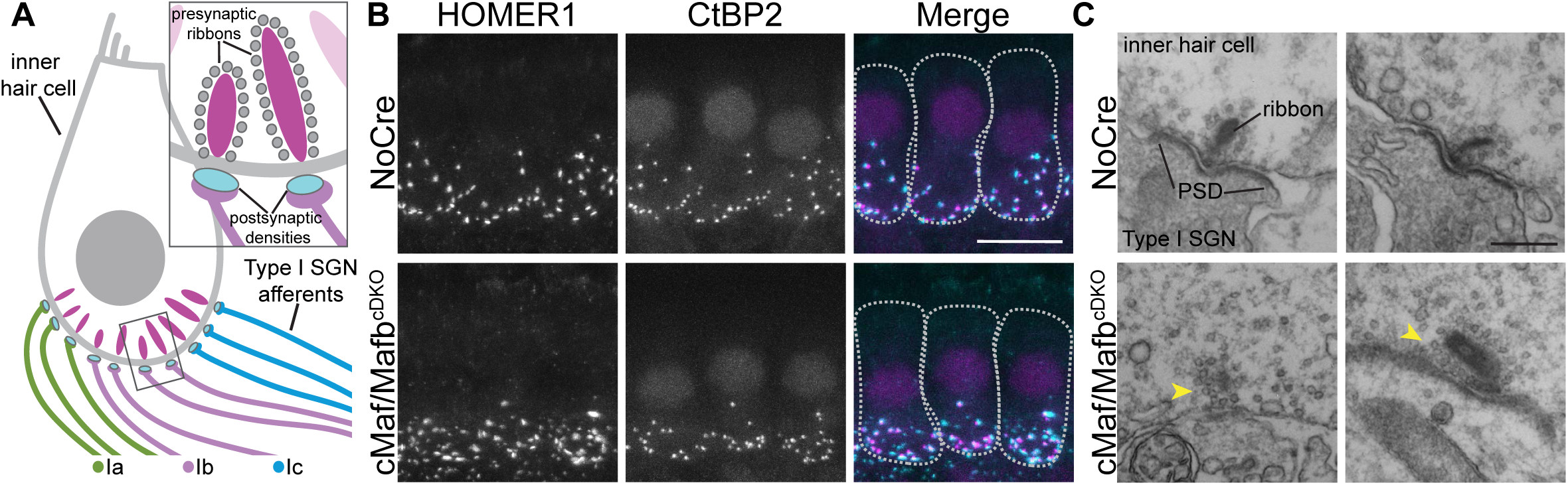
Aberrant synaptic morphology in *cMaf:Mafb* cDKO mice. **(A)** Schematic depicting cross section through an inner hair cell and its innervation by Ia, Ib, and Ic spiral ganglion neuron (SGN) afferents. Inset shows close up view of two ribbon synapses. **(B)** Wholemount immunostaining for the synaptic proteins HOMER1 and CtBP2 (cyan and magenta in merge, respectively) in the 16kHz region of cochleae from control (NoCre, *Mafb^fl/fl^;c-Maf^fl/fl^*, N=4) and *c-Maf:Mafb* double knockouts (c-Maf/Mafb^cDKO^, *bhlhe22^Cre/+^;Mafb^fl/fl^;c-Maf^fl/fl^,* N=4). Hair cells were stained for Calb2 (not shown) and are outlined in dashed lines in the merge panels. Scale bar= 10 μm. **(C)** Electron micrographs of ribbon synapses in control animals (NoCre, *Mafb^fl/fl^;c-Maf^fl/fl^*, N=1, n=3) reveal electron-dense ribbons decorated with synaptic vesicles that are opposed to a prominent postsynaptic density (PSD) on a single Type I SGN. In c-Maf/Mafb double knockout animals (c-Maf/Mafb^cDKO^, *bhlhe22^Cre/+^;Mafb^fl/fl^;c-Maf^fl/fl^,* N=1, n=3), irregularly sized and malformed pre-synaptic ribbons (yellow arrowheads) are docked opposite diffuse and structurally abnormal PSDs in the opposing SGN terminals. In all panels, the inner hair cell is on top and the SGN is on the bottom. Scale bar= 200 nm. See also Figure S1.

Although IHC-SGN synapses are uniformly glutamatergic, Type I SGNs exhibit stereotyped distributions in their physiological and synaptic properties that shape how sound information is communicated to the central nervous system (Shrestha and Goodrich, 2019). At the physiological level, Type I SGNs exhibit different thresholds and spontaneous firing rates (SR) that allow them to capture the complete range of sound intensities found in the environment (Kiang, 1965; Liberman, 1982). Some of this functional diversity likely reflects intrinsic differences among the Type I SGNs. There are three molecularly defined subtypes of Type I SGNs, known as the Ia, Ib, and Ic subtypes, that differ in expression of many transcription factors, ion channels, and synaptic genes (Petitpré et al., 2018; Shrestha et al., 2018; Sun et al., 2018). Ia, Ib, and Ic SGNs match the anatomical features of the physiologically defined SGN subtypes, exemplified by the orderly arrangement of their processes along the basolateral surface of the IHC, with Ia SGN processes in the same position as high SR SGNs, Ic SGN processes in the low SR SGN position, and Ib SGNs in the middle, where medium SR SGN processes reside (Shrestha et al., 2018) (**Figure 1A**). Likewise, genetically labeled Ic SGNs consistently exhibit low spontaneous firing rates when recorded from *in vitro* (Siebald et al., 2023). On the other hand, genetically labeled Ia/Ib SGNs show extensive variability in their firing rates (Siebald et al., 2023), possibly due to differences in their local connectivity. Indeed, individual IHC-SGN synaptic structures show extensive heterogeneity in composition, size, and morphology, and in presynaptic calcium channel density (Hu et al., 2020; Liberman et al., 2011; Liberman and Liberman, 2016; Michanski et al., 2019; Moser et al., 2023; Payne et al., 2021). Some of these differences are linked to subtype identity. For instance, Type Ia SGNs make synapses with larger ribbons and smaller glutamate receptor puncta than the Type Ib and Ic SGNs, again correlating with previously described electrophysiological subtypes (Shrestha et al., 2018). This suggests that synaptic diversity in SGNs is in part dictated by distinct transcriptional networks and contributes to differences in their functional output. By identifying and analyzing the effects of transcription factors that impart subtype-specific differences in synaptic properties, we can better understand how intrinsic differences in subtype identity influence sensory neuron functional diversification.

Diverse SGN subtypes develop from neuronal progenitors in the otocyst and acquire their mature identities through the sequential activity of transcription factors (Goodrich, 2016). Early in development, Gata3 guides neuronal progenitors towards an auditory fate and promotes their differentiation, including guidance of peripheral processes towards hair cells (Appler et al., 2013; Pata et al., 1999). As differentiation progresses, SGNs begin to produce Mafb, which acts downstream of Gata3 and is required for the formation of morphologically normal synapses (Yu et al., 2013). Meanwhile, Runx1 guides the proper specification and maintenance of SGN subtype identity. Removing *Runx1* results in the depletion of Ib and Ic SGNs in favor of Ia SGNs (Shrestha et al., 2023). Following this shift in identity, more neurons form Ia-like synapses. Thus, in *Runx1* conditional knock-out mice, postsynaptic glutamate receptor clusters on the modiolar side of the IHC, i.e. where Ib and Ic SGNs form synapses, are larger in size, matching the cluster sizes normally seen among Ia synapses. The formation of synapses with both subtype-specific and variable properties seems to involve dynamic changes in the position, shape, and size of pre- and post-synaptic elements that begin around birth and continue through the first month of life (Liberman and Liberman, 2016). The final pattern is shaped by thyroid hormone signaling and is maintained by inputs from the olivocochlear efferent system (Coate et al., 2019; Sendin et al., 2007; Yin et al., 2014). However, the terminal effector programs that are deployed to determine and maintain SGN subtype-specific synaptic properties are unknown.

The basic leucine zipper transcription factors c-Maf and Mafb are excellent candidates for regulation of SGN synaptic features. In other systems, Maf family members control terminal cell differentiation by inducing and maintaining identity-specific features such as the production of glucagon by pancreatic cells (Yang and Cvekl, 2016). Deletion of *Mafb* from SGNs in mice disrupted development of the postsynaptic density, resulting in reduced auditory responses (Yu et al., 2013). However, many functional synapses remained, raising the possibility of regulation by another family member. Like *Mafb*, *c-Maf* is expressed downstream of Gata3 in SGNs, but its role has not been determined (Appler et al., 2013; Yu et al., 2013). Work in the somatosensory system demonstrated that c-Maf is essential for the development of vibration-sensitive neurons, acting in part through the regulation of ion channels needed for mature function (Wende et al., 2012). In addition, cultured cortical interneurons from *c-Maf* knockout animals form more synapses, while those from *Mafb* knockouts form fewer, emphasizing the possibility that c-Maf and Mafb have independent roles that could expand their combined effect on synaptic heterogeneity (Pai et al., 2019). c-Maf and Mafb can act cooperatively through dimerization with each other and with other transcription factors (Pogenberg et al., 2014; Rodríguez-Martínez et al., 2017; Suda et al., 2014; Yang and Cvekl, 2016). c-Maf and Mafb have also been implicated in interneuron fate diversification (Pai et al., 2020). Therefore, combinations of c-Maf and Mafb could both influence neuronal identities and elicit a variety of differentiation programs, including genes needed to make diverse synapses.

In this study, we investigated the role of c-Maf and Mafb in the establishment of SGN peripheral synaptic properties. Analysis of anatomical and functional phenotypes as well as gene expression changes in single and double conditional knock-out mice suggests that a different combination of Maf effectors acts in each SGN subtype to establish diverse features, including the acquisition of subtype-appropriate synaptic properties. Our findings suggest a model where different levels of closely-related yet functionally diverse transcription factors execute both shared and subtype-appropriate programs of differentiation needed for nervous system function.

## Results

### SGN synaptic morphology is disrupted in *cMaf:Mafb* cDKO mice

Given the nature of the *Mafb* phenotype (Yu et al., 2013) and the evidence that c-Maf and Mafb both influence cortical interneuron synapse number (Pai et al., 2019), we hypothesized that Mafb and c-Maf work together to regulate SGN synaptic development and function. To test how c-Maf and Mafb contribute to SGN synaptic properties, we stained for pre- and postsynaptic proteins in *c-Maf:Mafb* double mutant mice. The *bhlhe22*^Cre^ driver was crossed to *c-Maf^f^*^lox/flox^;*Mafb*^flox/flox^ mice to generate *c-Maf:Mafb* conditional double knockout mice (cDKO, *bhlhe22^Cre/+^;c-Maf^flox/flox^;Mafb^flox/flox^*, N=4). In the inner ear, *bhlhe22^Cre^* is exclusively active in neurons (Druckenbrod and Goodrich, 2015; Ross et al., 2010). Although *bhlhe22^Cre^* also mediates recombination in olivocochlear efferents (Appler et al., 2013), neither *c-Maf* nor *Mafb* is detected in these neurons (Frank et al., 2023) (**Figure S1A-B**). Analysis of overall cochlear wiring revealed disorganized innervation of outer hair cells by Type II SGNs (**Figure S1C,D**). However, there was no obvious change in the number or position of Type I SGNs or of the organization of their peripheral processes. We therefore proceeded to assess synaptic connectivity among Type I SGNs and inner hair cells (IHCs) by staining wholemount cochlea preparations for CtBP2 and HOMER1 to visualize presynaptic ribbons and postsynaptic densities, respectively.

Synapses were qualitatively abnormal in cDKO cochlea. As compared to controls, the synaptic puncta appeared more variable in size and distribution in cDKOs, with irregular aggregates of small and large synaptic puncta (**Figure 1B**). This phenotype was reflected in structural abnormalities at the synapse. Electron micrographs revealed that cDKO synapses had abnormally small and large ribbons and less well-defined, dysmorphic postsynaptic densities (**Figure 1C**). These results suggest that c-Maf and Mafb may cooperate to establish mature SGN synaptic properties.

### cMaf and Mafb have both overlapping and opposing effects on SGN synaptic phenotypes

To determine how c-Maf and Mafb each contribute to the cDKO synaptic phenotype, we stained for pre- and postsynaptic proteins in *c-Maf* and *Mafb* single mutant mice made using the same Cre driver. We generated c-Maf (*c-Maf^cKO^*: *bhlhe22^Cre/+^;c-Maf^flox/flox^;Mafb^flox/+^,* N=9) and Mafb (*Mafb*^cKO^: *bhlhe22^Cre/+^;c-Maf^flox/+^;Mafb^flox/flox^*, N=7) knockouts, as well as control (NoCre: *c-Maf^f^*^lox/flox^;*Mafb*^flox/flox^, N=15), and conditional double knockout (cDKO: *bhlhe22^Cre/+^;c-Maf^flox/flox^;Mafb^flox/flox^*, N=10) littermates, which allowed us to analyze the relative contributions of these two transcription factors as well as the effects of the combined loss of both within the same cross. To compare littermates of all four genotypes, within these experiments, the single mutants in these stains lack a copy of the other factor. In addition, we stained for the GluA2 glutamate receptor subunit to gain additional insights into the nature of the effect on the post-synaptic density (PSD).

This analysis revealed that c-Maf and Mafb have both independent and overlapping effects on cochlear synapses. As previously observed, synaptic puncta were highly abnormal in the cDKO cochlea, whereas the single mutants exhibited subtle yet complementary changes in synaptic morphology. In cDKOs, the post and pre-synaptic puncta were disorganized and variably sized, though the pre- and post-synaptic puncta still generally apposed each other (**Figure 2A**). Synapses were grossly normal in the single mutants, but the GluA2 puncta appeared larger in c-Maf^cKO^ cochleae and smaller in Mafb^cKO^ cochleae compared to controls. To quantify postsynaptic differences, we created three-dimensional reconstructions of GluA2 puncta in the 16 kHz region of control (NoCre), c-Maf^cKO^, Mafb^cKO^, and cDKO cochleae. This is the region of highest frequency sensitivity in mice and accordingly has the most synaptic puncta (Meyer et al., 2009). To account for any technical differences across experiments, we normalized volume measurements by the median of the control values within each batch. Quantification confirmed opposing effects on synaptic puncta volume in c-Maf^cKO^ and Mafb^cKO^ mice. As in controls, puncta volumes ranged widely in each single mutant strain. However, this distribution was shifted significantly towards larger GluA2 puncta volumes in c-Maf^cKO^ mice (Kruskal-Wallis with Bonferroni adjusted posthoc-Dunn, p=2.85E-23) and towards smaller volumes in Mafb^cKO^ mice (Kruskal-Wallis with Bonferroni adjusted posthoc-Dunn, p=4.17E-02) compared to controls (**Figure 2B**). Median volumes per mouse also trended in opposite directions, with median GluA2 volumes in c-Maf^cKO^ mice significantly larger than those in Mafb^cKO^ mice (**Figure 2C**) (Kruskal-Wallis with Bonferroni adjusted posthoc-Dunn, p=0.033). Although median punctum volume in c-Maf^cKO^ mice showed a trend to be larger and Mafb^cKO^ mice showed a trend to be smaller, there was no statistically significant difference for either strain compared to littermate controls (Kruskal-Wallis test with Bonferroni adjusted posthoc-Dunn, p_cMafcKO_=0.100, p_MafbcKO_=1.00), likely due to the high degree of variability in each genotype. Median GluA2 punctum volume was also not significantly different in cDKO animals compared with controls (Kruskal-Wallis with Bonferroni adjusted posthoc-Dunn, p=1.00), as expected given the presence of both very small and very large puncta in the aggregates that formed. Similar effects were observed in the 8kHz region of the cochlea (**Figure S2A-B**) and in unnormalized 16kHz data (**Figure S2C-E**). Although the *c-Maf^cKO^* and *Mafb*^cKO^ mutants were also heterozygous for the other Maf factor, the same basic phenotypes were observed in single knockouts with two full copies of the other Maf factor, i.e. *bhlhe22^Cre/+^;c-Maf^flox/flox^* and *bhlhe22^Cre/+^; Mafb^flox/GFP^* mice (**Figure S2F-G).** Our analysis also suggests that the smaller post-synaptic puncta observed in Mafb^cKO^ animals was previously interpreted to be a loss of synapses (Yu et al., 2013), likely due to differences in the sensitivity of the immunostaining protocol.

**Figure 2.**
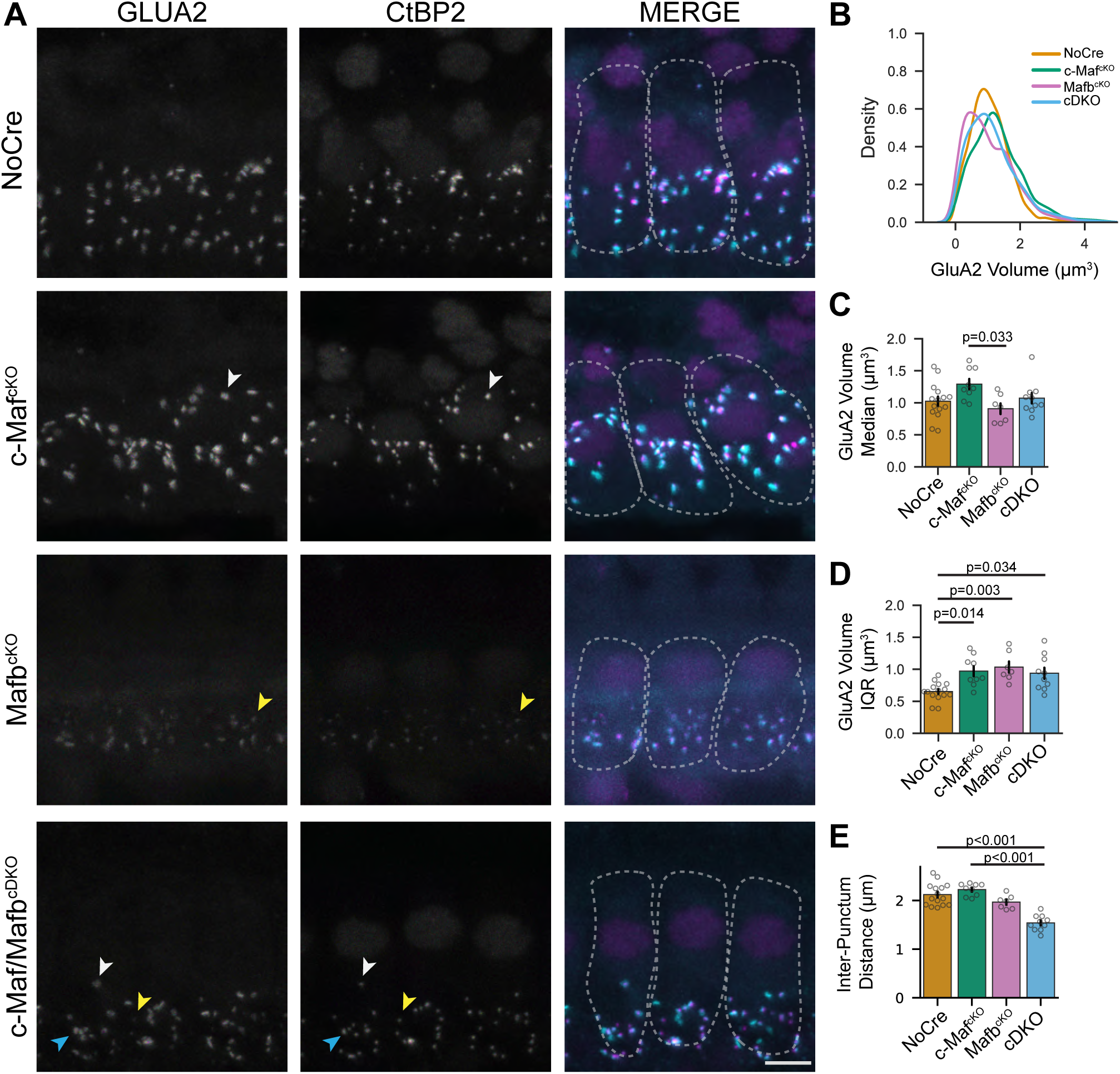
c-Maf and Mafb have overlapping and opposing effects on SGN synapses. **(A)** Wholemount immunostaining for the synaptic proteins GluA2 and CtBP2 in the 16 kHz region of cochleae from control (NoCre, *Mafb^fl/fl^;c-Maf^fl/fl^,* N=15, orange), *c-Maf* knockout (*c-Maf*^cKO^*, bhlhe22^Cre/+^;Mafb^fl/+^;c-Maf^fl/fl^*, N=9, green), *Mafb* knockout (*Mafb*^cKO^, *bhlhe22^Cre/+^;Mafb^fl/fl^;c-Maf^fl/+^*, N=7, magenta), and double knockout (cDKO, *bhlhe22^Cre/+^;Mafb^fl/fl^;c-Maf^fl/fl^,* N=10, blue) littermates. Hair cells were stained for Calb2 (not shown) and are outlined in dashed lines in the merge panels. Volume measurements were normalized by the median of the control values within each stain batch. Scale bar, 5 μm. Post-synaptic puncta opposite pre-synaptic ribbons are often larger in c-MafCKOs (white arrowheads) and smaller in MafbcKOs (yellow arrowheads). In cDKOs, both very large (white arrowheads) and very small (yellow arrowheads) post-synaptic puncta are detected, as are aberrant synaptic aggregates (blue arrowheads). **(B)** Distribution of GluA2 puncta volumes in control (NoCre), *Mafb*^cKO,^ *c-Maf*^CKO^ and cDKO animals (Kruskal-Wallis, p<0.001). **(C)** Median GluA2 puncta volumes per animal across genotypes (Kruskal-Wallis, p=0.030). **(D)** Mean interquartile range of GluA2 puncta volumes per mouse across genotypes (Kruskal-Wallis, p<0.001). **(E)** Mean GluA2 inter-punctum distance per animal across genotypes (Kruskal-Wallis, p<0.001). All error bars are standard error from the mean. See also Figure S2.

Consistent with qualitative observations, the distribution of post-synaptic puncta volumes was broader in cDKO animals (**Figure 2B**). Accordingly, the distribution of GluA2 volumes in cDKO animals had a larger interquartile range than in controls (Kruskal-Wallis with Bonferroni adjusted posthoc-Dunn, p=0.034). Given the opposing effects of c-Maf and Mafb on punctum volume, the larger variance in volumes could be caused by dysregulation of synaptic properties that are normally controlled independently by each Maf factor. In fact, compared to control animals, c-Maf^cKO^ (Kruskal-Wallis with Bonferroni adjusted posthoc-Dunn, p=0.014) and Mafb^cKO^ animals (Kruskal-Wallis with Bonferroni adjusted posthoc-Dunn, p=0.003) also had larger GluA2 punctum volume interquartile ranges compared to control animals; the same differences were observed in unnormlized data (**Figure 2D** and **Figure S2E**). Taken together, these data indicate that GluA2 puncta become larger in c-Maf^CKO^ mice and smaller in Mafb^CKO^ mice, with both larger and smaller puncta present in cDKO animals (arrows, **Figure 2A**). In cDKO animals, the increased presence of very small puncta is balanced by the increased presence of very large puncta, resulting in no change in the median punctum volume per hair cell per animal (**Figure 2B-C**).

Another striking feature of the cDKO phenotype is the abnormal distribution of synaptic puncta along the bottom of the hair cell. Qualitatively, the puncta appeared to be less evenly spread out, clustering towards the center of the IHC’s basal pole. Consistent with this assessment, inter-punctum distances, i.e. the closest distance of each punctum to another punctum, were significantly smaller in cDKO animals compared to controls (Kruskal-Wallis with Bonferroni adjusted posthoc-Dunn, p=2.17E–4). The inter-punctum distances were not changed in Mafb^cKO^ or c-Maf^cKO^ animals relative to controls (Kruskal-Wallis with Bonferroni adjusted posthoc-Dunn, p=1.00 for both comparisons) (**Figure 2E**). Collectively, these studies demonstrate that cDKO animals have more severe synaptic phenotypes than either single mutant and that c-Maf and Mafb affect synaptic structure in different ways.

### Altered auditory responses in *c-Maf* and *Mafb* mutants

Glutamatergic signaling at ribbon synapses is necessary for transmitting auditory information from IHCs to SGNs. To test if the synaptic effects observed in c-Maf and Mafb mutants have functional consequences for the ability to detect sound, we recorded auditory brainstem responses (ABRs) from control, single, and double knockout mice. ABRs are recorded by placing electrodes near the base of the skull of anesthetized mice to measure electric field potentials generated by synchronous firing of neurons at different steps along the ascending auditory pathway in response to sound stimuli. We presented pure tone bursts of 8,16, 32, and 45 kHz each at increasing sound pressure levels from 20 to 90 dB SPL (decibels sound pressure level) in 5 dB increments to control (NoCre: *c-Maf^flox/flox^;Mafb^flox/flox^,* N=14), *c-Maf* knockout (cMaf^cKO^: *bhlhe22^Cre/+^;c-Maf^flox/flox^;Mafb^flox/+^*, N=6), *Mafb* knockout (Mafb^cKO^: *bhlhe22^Cre/+^;c-Maf^flox/+^;Mafb^flox/flox^,* N=11), and conditional double knockout (cDKO: *bhlhe22^Cre/+^;c-Maf^flox/flox^;Mafb^flox/flox^,* N=9) littermates. Qualitatively, the peaks in average ABR waveforms seemed larger in amplitude in c-Maf^cKO^ animals, smaller in Mafb^cKO^ animals, and nearly undetectable for most sound pressure levels in the cDKO animals (**Figure 3A-D**). Auditory sensitivity is measured by identifying the lowest sound pressure level, or threshold, that elicits a brainstem response at each frequency. We only analyzed the ABR responses to 8 and 16 kHz stimuli to avoid confounds of high frequency hearing loss characteristic of certain strains of mice (Kane et al., 2012); indeed, the ABR threshold was elevated for control animals beyond 32 kHz (**Figure 3E**). We also measured the amplitude and latency of the first peak of the ABR response (P1), which reflects the degree and speed of synchronous firing of SGNs (Melcher and Kiang, 1996).

**Figure 3.**
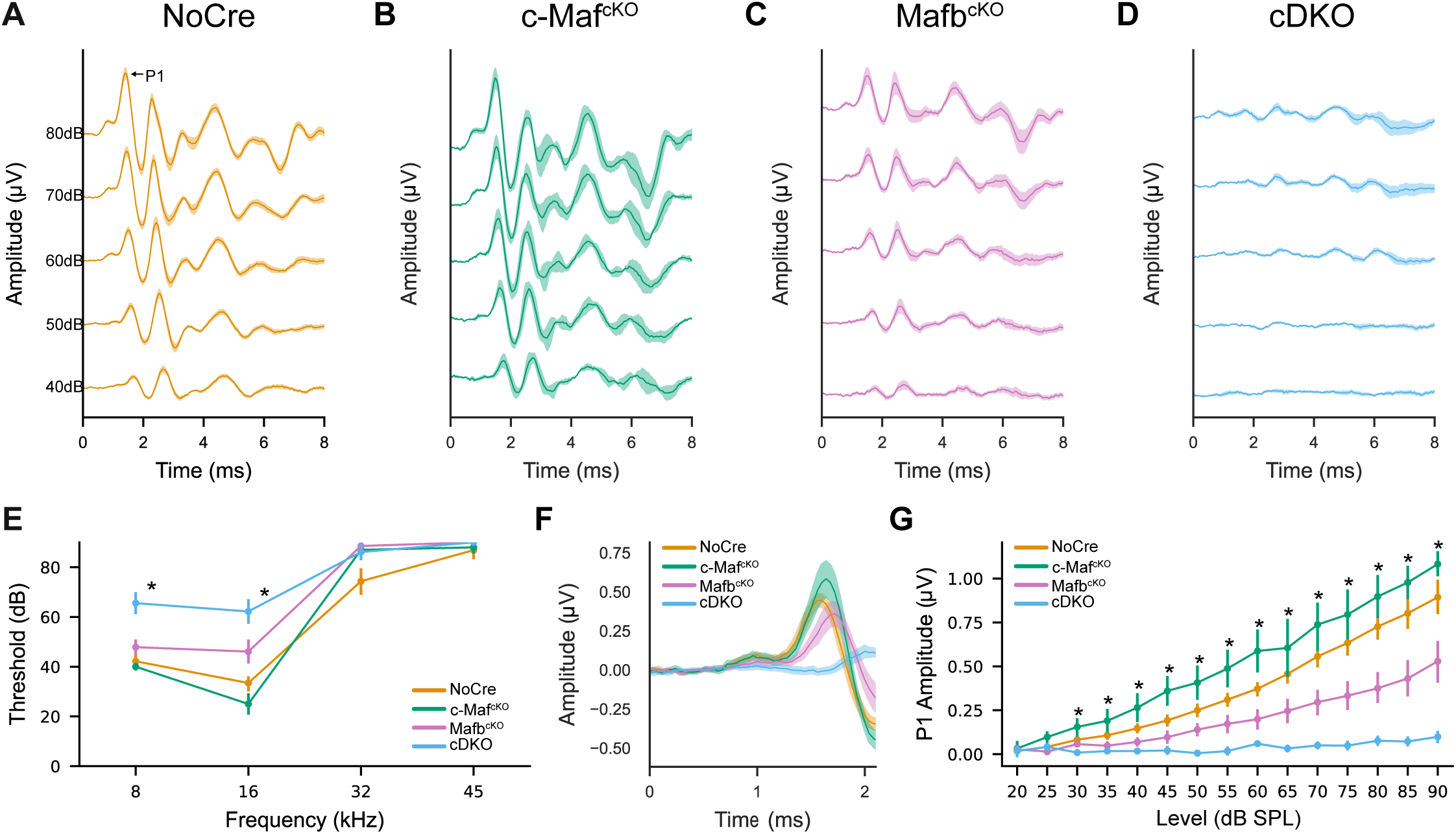
*c-Maf* and *Mafb* have opposing and additive effects on auditory function. Auditory brainstem responses (ABRs) were recorded from 8-12 week old control animals (NoCre, *Mafb^fl/fl^;c-Maf^fl/fl^,* N=14, orange), *c-Maf* knockout (c-Maf^cKO^*, bhlhe22^Cre/+^;Mafb^fl/+^;c-Maf^fl/fl^*, N=6, green), *Mafb* knockout (Mafb^cKO^, *bhlhe22^Cre/+^;Mafb^fl/fl^;c-Maf^fl/+^*, N=11, magenta), and double knockout (cDKO, *bhlhe22^Cre/+^;Mafb^fl/fl^;c-Maf^fl/fl^,* N=9, blue) littermates. **(A-D)** Average ABR waveforms recorded after presentation of a 16 kHz stimulus at different sound intensity levels (in decibels, dB) across genotypes. Standard error shown by shaded bands. **(E)** ABR threshold measurements across frequencies (Kruskal-Wallis, p_8kHz_<0.001, p_16kHz_<0.001, p_32kHz_=0.157, p_45kHz_=0.293). **(F)** Overlaid average ABR waveforms of Peak 1 (P1) across genotypes in response to a 16 kHz, 80 dB SPL (sound pressure level) sound stimulus in control animals (NoCre, orange), *c-Maf* knockout (c-Maf^cKO^, green), *Mafb* knockout (Mafb^cKO^, magenta), and double knockout (cDKO, blue) littermates. **(G)** P1 amplitude across all sound intensities for a 16 kHz stimulus. Comparisons with an asterisk were statistically significant (p<0.05, Kruskal-Wallis). Significant differences amongst groups that had a Kruskal-Wallis p-value<0.05 were followed with a pairwise post-hoc Dunn test. Kruskal-Wallis p-values and pairwise post-hoc Dunn p-values can be found in Supplemental Table 1. All error bars are bootstrapped 68% confidence interval. See also Figure S3.

Consistent with the observed synaptic defects, loss of *c-Maf* or *Mafb* had different effects on ABR thresholds and strength of synchronous SGN firing. There were significant changes in threshold, P1 amplitude, and P1 latency across groups for most sound pressure levels (statistical comparisons in **Supplemental Table 1**). Corroborating previous findings (Yu et al., 2013), Mafb^cKO^ animals had significantly elevated thresholds at 16kHz (Kruskal-Wallis with Bonferroni adjusted posthoc-Dunn, p=0.0355), significantly smaller P1 amplitudes, and delayed latencies compared to controls (**Figure 3E-G**, **Figure S3**). By contrast, c-Maf^cKO^ animals showed no significant threshold shifts (Kruskal-Wallis with Bonferroni adjusted posthoc-Dunn, p=0.240). However, P1 occurred significantly later and trended towards larger amplitudes compared to controls (**Figure 3E-G**, **Figure S3B-D**). cDKO animals had significantly elevated thresholds (Kruskal-Wallis with Bonferroni adjusted posthoc-Dunn, p=1.140E-4) and decreased P1 amplitudes compared to all the other genotypes (**Figure 3E-G**). We did not measure latency in cDKO animals since it was difficult to reliably identify the crest of P1 in these highly aberrant ABRs. There were no significant differences in distortion product otoacoustic emissions (DPOAE) thresholds across genotypes (16kHz, Kruskall-Wallis, p=0.115) (**Figure S3A**). Since DPOAE are indicators of outer hair cell function, these results suggested that the observed changes in ABRs originate with IHC, SGNs, or the synapses that link them.

The exacerbated effects on threshold and P1 amplitudes in cDKO animals suggest that *c-Maf* and *Mafb* act synergistically to control SGN properties needed for synchronous firing in response to IHCs. In further support of this idea, cDKO animals also exhibited higher thresholds and lower P1 amplitudes than Mafb^cKO^ animals. Since the only difference between Mafb^cKO^ animals (*bhlhe22^Cre/+^;Mafb^flox/flox^;c-Maf^flox/+^*) and cDKO littermates (*bhlhe22^Cre/+^;Mafb^flox/flox^;c-Maf^flox/flox^)* was a single copy of *c-Maf,* this result confirms a critical role for c-Maf, despite the relatively mild phenotypes observed in single c-Maf^cKO^ mice. Notably, the ABRs matched predictions from the observed synaptic defects, with smaller GluA2 puncta and smaller responses in Mafb^cKO^ mice *vs.* larger GluA2 puncta and slightly larger responses in c-Maf^cKO^ mice. Further, by comparison to the single mutants, cDKO animals showed more severe synaptic defects and accordingly poor auditory responses. Thus, Mafb and c-Maf seem to have both independent and combinatorial effects on SGN synaptic differentiation and function.

### c-Maf and Mafb are expressed at different levels across SGN subtypes

SGN molecular subtypes express both shared and distinct synaptic genes, including postsynaptic receptors, synaptic adhesion molecules, and potassium, sodium and calcium channels (Petitpré et al., 2018; Shrestha et al., 2018; Sun et al., 2018). The opposing effects seen in the single mutants together with the exacerbated phenotypes in double mutants raised the possibility that c-Maf and Mafb influence subtype-specific programs for synaptic differentiation. In support of this idea, re-analysis of published single cell RNA sequencing (scRNA-seq) datasets showed that there are more *c-Maf* transcripts in Ia SGNs, which express the Ia-enriched gene *Calb2*, and more *Mafb* transcripts in Ib and Ic SGNs, which express the Ib/Ic-enriched gene *Ntng1*, which encodes NetrinG1 (Petitpré et al., 2022; Shrestha et al., 2018) (**Figure 4A-B**). We confirmed differential expression of c-Maf protein across SGN subtypes by double staining for c-Maf and Calb2 in cochlear sections from P27-P30 mice (N=3) (**Figure 4C**). Quantification of staining intensity in 3D reconstructions of individual SGNs showed that c-Maf staining intensity correlated positively with Calb2 staining intensity (R=0.550, p=1.818E-24) (**Figure 4D**). Calb2 is highest in Ia SGNs, lower in Ib, and lowest in Ic SGNs (Shrestha et al., 2018) (**Figure 4A**). Thus, at both the RNA and protein levels, c-Maf is higher in Ia SGNs than in Ib and Ic SGNs. We were unable to measure adult Mafb protein expression because Mafb protein is not detectable in the nucleus after the first postnatal week and available antibodies are not sensitive enough to make quantitative immunohistochemical assessments.

**Figure 4.**
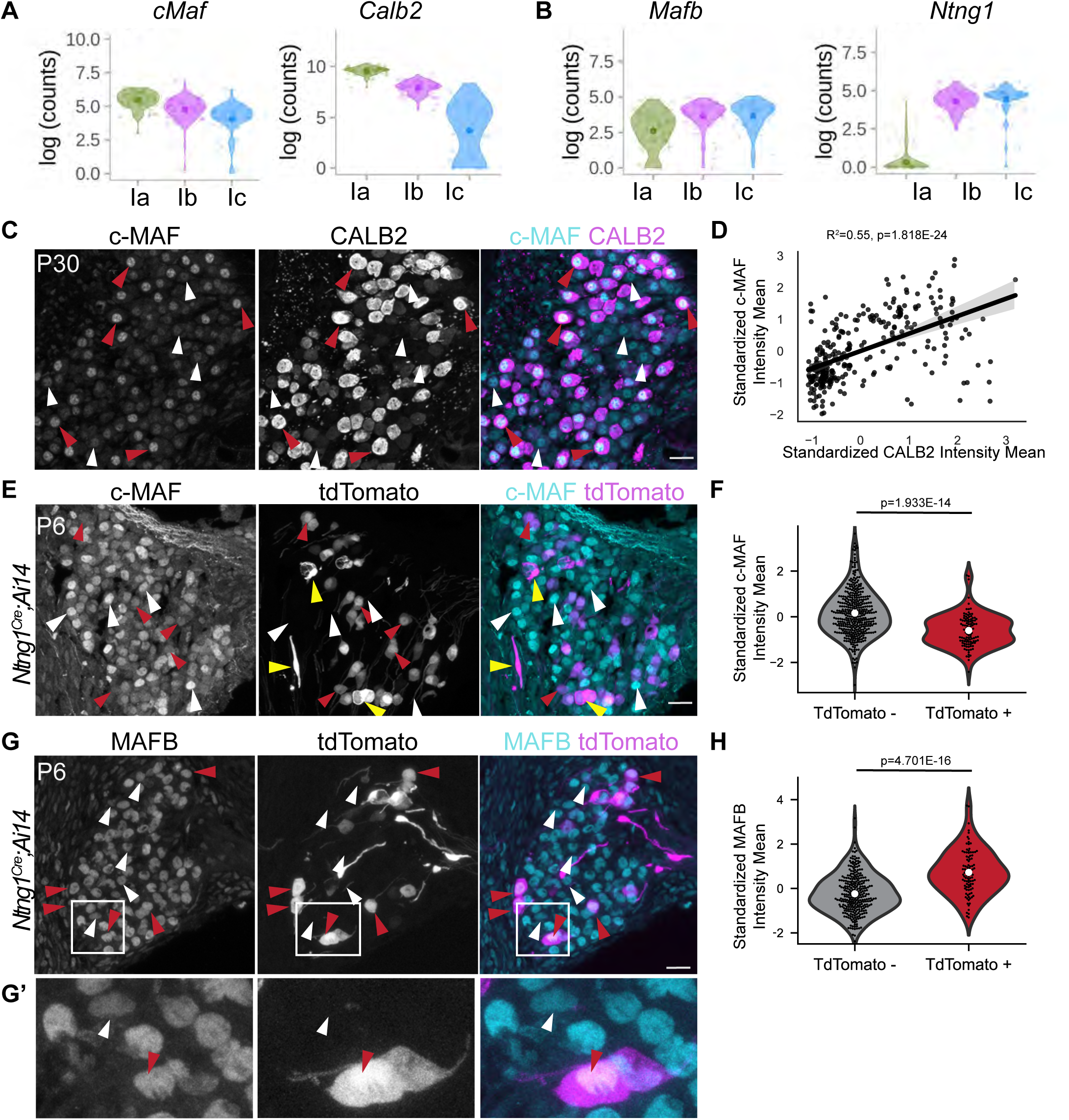
c-Maf and Mafb are expressed at different levels across SGNs subtypes. **(A)** scRNAseq expression profiles of *c-Maf* and a Ia-enriched gene, *Calb2* (Shrestha et al., 2018). **(B)** scRNAseq profiles of *Mafb* and a Ib/Ic-enriched gene *Ntng1* (Shrestha et al., 2018)**. (C)** Immunolabeling of c-Maf and Calb2 in cochlear sections of P28-P30 wildtype mice (N=3). High c-Maf expressing SGNs (red arrowheads) also express abundant Calb2, whereas low c-Maf expressing SGNs (white arrowheads) do not. **(D)** Positive correlation between standardized (z-scored) staining intensities for c-Maf and Calb2 (R^2^=0.55, p<0.001). Shaded error bar is 95% confidence interval of the regression fit. **(E)** Immunostaining for c-Maf (cyan, merge) and tdTomato (magenta, merge) in *Ntng1^Cre/+^;Ai14/+* cells (magenta) in sections through the cochlea of P6 mice (N=4). Examples of c-Maf staining in tdTomato+ (red arrowheads) and tdTomato-(white arrowheads) SGNs are indicated. tdTomato+ SGNs will develop as Ib or Ic SGNs, whereas tdTomato-could develop as any of the three subtypes. C-Maf levels appear lower in presumptive Ib and Ic SGNs. Glia (yellow arrowheads) were excluded by the intensity of tdTomato, smaller cell body size, fried-egg like morphology and lack of Calb2 expression, yellow arrowheads. **(F)** Quantification of standardized c-Maf staining intensity in *Ntng1^Cre/+^;Ai14/+* labeled and unlabeled P6 SGNs. Each dot corresponds to a single reconstructed cell (p=<0.001, Mann-Whitney Rank sum with Bonferroni correction). **(G)** Immunostaining for Mafb (cyan) and tdTomato (magenta) in sections through the cochlea of P6 *Ntng1^Cre/+^;Ai14/+* mice (N=4), with example tdTomato+ and tdTomato-SGNs indicated and shown at higher power in **G’**. Mafb levels appear higher in tdTomato+ SGNs (presumptive Ib and Ic SGNs, red arrowheads) than in tdTomato-SGNs (white arrowheads). **(H)** Quantification of standardized Mafb staining intensity in *Ntng1^Cre/+^;Ai14/+* labeled and unlabeled P6 SGNs. Each dot corresponds to a single reconstructed cell. (p=<0.001, Mann-Whitney Rank sum with Bonferroni correction).

Further analysis demonstrated that c-Maf and Mafb are also expressed differentially across SGN subtypes during synaptogenesis, which occurs during the first two postnatal weeks of life in mice (Coate et al., 2019; Moser et al., 2023). To mark developing SGN subtypes definitively, we used *NetrinG1^Cre/+^*, which selectively marks adult Ib and Ic subtypes when combined with the *Ai14* Cre-dependent tdTomato reporter (RCL-tdT), consistent with its scRNA-seq expression profile (**Figure 4B**). This activity begins postnatally, with a sparse subset of SGNs expressing tdTomato in postnatal day 6 (P6) *NetrinG1^Cre/+^;Ai14/+* mice (**Figure 4E,G**). Since we find nearly complete coverage of Ib and Ic SGNs in adult *NetrinG1^Cre/+^;Ai14/+* mice (Kreeger et al., 2024), any tdTomato labeling at P6 is a reliable indication of a Ib or Ic identity. Although *NetrinG1^Cre/+^* also drives recombination in a subset of myelinating glia at this stage, these cells are readily distinguishable from neurons by the smaller size of their cell bodies, more intense tdTomato expression, and fried-egg-like morphologies (**Figure 4E**). c-Maf and Mafb levels were quantified by measuring mean staining intensities in 3D reconstructions of individual tdTomato+ (Ib/Ic) and tdTomato- (undetermined identity) SGN cell bodies in P6 *NetrinG1^Cre/+^;Ai14/+* mice (N=5) (**Figure 4E,G**). To account for any technical differences in staining intensity across experiments, we standardized (z-scored) the staining intensity for cells in each animal so that the mean staining intensity for each animal is 0 and the standard deviation across cells is 1. Consistent with what was observed at the RNA level in adults (**Figure 4A-B**), developing tdTomato+ Ib and Ic SGNs had lower c-Maf staining intensity (Mann Whitney Rank Sum, statistic=32117.0, p=1.934E-14) (**Figure 4E,F**) and higher Mafb staining intensity (Mann Whitney Rank Sum, statistic=6923.0, p=4.701E-16) (**Figure 4G,H**) than tdTomato-cells. These results suggest that c-Maf and Mafb are already expressed at different levels in SGN subtypes during peak synaptogenesis (Coate et al., 2019; Huang et al., 2012; Yu and Goodrich, 2014). Transcriptomes from P3 SGNs show enrichment of *c-Maf* in developing Ia SGNs whereas *Mafb* was expressed at similar levels in all SGNs at this stage (Petitpré et al., 2018), suggesting that the adult pattern of expression emerges while identities are consolidating and concomitant with synaptic differentiation. Thus, *c-Maf* and *Mafb* could be playing distinct roles in regulating gene expression needed for subtype-specific features of the synapse.

### c-Maf and Mafb have both shared and unique effects on gene expression

The constellation of phenotypes observed in single and double mutants is consistent with the idea that combinations of Mafb and c-Maf can have complex effects on gene expression. To assess gene expression changes conferred by Mafb and c-Maf alone or together, we used single-cell RNA sequencing to compare the transcriptomes of SGNs from *c-Maf* and *Mafb* single and double knockout mice and littermate controls. In these experiments, the single mutants were wild-type for the other Maf factor, thereby allowing us to confidently attribute gene expression changes to the loss of c-Maf, Mafb, or both c-Maf and Mafb. In control transcriptomes, SGNs segregated into the expected Ia, Ib, and Ic clusters, as identified by previously defined marker genes. As previously observed, *c-Maf* and *Mafb* were expressed at different levels across SGN subtypes, with more *c-Maf* in Ia SGNs and more *Mafb* in Ib and Ic SGNs (**Figure 5A**).

**Figure 5.**
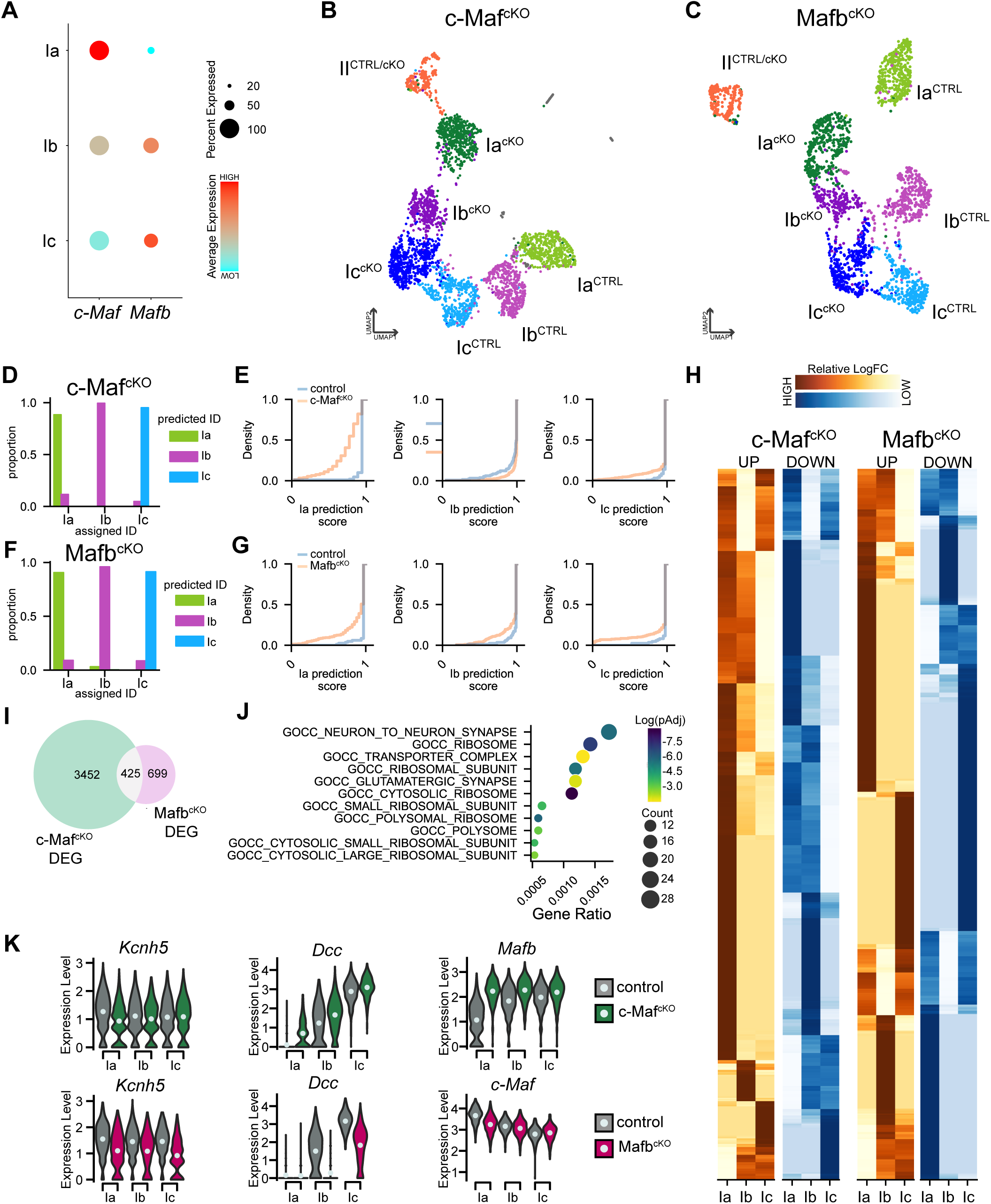
c-Maf and Mafb influence subtype differentiation and regulate overlapping yet distinct sets of synaptic genes. **(A)** Dot plot summarizing relative *c-Maf* and *Mafb* expression across control SGN subtypes, as determined by scRNA-seq. Red denotes higher relative expression and cyan denotes lower relative expression. **(B,C)** UMAP plots summarizing sequencing data of 1669 control SGNs (*bhlhe22^Cre/+^; c-Maf^fl/+^,* N=6) and 1708 c-Maf^cKO^ SGNs (*bhlhe22^Cre/+^; c-Maf^fl/fl^,* N=3) and 1172 control SGNs (*Mafb^fl/+^,* N=3) and 1058 Mafb^cKO^ SGNs (*bhlhe22^Cre/+^; Mafb^fl/GFP^,* N=4). In both single mutants, Ia, Ib, and Ic SGNs can be identified, but they do not cluster with control SGN subtypes. Control and mutant Type II SGNs co-cluster in both datasets. (**D-G**) The proportion of c-Maf^cKO^ (**D**) and Mafb^cKO^ (**F**) cells assigned as each subtype when projected onto control cells relative to the identities that were assigned based on cell type markers. Prediction scores of c-Maf^cKO^ **(E)** and Mafb^cKO^ **(G)** cells assigned to each class compared to withheld control cells. **(H)** Heat maps of relative log fold changes in gene expression of all c-Maf^cKO^ (left) and Mafb^cKO^ DEGs, with upregulated genes in orange and downregulated genes in blue. For each gene, the biggest expression difference (normalized to 1) is illustrated in the most intense colors. **(I)** Relative number of genes that are differentially expressed in either or both Mafb^cKO^ and c-Maf^cKO^ animals. **(J)** Cellular component gene ontology analysis of 425 overlapping DEG in Mafb^cKO^ and c-Maf^cKO^ animals. **(K)** Expression of *Kcnh5, Dcc, c-Maf,* and *Mafb* in Mafb^cKO^ and c-Maf^cKO^ animals compared to controls. *Kcnh5* decreases in both c-Maf^cKO^s and Mafb^cKO^s, whereas *Dcc* increases in c-Maf^cKO^s and decreases in Mafb^cKO^s. *Mafb* levels are significantly increased in all c-Maf^cKO^ SGN subtypes, whereas *c-Maf* levels are significantly decreased in *Mafb^cKO^* Ia SGNs. See also Figure S5.

Analysis of single mutant transcriptomes confirmed independent roles for *c-Maf* and *Mafb* in Ia, Ib, and Ic SGN subtypes. We performed unsupervised clustering analysis on c-Maf^cKO^ (*bhlhe22^Cre/+^;c-Maf^fl/fl^*, N=3 mice, n= 1708 neurons) and control SGNs (*c-Maf^fl/fl^*, N=3 mice, n= 1669 neurons) and independently on SGNs from *Mafb^cKO^* mice (*bhlhe22^Cre/+^;Mafb^fl/GFP^*, N=3, n= 1058 neurons) and their littermate controls (*Mafb^fl/GFP^*, N=3 mice, n= 1172 neurons). Each comparison revealed control Ia, Ib, and Ic clusters and three knockout clusters that were separate from the controls but still corresponded to the three SGN subtypes, as identified by expression of known marker genes. Meanwhile, Type II control and cKO SGNs co-clustered (**Figure 5B,C**). To verify that cKO SGNs were still assuming Ia, Ib and Ic subtype identities, we evaluated cell classification computationally by comparing individual cKO and control transcriptomes. Specifically, we utilized a canonical correlation alignment analysis in Seurat that projects cells onto an anchor dataset. In this case, the anchor dataset for each comparison was the control SGNs. Indeed, cKO SGNs projected onto the same identity classes that we had assigned based on subtype marker expression (**Figure 5D,F**). Thus, the changes in gene expression that separate cKO SGNs from control SGNs do not fundamentally alter their identity. Further, these data indicate that c-Maf and Mafb regulate gene expression in all three Type I SGN subtypes, consistent with their pan-SGN expression.

Since Type I cKO SGNs could be assigned Ia, Ib and Ic identities, the separation in control and cKO clusters is likely driven by abnormal execution of downstream gene expression programs. Given their complementary patterns of expression and distinct effects on synapse morphology and auditory responses, we reasoned that c-Maf and Mafb might have different effects on synaptic gene expression in each subtype. To investigate this possibility, we first measured how well mutant subtype classes aligned with the assigned identity, as indicated by a prediction score. A prediction score of 1 indicates high certainty that the cell was properly projected into the right class. When 25% of the control SGNs were withheld and then projected onto the remaining 75%, most of the cells in each class had prediction scores close to 1 (**Figure 5E**). By contrast, when projected onto each control SGN subtype, c-Maf^cKO^ cells had lower prediction scores for Ia SGNs (**Figure 5E**) and Mafb had lower prediction scores for all three subtypes with a pronounced lower tail for the Ic SGNs (**Figure 5G**). These results suggest that c-Maf has a particularly strong effect on gene expression in Ia SGNs, whereas Mafb affects all three subtypes, but especially the Ic SGNs.

Comparative bioinformatic analysis of gene expression changes in each cKO confirmed that c-Maf and Mafb contribute differentially to Ia, Ib, and Ic transcriptomes, with prominent effects on synapse-related genes. We identified 3,877 genes differentially expressed genes (DEG) in c-Maf^cKO^ SGNs and 1,124 DEG in *Mafb^cKO^* SGNs when compared to their respective controls. 425 genes were changed in both datasets (**Figure 5I**). Among those shared genes, 235 changed in the same direction in each mutant and 190 changed in opposite directions. For instance, *Kcnh5,* which encodes for a voltage-gated potassium channel, was significantly downregulated in both cMaf^cKO^ and Mafb^cKO^ SGNs (**Figure 5K**). Meanwhile, *Dcc*, which encodes a Netrin-1 receptor and is enriched in Ib/Ic SGNs, was upregulated in c-Maf^cKO^ SGNs and downregulated in Mafb^cKO^ neurons. Thus, c-Maf and Mafb have both unique and shared effects on gene expression. Gene ontology cellular component analysis on overlapping genes revealed overrepresentation of molecules involved in synaptic composition, protein translation, and cell metabolism (**Figure 5J**, **Supplemental Table 2**); synaptic genes were also enriched in each individual mutant (**Figure S5**).

The complementary expression patterns of c-Maf and Mafb (**Figure 4**, **Figure 5A**) and their overlapping and disparate effects on subtype identity and synaptic gene expression suggest that these transcription factors impart subtype-specific influence over gene expression. To assess the relative contributions of c-Maf and Mafb across subtypes, we compared the relative log fold change for each DEG in each SGN subtype. This analysis revealed that more genes showed the biggest changes in Ia SGNs upon loss of c-Maf (59% of all c-Maf DEGs) than upon loss of Mafb (35% of all Mafb DEGs) (**Figure 5H**). The opposite was true for Mafb^cKO^ SGNs, which showed more prominent changes in gene expression in Ic SGNs (44% of all Mafb DEGs vs 16% of all c-Maf DEGs). Collectively, these data suggest that the combination of c-Maf and Mafb shapes the program of synaptic gene expression in each SGN subtype.

In addition, we found that *Mafb* expression increased in all *c-Maf* mutant SGN subtypes and *c-Maf* expression decreased in *Mafb* mutant Ia SGNs (**Figure 5K**). Therefore, it is difficult to extract the contribution of each individual factor by analyzing single mutant transcriptomes alone. Indeed, many of the genes that changed in Ia SGNs in *Mafb* mutants were c-Maf driven genes, consistent with the decrease in c-Maf expression in *Mafb* mutant Ia SGNs. The upregulation of Mafb in all *c-Maf* mutant SGNs may explain why many genes change in opposite directions. These results suggest that c-Maf and Mafb influence how much of the other factor is available and thus reinforce distinct *Maf* expression patterns across SGN subtypes.

### Emergent gene expression changes and defects in subtype diversification in cDKOs

To identify the combined effects of Maf factors on SGN gene expression, we collected SGN transcriptomes from cDKO (*bhlhe22^Cre/+^;Mafb^fl/fl^;c-Maf^fl/fl^*, N=3 mice, n=705 SGNs) and control (*Mafb^fl/fl^;c-Maf^fl/fl^*, N=3 mice, n=907 SGNs) littermates. Unsupervised graph-based clustering revealed that control SGNs formed four distinct clusters that corresponded to Type II, Ia, Ib, and Ic SGN subtypes. By contrast, cDKO neurons formed two clusters, one for Type II SGNs and one cluster for all Type I SGNs that was well-separated from control neurons (**Figure 6A**). Comparison to controls suggested that the cDKO neurons acquire a Ib/Ic identity, since most of the cDKO cells expressed Ib and Ic genes such as *Runx1* and lacked expression of Ia genes such as *Rxrg*. However, cDKO neurons did not cluster with control Ib or Ic neurons and had lower expression of *Lypd1,* a Ic marker, compared to Ic neurons in controls. There were also a few *Lypd1-*negative neurons that were slightly segregated from the other cDKO SGNs, suggesting a low degree of molecular diversification. Recent work has shown that Ia SGNs downregulate *Runx1* over development whereas the sustained expression of *Runx1* directs SGNs to a Ib or Ic subtype identity (Petitpré et al., 2022; Sanders and Kelley, 2022; Shrestha et al., 2023). Unlike c-Maf and Mafb single knockouts, cDKO SGNs did not segregate into three distinct subtypes despite sustaining expression of Runx1. This means that Runx1, cannot direct SGNs to mature Ib or Ic identities unless there is at least c-Maf or Mafb present. These results suggest that *c-Maf* and *Mafb* function together to execute SGN diversification.

**Figure 6.**
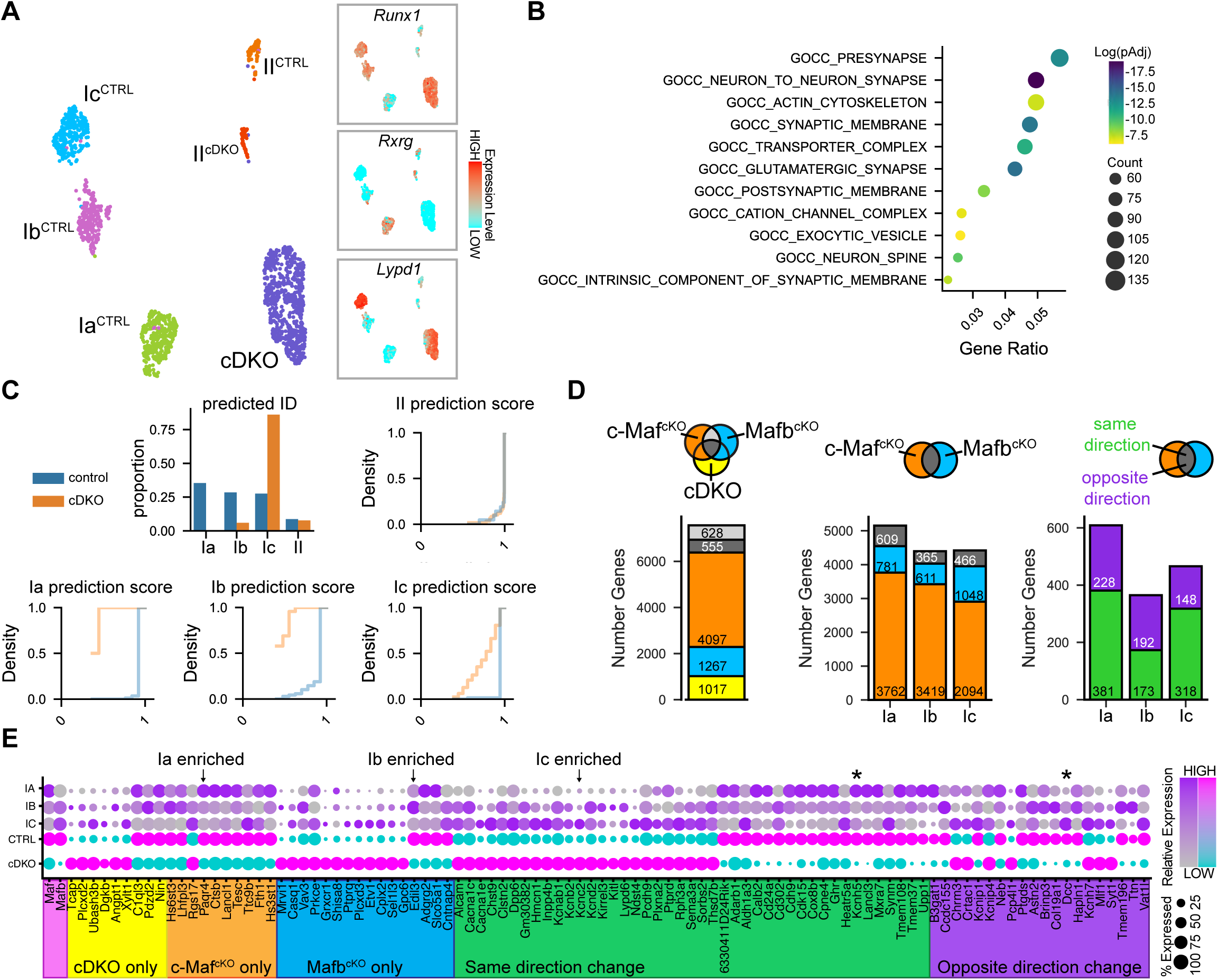
c-Maf and Mafb synergistically regulate subtype identity and synaptic gene expression but also have independent, subtype-specific effects. **(A)** UMAP plot summarizing sequencing data of double knockout (cDKO, *bhlhe22^Cre/+^;Mafb^fl/fl^;c-Maf^fl/fl^*, N=3, n=763) SGNs and controls (*Mafb^fl/fl^;c-Maf^fl/fl^*, N=3, n=974). Inset plots display expression of the Ia marker *Rxrg*, the Ib/Ic marker *Runx1*, and the Ic marker *Lypd1* in control and cDKO SGNs. (B) Cellular component gene ontology analysis of differentially expressed genes (DEG) in cDKO animals reveals strong enrichment for synapse-related genes. **(C)** The proportion of predicted SGN subtypes for cDKO cells and prediction scores compared to withheld control cells. **(D)** Meta-analysis of all differentially expressed genes (DEG) from all three scRNA-seq datasets identified both synergistic and independent effects, as indicated by whether they changed only in c-Maf^cKO^ SGNs (orange), Mafb^cKO^ SGNs (blue), or cDKO SGNs (yellow) or were jointly regulated (gray), shown for all SGNs (left) and for SGN subtypes (middle). Jointly regulated genes changed either in the same direction (green) or opposite directions (purple) in all three SGN subtypes. When considering subtype-specific effects on gene expression, cDKO SGNs were not included as they do not take on a clear identity. **(E)** Dot plot of the top 50 upregulated (magenta) and top 50 downregulated (cyan) DEGs in cDKO SGNs and their normal pattern of expression in Ia, Ib, and Ic SGN subtypes. Examples of Ia, Ib, and Ic enriched genes are indicated by arrows. Asterisks indicate an example of one gene that changes in the same direction (*Kcnh5*) and one that changes in the opposite direction (DCC). The size of the dot represents the percent of cells that had an expression level above 0. Genes are organized by whether they were changed only in the double mutants (yellow), only in one single mutant (orange and blue), in the same direction in both single mutants (green) or in opposite directions in both single mutants (purple). Color schemes in E match the colors illustrated in the Venn diagrams in D.

Double mutant SGNs showed many changes in gene expression that support a combinatorial Maf code that both reinforces SGN identity and shapes subtype-appropriate synaptic properties. Comparison of control and cDKO SGNs identified 3011 significantly differentially expressed genes (DEG) that were both up (1973 genes) and downregulated (1038 genes). Gene ontology analysis demonstrated that many DEGs were synaptic (**Figure 6B**). In fact, all top 10 cellular component GO terms were synaptic, whereas in the overlapping single knockout genes from (**Figure 5J**) only 2/10 of the top cellular component GO terms were synaptic. These results underscore the combined effect of c-Maf and Mafb on SGN synaptic gene expression.

While the cDKO cluster primarily expressed Ic genes, we wanted to test the degree and completeness of this apparent shift in identity. Again using a canonical correlation analysis, we projected cDKO cells onto each control population of SGN subtypes. Most cDKO cells were predicted to be Ic SGNs with a small proportion being assigned as Ia or Ib SGNs (**Figure 6C**). Although the Type II control and mutant clusters separated, there was no obvious change in the proportion of Type II SGNs in the cDKO, and these prediction scores were both close to 1 and overlapped obviously. Thus, gene expression changes in Type II SGNs do not seem to cause any notable shift in Type II identity. By contrast, prediction scores were low for all three Type I SGNs subtypes, but slightly higher when compared to Ic control SGNs, confirming that Type I cDKO SGNs are closest to, but not perfectly aligned with, control Ic SGNs. Consistent with a Ic identity, closer examination of the top 50 upregulated and downregulated genes (n=100 total), demonstrated that the majority (36/50) of the top upregulated genes in cDKO SGNs were normally expressed at higher levels in Ic SGNs, whereas most of the downregulated genes were enriched in Ia SGNs (30/50) (**Figure 6E**).

Comparison to all DEGs that change in any dataset (7564 genes) revealed 1017 genes whose expression changed only in the double mutants, indicating that c-Maf and Mafb can substitute for each other at many loci (**Figure 6D**, left). However, consistent with the presence of opposing phenotypic effects, many genes changed only in the c-Maf^cKO^ (4097 DEG) or Mafb^cKO^ dataset (1267 DEGs). Thus, each Maf factor also has independent functions on the maintenance of adult synapses, with a dominant role for c-Maf. For genes that changed in all three subtypes in the single knockouts, more genes were changed in Ia SGNs than Ic SGNs in the c-Maf dataset (3762 *vs.* 2094 genes) but in the Ic SGNs (1048 *vs.* 781 genes) in the Mafb dataset (**Figure 6D**, middle). DEGs common to both datasets changed in both the same and opposite directions when in the same subtype (**Figure 6D**, right). These results provide further support for both overlapping and distinct subtype-specific gene expression influence of c-Maf and Mafb.

### Dose-dependent, combinatorial gene expression by c-Maf and Mafb

The graded expression of c-Maf and Mafb across SGN subtypes (**Figure 4**) and subtype-specific influence on gene expression (**Figure 5**) indicate that these factors may modulate expression of genes in a dose-dependent manner instead of acting as binary switches. To test this idea, we characterized the response of a single DEG that changed in the same direction but to different degrees upon loss of c-Maf or Mafb. *Calb2*, which encodes a calcium binding protein that is enriched in Ia SGNs (**Figure 4,7** and Petitpré et al., 2018; Shrestha et al., 2018; Sun et al., 2018), was strongly downregulated, but still detectable in c-Maf^cKO^ SGNs, barely affected in Mafb^cKO^ SGNs, and undetectable in cDKO SGNs (**Figure 7A**). Thus, in this case, c-Maf appears to be a stronger regulator than Mafb, although both contribute. To validate these effects at the protein level, we generated mouse lines with different copy numbers of each factor (homozygotes = 2 copies, heterozygotes = 1 copy, null = 0 copies). We found that different dosages of c-Maf and Mafb resulted in different patterns of CALB2 expression, with a dominant effect for c-Maf. Reflecting transcriptomic results, comparison of c-Maf^cKO^ animals (**Figure 7B’**) to control littermates (**Figure 7B**) revealed clear downregulation of CALB2; however, several cells retained low levels of CALB2 expression. By contrast, comparison of Mafb^cKO^ animals (**Figure 7C’**) to control littermates (**Figure 7C**) did not reveal an appreciable effect on CALB2 protein expression. For each of these comparisons, knockout animals had two full copies of the other factor. Comparison of *c-Maf* heterozygous animals to littermate controls also revealed no difference in CALB2 expression (**Figure S7**). However, concomitant loss of one copy of *c-Maf* and both copies of *Mafb* revealed slightly decreased CALB2 expression (**Figure 7D,D’**). Meanwhile, loss of *c-Maf* in the context of only one copy of *Mafb* (**Figure 7D”**) revealed near complete loss of CALB2 expression. The subtle dose-dependent effect of *Mafb* on CALB2 expression can be appreciated by comparing a c-Maf^cKO^ with both copies of *Mafb*, which retains low levels of CALB2 (**Figure 7B’**), to a c-Maf^cKO^ with only one copy of *Mafb*, which contains no detectable CALB2 (**Figure 7D”**). Previous experiments with reporter mice have shown that occasionally a cell will not undergo Cre-mediated recombination, explaining the single cell that retains high expression of CALB2 in this micrograph. As expected, cDKO animals also had no detectable levels of CALB2 (**Figure 7D’’’**). Therefore, c-Maf has strong, dose-dependent influence over CALB2 expression. Meanwhile *Mafb* dosage also affects CALB2 expression but to a weaker extent that is only revealed upon loss of *c-Maf*. Remarkably, this is true even when both transcription factors are expressed in the same SGN subtype, indicating that subtype identity *per se* is not responsible for the observed differences in CALB2 expression.

**Figure 7.**
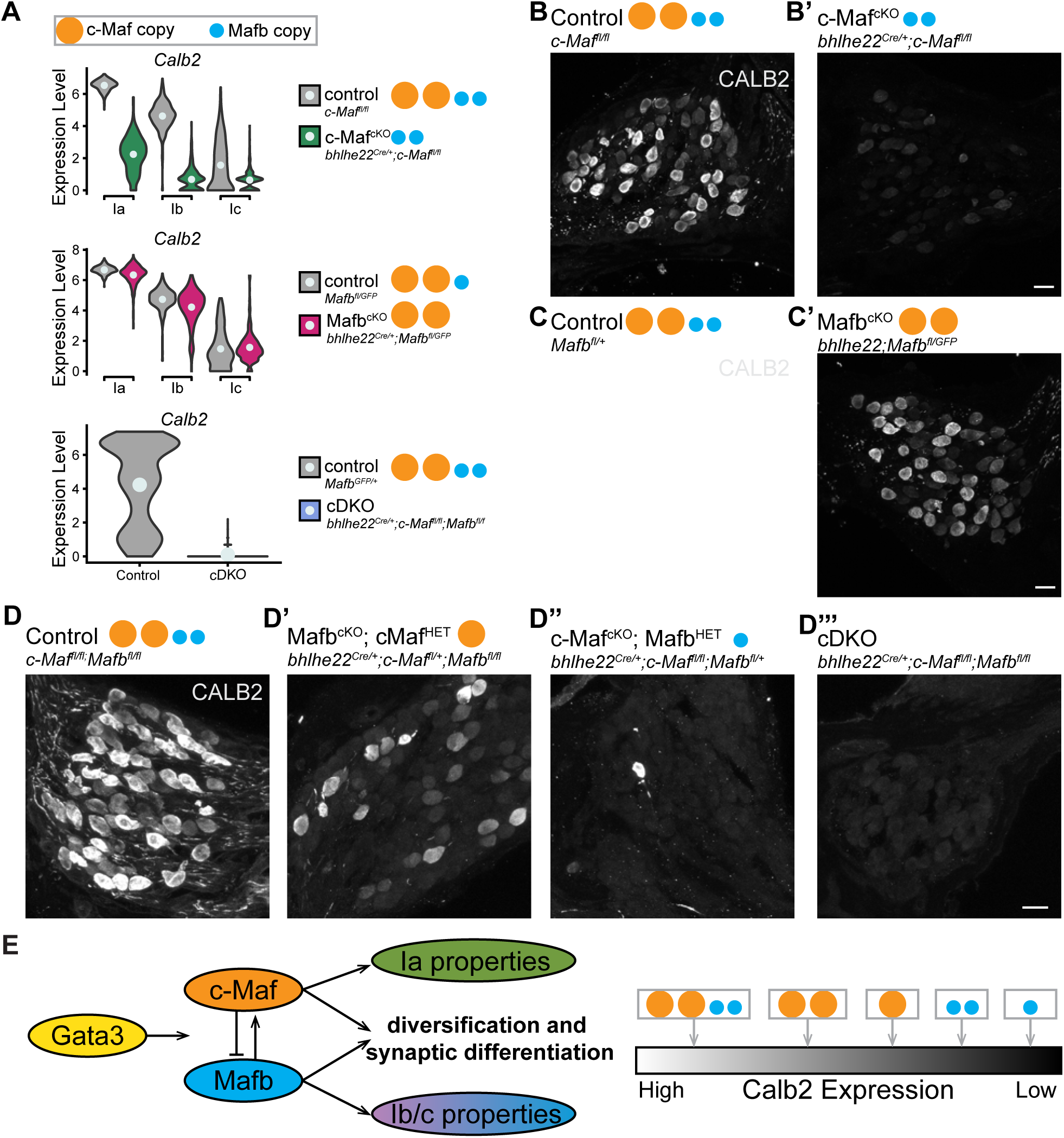
Dose-dependent, combinatorial gene expression of Calb2 by c-Maf and Mafb. **(A)** *Calb2* expression levels in c-Maf^cKO^, Mafb^cKO^ and cDKO SGNs as determined by scRNA-seq. The number of genomic copies of each TF is indicated by orange (*c-Maf*) and blue (*Mafb*) dots. **(B-D)** Immunostaining of CALB2 in sections through the cochleae of adult c-Maf^cKO^ mutants (**B’**) and littermate controls (**B**); Mafb^cKO^ mutants (**C’**) and littermate controls (**C**); and in an allelic series that includes control littermates (**D**), cKOs that retained only one copy of *c-Maf* (**D’**) or *Mafb* (**D’’**) and cDKOs (**D’’’**). CALB2 levels vary with Maf factor copy number. Scale bar= 20μm. See also Figure S7.

Together, these results support a model whereby c-Maf and Mafb combinatorially and dose-dependently coordinate subtype-specific gene expression across SGNs (**Figure 7E**). In this model, c-Maf and Mafb function redundantly to create differences among SGN subtypes and drive synaptic differentiation more generally. At the same time, c-Maf and Mafb each seem to have independent roles that allow them to act together to shape synaptic properties across SGN subtypes needed for the sense of hearing.

## Discussion

Functional diversity among neurons endows sensory systems with the capacity to encode complex sensory information. In the auditory system, spiral ganglion neurons (SGNs) exhibit functional, molecular, and synaptic differences that allow them to collectively capture the rich array of sounds encountered in the environment, even when there is background noise. Here, we show that two related transcription factors cooperate to generate synaptic and functional response properties across SGN subtypes. We find that c-Maf and Mafb establish both shared and subtype-specific features of SGN peripheral synapses that influence auditory responses. Likewise, c-Maf and Mafb regulate distinct programs of gene expression that include both shared and unique genes associated with the synapse. In parallel, c-Maf and Mafb seem to work redundantly to control expression of genes needed for both subtype diversification and synaptic differentiation, as revealed by analysis of double mutant mice. Collectively, our findings suggest that subtype-appropriate synaptic properties are shaped by the relative levels of c-Maf and Mafb, which vary across subtypes. We propose that by deploying both distinct and overlapping gene expression programs in SGN subtypes, c-Maf and Mafb combinatorially promote gene expression programs that establish and diversify SGN synaptic properties (**Figure 7E**).

Our data support the idea that a combinatorial Maf-dependent transcriptional program regulates gene expression across subtypes in a dose-dependent and partially redundant manner, thereby conferring both flexibility and resilience. We found that both c-Maf and Mafb RNA and protein are expressed in complementary patterns across SGN subtypes, with c-Maf highest in Ia SGNs and Mafb highest in Ib/Ic SGNs (**Figures 4** and **5**). Consistent with the idea that c-Maf has more influence over gene expression in Ia neurons and Mafb has more influence over gene expression in Ib/Ic SGNs, loss of either transcription factor had distinct effects on gene expression, synapses, and functional output. Further, many Maf-dependent genes are normally expressed in a subtype-specific fashion, exemplified by *Calb2,* which is expressed at high levels in cells with high c-Maf levels (**Figure 4**). Moreover, Mafb influences *Calb2* expression to a lesser degree than c-Maf, further indicating that these two Maf factors are not functionally equivalent, even in the context of the same neuronal subtype (**Figure 7**). As a result, CALB2 expression levels correlate with increasing copy numbers of Mafb and c-Maf, again with c-Maf having a dominant effect that is finely modulated by Mafb. This dose dependent regulation allows for tuning of *Calb2* expression across subtypes. The transcription factor Sox9 also exerts dose-dependent effects on gene expression in the cranial neural crest, underscoring the importance of levels differences more broadly (Naqvi et al., 2023).

At the same time, c-Maf and Mafb redundancy provides resilience against the loss of a single Maf factor. We find that many genes needed for synaptic differentiation can be regulated by either factor, providing a safeguard against a complete functional failure (**Figure 6**). For example, CALB2 protein levels are not strikingly different with the loss of a single copy of c-Maf or Mafb (**Figure 7** and **Figure S7**). Therefore, c-Maf and Mafb both fine-tune and ensure the reliable expression of genes that are differentially expressed across subtypes. The importance of combinatorial transcriptional codes is well-established, and indeed combinations of transcription factors are excellent predictors of neuronal identity (Filippopoulou et al., 2021; Hoermann et al., 2020; Kratsios et al., 2017; Sousa and Flames, 2022). To date, levels differences have not been routinely incorporated into published maps of TF expression. Our findings suggest an added layer of complexity in the production of neuronal subtypes, such that the presence of certain factors such as Runx1 and Gata3 may control key aspects of identity while more finely graded differences may be achieved by the functional diversity that exists within transcription factor families and in their expression levels, as we see with c-Maf and Mafb.

There are several mechanisms by which c-Maf and Mafb could be working together and separately. One possibility is that different levels of c-Maf and Mafb result in distinct transcriptional outcomes, due perhaps to differences in their binding affinity to target DNA due to the surrounding sequences or the presence of cofactors. Additionally, differences in their structure may alter how c-Maf and Mafb bind to genomic loci (Pogenberg et al., 2014; Rodríguez-Martínez et al., 2017; Suda et al., 2014; Yang and Cvekl, 2016). Finally, gene expression may be further shaped by which loci are accessible to either factor in each SGN subtype. More detailed analysis of chromatin structure and Maf factor binding affinities is needed to understand how a Maf code might work at the molecular level.

The effects of each Maf factor on gene expression may also be influenced by interactions with each other and other transcriptional regulators. c-Maf and Mafb can homodimerize with themselves and heterodimerize with each other and other AP-1 transcription factors (Pogenberg et al., 2014; Rodríguez-Martínez et al., 2017; Suda et al., 2014; Yang and Cvekl, 2016). This ability for homo- and heterodimerization provides a basis for combinatorial and synergistic control of gene expression (Rodríguez-Martínez et al., 2017). It has been shown that mutations that force Mafb to form only homodimers result in binding of symmetrical Maf recognition element (MARE) DNA binding sites. Meanwhile, mutant forms of Mafb that could only form heterodimers with the immediate early gene (IEG) c-Fos had entirely different DNA binding preferences (Pogenberg et al., 2014). The extent of hetero- and homodimerization among c-Maf, Mafb, and c-Fos could depend on the relative stoichiometries and binding preferences of available dimerization partners. Our discovery that c-Maf and Mafb stoichiometry varies across SGN subtypes suggests one potential mechanism for nuanced differences in the control of gene expression. Characterization of protein-protein interactions in SGNs is needed to determine which dimers are present, as well as their effects on gene expression.

The potential for interactions with IEGs such as c-Fos raises the possibility that c-Maf and Mafb act as conduits between incoming activity and synaptic refinement. Indeed, previous work has established an important role for neuronal activity in diversifying SGN subtypes (Shrestha et al., 2018; Sun et al., 2018). Neuronal activity stimulates the expression of IEGs such as c-Fos and c-Jun, which induce transcription of a cohort of genes that encode proteins that are localized to or act at synapses (Joo et al., 2016; Sheng and Greenberg, 1990). Both c-Maf and Mafb can dimerize with c-Fos, while only Mafb can dimerize with c-Jun, offering additional opportunity for activity-dependent effects within subtypes (Yang and Cvekl, 2016). Additionally, Mafb has been shown to dimerize with ATF transcription factors, a subset of which are strongly induced across SGN subtypes after noise exposure (Milon et al., 2021). Relevant activity could come either from the IHCs, which have highly heterogeneous presynaptic release sites (Meyer et al., 2009; Özçete and Moser, 2021; Payne et al., 2021) or from the olivocochlear efferents, which form more synapses on Ib and Ic SGNs than on Ia SGNs (Hua et al., 2021). Indeed, severing of olivocochlear efferents in adult mice is sufficient to alter synapse punctum size (Yin et al., 2014). Both c-Maf and Mafb are expressed in adults and could contribute to this kind of plasticity, as well as the maintenance of subtype-specific synaptic properties throughout life.

In addition to their relatively subtle effects on subtype-related synaptic properties, Maf factors act synergistically to diversify SGN subtypes and promote synaptic differentiation more generally. Using canonical correlation analysis, we confirmed that Ia, Ib, and Ic SGNs still form in single mutants, though identities were not perfectly normal (**Figure 5**). On the other hand, the double mutant SGNs formed a single cluster that were closest to a Ic identity (**Figure 6**). All of these phenotypes are fundamentally different from what occurs in *Runx1* conditional knock-out mice, where many mutant Ib and Ic SGNs take on Ia SGN molecular identities that co-cluster with control Ia SGNs (Shrestha et al., 2023). Without any Maf factors, SGNs do not diversify at all and also do not assume any recognizable identity. Thus, our work suggests that Maf factors help to execute subtype specification programs initiated by Runx1 and/or other undefined transcription factors. This phenotype fits with recent observations that developing SGNs pass through a shared Ib/Ic precursor state characterized by expression of *c-Maf* and *Mafb* (Sanders and Kelley, 2022). Although cDKO SGNs appear to retain some aspects of a Ic identity, development is not arrested in the cDKO SGNs, which still send peripheral processes towards hair cells and make post-synaptic densities that appose pre-synaptic ribbons, albeit in an aberrant manner. Thus, other as yet unidentified factors likely regulate additional gene expression programs necessary for these wiring events. Additionally, while we have focused on synaptic phenotypes, c-Maf and Mafb may also influence other subtype-specific properties, such as excitability and central projection patterns. Indeed, genes that encode channels also depend on Maf activity, though additional work is needed to determine which genes are direct targets and which might change as a result of altered synaptic signaling. These results fit with what was observed in cultured cortical interneurons where c-Maf and Mafb have antagonistic effects on synaptic properties and neuronal excitability (Pai et al., 2019) and also regulate interneuron fate (Pai et al., 2020). Altogether, the development of functional synapses with subtype-specific features seems to rely on the combined activity of Maf factors that work together to induce a general synaptic differentiation program and separately to fine-tune that program according to cell identity.

The formation of functionally and morphologically heterogeneous synapses is essential for proper circuit function. Our results show one mechanism by which synapse identity can be linked to neuronal identity. In more complex circuits, individual neurons can make many different types of synapses, making it challenging to identify relevant proteins or to understand how subtle differences in protein composition impact synaptic function. In SGNs, synaptic heterogeneity can be studied synapse by synapse and at the level of a single neuron. A deeper understanding of how c-Maf and Mafb and their target genes build synapses with different properties in SGNs will inform how synapse heterogeneity arises in other neuron types.

## Acknowledgements

Thank you to Dr. Winthrop Gillis for assistance with code, Dr. Bernardo Sabatini for access to the 10x Chromium controller, the Bauer Core Facility (Harvard University) for sequencing our samples, Dr. Chester Chia for technical assistance, and to Dr. Brikha Shrestha, Dr Michael Greenberg, and Dr. Corey Harwell for helpful feedback on the manuscript. This work was supported by DC R010009223 (to LVG), an HHMI Gilliam Fellowship (to IMB), DCF32DC019009 (to AAS), and DCF32DC012695 (to WY).

## Author contributions

IMB conceived of the project, performed experiments, analyzed results, and wrote the manuscript. LL, CMR, and AAS performed experiments and analyzed results. AY and WY initially discovered the *c-Maf* and *c-Maf;Mafb* double mutant phenotypes. LVG conceived of the project, interpreted results, and wrote the manuscript.

## Supplemental information

Figures S1-S3, S5, S7 and Table S1 and Table S2.

**Figure S1. Efferent expression and cochlear phenotypes in *Maf* mutants.** Single cell RNA sequencing reveals expression of *c-Maf* **(A)** and *Mafb* **(B) in** faciobrachial motor neurons but not in lateral olivocochlear efferents (LOC) or medial olivocochlear efferents (MOC) (Frank et al., 2023). **(D)** Wholemount immunostaining of neurofilament heavy chain (NF-H) in cochleae from control animals (NoCre, *Mafb^fl/fl^;c-Maf^fl/fl^,* N=6), *c-Maf* knockout (*c-Maf*^cKO^*, bhlhe22^Cre/+^;Mafb^fl/+^;c-Maf^fl/fl^*, N=1), *Mafb* knockout (*Mafb*^cKO^, *bhlhe22^Cre/+^;Mafb^fl/fl^;c-Maf^fl/+^*, N=1), and double knockout (cDKO, *bhlhe22^Cre/+^;Mafb^fl/fl^;c-Maf^fl/fl^,* N=7) littermates. Bottom panels provide close-up views of boxed regions showing qualitatively normal Type I SGN wiring in all genotypes and abnormal Type II SGN wiring in Mafb^cKO^ and cDKOs. Scale bar, 100μm.

**Figure S2. Synapse morphology quantification in *Maf* mutants. (A)** Unnormalized data from 8 kHz region of the cochlea. **(B)** Normalized data from 8 kHz region of the cochlea. Volume measurements were normalized by the median of the control values. **(C-E)** Unnormalized data from Figure 1B-D. **(F)** GluA2 puncta volumes in control (NoCre, *c-Maf^fl/fl^*) and c-Maf^cKO^ (*bhlhe22^Cre/+^;c-Maf^fl/fl^*) animals (t-test, p<0.001). **(G)** Median GluA2 puncta volumes in control and c-Maf^cKO^ animals. All error bars are standard error from the mean.

**Figure S3. Outer hair cell function and Peak I latency in *Maf* mutants. (A)** Mean Distortion Product Otoacoustic Emission (DPOAE) thresholds across frequencies. Standard error shown by vertical bars (all comparisons not significant, p>0.05, Kruskal-Wallis). **(B)** P1 latency across sound pressure levels for a 16 kHz stimulus. Comparisons with an asterisk were statistically significant (p<0.05, Kruskal-Wallis). Significant differences amongst groups that had a Kruskal-Wallis p-value<0.05 were followed with a pairwise post-hoc Dunn test. **(C)** P1 amplitude and **(D)** P1 latency across sound pressure levels for an 8kHz stimulus. Kruskal-Wallis p-values and pairwise post-hoc Dunn p-values for all comparisons can be found in Supplemental Table 1. All error bars are bootstrapped 68% confidence interval.

**Figure S5. Gene Ontology Analysis of c-Maf^cKO^ and Mafb^cKO^ differentially expressed genes.** Gene ontology analysis of differentially expressed genes (DEG) in cMaf **(A)** and Mafb **(B)** mutant animals.

**Figure S7. Loss of one copy of c-Maf does not affect CALB2 expression. (A)** Immunohistochemical stains of CALB2 in sections through the cochlea of adult c-Maf^HET^ animals and littermate controls. Scale bar= 20μm.

## STAR Methods

### RESOURCE AVAILABILITY

#### Lead contact

Further information and requests for resources and reagents should be directed to and will be fulfilled by the lead contact, Lisa Goodrich

#### Materials availability

This study did not generate any new unique reagents.

#### Data and code availability

Raw scRNA-seq data generated in this study, processed gene expression matrices and related metadata have been deposited at the NCBI Gene Expression Omnibus (GEO) repository with the accession numbers XXX. Custom R and python scripts are available on GitHub. Any additional information required to reanalyze the data reported in this work is available from the lead contact on request.

### EXPERIMENTAL MODEL AND SUBJECT DETAILS

Mice were handled and housed in accordance with standards and guidelines set by the Institutional Animal Care and Use Committees (IACUC) at Harvard Medical School. Animals of both sexes were used in equal proportions. No analysis was performed to determine the influence or association of sex on the results of the study. Animal genotype and age are reported where applicable, including in the Key Resources table.

#### Animal Models

*Bhlhe22^Cre/+^* (*Bhlhe22^Cre/+^*, MGI:4440745), *c-Maf^fl^* (MGI:5316775), *Mafb^fl^* (MGI:5581666) and Rosa26-LSL-tdTomato (*Ai14*; Jax strain 007914) are all previously described. *NetrinG1^Cre/+^* mice were kindly provided by Dr. Fan Wang (M.I.T) (Bolding et al., 2020). Animals were maintained on a mixed background. Animal work was conducted in compliance with protocols approved by the Institutional Animal Care and Use Committee at Harvard Medical School.

#### Immunostaining

Animals were anesthetized via isoflurane exposure in an open-drop chamber and subsequently perfused with cold 4% paraformaldehyde (PFA) in 1X PBS. Both temporal bones were dissected and cold 4% PFA was perfused through the oval window with a syringe. For wholemount immunohistochemistry, cochleae were drop-fixed in 4% PFA at room temperature and transferred to 10% EDTA overnight at 4C. Cochlea were then micro-dissected and transferred to blocking solution (16%v/v normal donkey serum, 3%v/v Triton-X in 1xPBS) overnight at 4°C. Cochlear turns were stained overnight with anti-Calb2 (1:1000, Swant CG1), anti-CTBP2 (1:500, BD Transduction Laboratories, Clone 16) and anti-GluA2 (1:500, EMB Millipore MAB 397) and subsequently with corresponding secondaries at 37°C overnight. Rinses after primary and secondary antibody incubation were done using 1% PBST for 10 minutes at room temperature. Cochleae used for cryosectioning and staining were drop-fixed overnight at 4°C in 4% PFA in 1x PBS and then transferred to 120mM EDTA for three nights. Cochleae were then immersed in a sucrose gradient from 10% to 30% at 4°C prior to embedding. 18 μm cochlear sections were washed 1x for 5 minutes in 1x PBS and then 2x for 5 minutes in 0.25% Triton X-100 in 1x PBS. Sections were blocked with 5%v/v normal donkey serum and 0.3% Triton X-100 in 1x PBS. Sections were stained with a compatible combination of: rabbit anti-MAFB (1:250, Novus Biologicals, NBP1-81342), rabbit anti c-Maf (1:250, Bethyl Labs A700-045), goat anti CALB2 (1:500, Swant CG1) for four hours at room temperature. After rinsing with 1x PBS, sections stained with appropriate secondary antibodies at 4°C overnight. Sections were then rinsed with 1x PBS for 10 minutes. The second wash contained DAPI at 1 mg/uL.

#### Image acquisition

All tissues were imaged using a Leica SP8 point-scanning confocal microscope with HyD and photomultiplier tube (PMT) detectors. For wholemount cochleae, frequency maps were generated by taking 10X stacks of the cochlear turns and using the Measure_Line ImageJ plugin available through the Histology Core at Mass Eye and Ear (Boston, MA). Synaptic puncta were then captured using HyD detectors while hair cells were captured using PMT detectors using a 63x oil-immersion objective (voxel size= 0.901 x 0.901 x 0.299um^3^). Special attention was taken not to oversaturate any pixels in channels that were to be used for intensity-based morphometric quantification.

#### Auditory response testing

Mice were anesthetized with an intraperitoneal injection of ketamine (100 mg/kg) and xylaxine (10 mg/kg). Meloxicam (1 mg/kg) was administered intraperitoneally for analgesia. Animals were placed on a 37°C heating pad (ATC1000, World Precision Instruments) and additional ketamine (30-40 mg/kg) was administered as needed to maintain the anesthetic plane throughout the procedure. ABRs and DPOAEs were measured using a custom acoustic system (Eaton-Peabody Laboratories, Massachusetts Eye and Ear) in an electrically shielded and sound attenuating chamber. All recordings were performed with the researcher blinded to genotype.

Auditory brainstem responses (ABRs) were recorded from three subcutaneous needle electrodes: a recording electrode caudal to the pinna, a reference electrode at the vertex, and a ground electrode by the tail. ABR stimuli were presented from 20 to 90 dB SPL in 5 dB steps at 8, 16, 32, and 45 kHz. Each stimulus was presented as a 5-ms tone-pip at a rate of 31/s, with a 0.5 ms rise-fall time and alternating polarities. Responses were amplified 10,000x, filtered with a 0.3–3 kHz passband (P511, Grass), and averaged 512 times. Recordings with peak to peak amplitudes exceeding 15 μV were rejected as artifacts.

Distortion product otoacoustic emissions (DPOAEs) were recorded from a probe-tube microphone aligned above the ear canal. In-ear calibrations were performed using the Cochlear Function Test Suite (v 2.36, Massachusetts Eye and Ear) prior to reach recording. DPOAE stimuli were presented at primary tones f1 and f2, where f2 varied from 5.6 to 32 kHz in half-octave steps. Primary tones were presented at frequency ratios of f2/f1 = 1.2 and level differences of L1 = L2 + 10. Levels of primary tone f2 were incremented in 10 dB steps from 0 to 70 dB SPL.

### Single-cell RNA sequencing

#### Library Preparation

Temporal bones were extracted, and the spiral ganglion was microdissected from the cochlea in cold Leibovitz L-15 buffer. Cells were then treated with collagenase type IV followed by papain for 25 minutes each at 37°C. The cells were passed through ovomucoid as recommended for the Papain Dissociation System and then passed through a 40 µm cell strainer. Dissociated cells were resuspended in cold EBSS. Cell concentration was estimated using a hemocytometer. Cells were then loaded into a single cell chip from 10x Genomics following manufacturer’s recommendations. Datasets were processed with the Chromium single-cell 3’ library and gel bead kit v2.0. cDNA libraries were generated according to the manufacturer’s directions. The final libraries were sequenced on an Illumina NovaSeq SP.

## QUANTIFICATION AND STATISTICAL ANALYSIS

All statistical analysis was done in Python. Normality of each distribution was tested using the Shapiro-Wilk test. If both groups showed a normal distribution, a parametric t-Test or ANOVA was used. Otherwise, the non-parametric Mann-Whitney Rank Sum or Kruskal-Wallis Test was applied. Statistical analysis for differentially expressed genes was done in R. Differential gene expression was tested using a Wilcoxon rank sum test with Bonferroni post-hoc correction. Gene ontology categories with a Bonferroni-adjusted P<0.001 were considered significant.

### Analysis of ABRs

ABR thresholds, amplitudes, and latencies were analyzed using ABR Peak Analysis software (v.1.1.1.9; Massachusetts Eye and Ear). ABR threshold was defined as the lowest stimulus level at which wave 1 could be identified by visual inspection.

The distortion products at 2f1-f2 were temporally and spectrally averaged. Iso-response contours were generated by the Cochlear Function Test Suite software for various criterion response amplitudes. DPOAE threshold was defined as the f1 level required to produce a DPOAE of 5 dB SPL.

### Image analysis

Pre- and post-synaptic puncta were reconstructed semi-automatically using the “Surfaces” function in Imaris. A local contrast-background subtraction algorithm was applied to all z-stacks for thresholding. All reconstructions were created and reviewed with the researcher blinded to genotype. Inaccuracies in automatic segmentation were corrected manually. Reconstructions located outside the volume of hair cells were excluded from analysis. Image-based coordinates of reconstructed puncta were transformed into hair cell-centric coordinates using the “Reference Frames” function in Imaris. Each hair cell-centric coordinate system was defined with the following three planes: an XZ plane bisecting the hair cell through its plane of symmetry, a YZ plane bisecting the nucleus through the plane parallel to the tilt of the hair cell, and an XY plane tangential to the basolateral pole of the hair cell. Metrics regarding the size and localization of reconstructed pre- and postsynaptic puncta were exported into Excel for further analysis. For quantification of c-MAF and MAFB stains, 3D reconstructions were made for each SGN cell body in Imaris similar as above using the HuD channel that marked all SGN cell bodies. Metrics regarding fluorescence intensity were exported into Excel for further analysis.

### Bioinformatics analysis

#### Alignment

Raw reads were converted to fastq files using the Cellranger pipeline from 10X Genomics v 3.0.1. Reads were then aligned to the mouse reference genome (mm10) in a Linux-based high performance computing cluster at Harvard Medical School.

#### Normalization

The aligned data was imported into R and analyzed for statistical analysis and graphical representation. The library was normalized by fitting the gene counts to a regularized binomial regression function implemented by the scTransform package for Seurat using all default settings and regression on percent mitochondrial reads.

#### Clustering and Subclustering

Clustering was performed using the Seurat FindClusters command using 30 principle components. SGN clusters were identified and subsetted out by the expression of neuronal genes such as Tubb3 and Nefh. Utilizing the RNA assay, subsetted SGNs were normalized and subclustered using similar parameters as above. Subtypes of SGNs were identified by the expression of subtype specific markers previously identified (*Calb2*, *Lypd1,* and *Runx1*) (Petitpré et al., 2018; Shrestha et al., 2018; Sun et al., 2018). Differential expression was tested using the FindAllMarkers command on the SCT assay.

#### Differential Gene Expression and Gene Ontology Analysis

Differential expression was tested using the FindAllMarkers command on the SCT assay and probing for significantly regulated genes between control and mutant SGNs from each genotype. The differential expression of cDKO genes was analyzed by subtype in each single cKO dataset. Genes were considered differentially expressed in mutant SGNs if they were significantly changed (P<0.05) at a Log2Fold change greater than 0.25. Differentially expressed genes were subjected to Gene Ontology term enrichment analysis using Fisher’s exact test and ranking the top 10 cellular component GO terms.

#### Cell Identity Projection

Knockout datasets were projected onto control datasets as described for previously for dataset integration (Stuart et al., 2019). Briefly, canonical correlation analysis was used to assign the highest correlated identity to each knockout cell. Data were not integrated for UMAP visualization, as this assumes that the cells are the same identity and occludes shifts in subtype identity. Prediction scores for each assigned identity were extracted. As a comparison, in a separate analysis, 25% of the control data were withheld and projected onto the remaining 75% to demonstrate that cells can be appropriately assigned with this method.

## References

1. Appler, J.M., Lu, C.C., Druckenbrod, N.R., Yu, W.-M., Koundakjian, E.J., Goodrich, L.V., 2013. Gata3 Is a Critical Regulator of Cochlear Wiring. Journal of Neuroscience 33, 3679– 3691. 10.1523/JNEUROSCI.4703-12.2013

2. Bolding, K.A., Nagappan, S., Han, B.-X., Wang, F., Franks, K.M., 2020. Recurrent circuitry is required to stabilize piriform cortex odor representations across brain states. eLife 9, e53125. 10.7554/eLife.53125

3. Coate, T.M., Scott, M.K., Gurjar, M., 2019. Current concepts in cochlear ribbon synapse formation. Synapse 73, e22087. 10.1002/syn.22087

4. Druckenbrod, N.R., Goodrich, L.V., 2015. Sequential Retraction Segregates SGN Processes during Target Selection in the Cochlea. J. Neurosci. 35, 16221–16235. 10.1523/JNEUROSCI.2236-15.2015

5. Filippopoulou, K., Couillault, C., Bertrand, V., 2021. Multiple neural bHLHs ensure the precision of a neuronal specification event in *Caenorhabditis elegans*. Biology Open 10, bio058976. 10.1242/bio.058976

6. Frank, M.M., Sitko, A.A., Suthakar, K., Torres Cadenas, L., Hunt, M., Yuk, M.C., Weisz, C.J., Goodrich, L.V., 2023. Experience-dependent flexibility in a molecularly diverse central-to-peripheral auditory feedback system. eLife 12, e83855. 10.7554/eLife.83855

7. Goodrich, L.V., 2016. Early Development of the Spiral Ganglion, in: Dabdoub, A., Fritzsch, B., Popper, A.N., Fay, R.R. (Eds.), The Primary Auditory Neurons of the Mammalian Cochlea, Springer Handbook of Auditory Research. Springer New York, New York, NY, pp. 11–48. 10.1007/978-1-4939-3031-9_2

8. Hoermann, N., Schilling, T., Haji Ali, A., Serbe, E., Mayer, C., Borst, A., Pujol-Martí, J., 2020. A combinatorial code of transcription factors specifies subtypes of visual motion-sensing neurons in *Drosophila*. Development dev.186296. 10.1242/dev.186296

9. Hu, N., Rutherford, M.A., Green, S.H., 2020. Protection of cochlear synapses from noise-induced excitotoxic trauma by blockade of Ca ^2+^ -permeable AMPA receptors. Proc. Natl. Acad. Sci. U.S.A. 117, 3828–3838. 10.1073/pnas.1914247117

10. Hua, Y., Ding, X., Wang, H., Wang, F., Lu, Y., Neef, J., Gao, Y., Moser, T., Wu, H., 2021. Electron Microscopic Reconstruction of Neural Circuitry in the Cochlea. Cell Reports 34, 108551. 10.1016/j.celrep.2020.108551

11. Huang, L.-C., Barclay, M., Lee, K., Peter, S., Housley, G.D., Thorne, P.R., Montgomery, J.M., 2012. Synaptic profiles during neurite extension, refinement and retraction in the developing cochlea. Neural Dev 7, 38. 10.1186/1749-8104-7-38

12. Joo, J.-Y., Schaukowitch, K., Farbiak, L., Kilaru, G., Kim, T.-K., 2016. Stimulus-specific combinatorial functionality of neuronal c-fos enhancers. Nat Neurosci 19, 75–83. 10.1038/nn.4170

13. Kane, K.L., Longo-Guess, C.M., Gagnon, L.H., Ding, D., Salvi, R.J., Johnson, K.R., 2012. Genetic background effects on age-related hearing loss associated with Cdh23 variants in mice. Hearing Research 283, 80–88. 10.1016/j.heares.2011.11.007

14. Kiang, N., 1965. Discharge Patterns of Single Fibres in the Cat’s Auditory Nerve, Massachusetts Institute of Technology. Research Laboratory of Electronics. Special technical report no. 13. Cambridge, Mass., M.I.T. Press.

15. Kratsios, P., Kerk, S.Y., Catela, C., Liang, J., Vidal, B., Bayer, E.A., Feng, W., De La Cruz, E.D., Croci, L., Consalez, G.G., Mizumoto, K., Hobert, O., 2017. An intersectional gene regulatory strategy defines subclass diversity of C. elegans motor neurons. eLife 6, e25751. 10.7554/eLife.25751

16. Kreeger, L.J., Honnuraiah, S., Maeker, S., Shea, S., Fishell, G., Goodrich, L.V., 2024. An Anatomical and Physiological Basis for Coincidence Detection Across Time Scales in the Auditory System (preprint). Neuroscience. 10.1101/2024.02.29.582808

17. Liberman, L.D., Liberman, M.C., 2016. Postnatal maturation of auditory-nerve heterogeneity, as seen in spatial gradients of synapse morphology in the inner hair cell area. Hearing Research 339, 12–22. 10.1016/j.heares.2016.06.002

18. Liberman, L.D., Wang, H., Liberman, M.C., 2011. Opposing Gradients of Ribbon Size and AMPA Receptor Expression Underlie Sensitivity Differences among Cochlear-Nerve/Hair-Cell Synapses. Journal of Neuroscience 31, 801–808. 10.1523/JNEUROSCI.3389-10.2011

19. Liberman, M.C., 1982. Single-Neuron Labeling in the Cat Auditory Nerve. Science 216, 1239– 1241. 10.1126/science.7079757

20. Melcher, J.R., Kiang, N.Y.S., 1996. Generators of the brainstem auditory evoked potential in cat III: identified cell populations. Hearing Research 93, 52–71. 10.1016/0378-5955(95)00200-6

21. Meyer, A.C., Frank, T., Khimich, D., Hoch, G., Riedel, D., Chapochnikov, N.M., Yarin, Y.M., Harke, B., Hell, S.W., Egner, A., Moser, T., 2009. Tuning of synapse number, structure and function in the cochlea. Nat Neurosci 12, 444–453. 10.1038/nn.2293

22. Michanski, S., Smaluch, K., Steyer, A.M., Chakrabarti, R., Setz, C., Oestreicher, D., Fischer, C., Möbius, W., Moser, T., Vogl, C., Wichmann, C., 2019. Mapping developmental maturation of inner hair cell ribbon synapses in the apical mouse cochlea. Proc. Natl. Acad. Sci. U.S.A. 116, 6415–6424. 10.1073/pnas.1812029116

23. Milon, B., Shulman, E.D., So, K.S., Cederroth, C.R., Lipford, E.L., Sperber, M., Sellon, J.B., Sarlus, H., Pregernig, G., Shuster, B., Song, Y., Mitra, S., Orvis, J., Margulies, Z., Ogawa, Y., Shults, C., Depireux, D.A., Palermo, A.T., Canlon, B., Burns, J., Elkon, R., Hertzano, R., 2021. A cell-type-specific atlas of the inner ear transcriptional response to acoustic trauma. Cell Reports 36, 109758. 10.1016/j.celrep.2021.109758

24. Moser, T., Karagulyan, N., Neef, J., Jaime Tobón, L.M., 2023. Diversity matters — extending sound intensity coding by inner hair cells via heterogeneous synapses. The EMBO Journal 42, e114587. 10.15252/embj.2023114587

25. Naqvi, S., Kim, S., Hoskens, H., Matthews, H.S., Spritz, R.A., Klein, O.D., Hallgrímsson, B., Swigut, T., Claes, P., Pritchard, J.K., Wysocka, J., 2023. Precise modulation of transcription factor levels identifies features underlying dosage sensitivity. Nat Genet 55, 841–851. 10.1038/s41588-023-01366-2

26. Özçete, Ö.D., Moser, T., 2021. A sensory cell diversifies its output by varying Ca ^2+^ influx-release coupling among active zones. The EMBO Journal 40. 10.15252/embj.2020106010

27. Pai, E.L.-L., Chen, Jin, Fazel Darbandi, S., Cho, F.S., Chen, Jiapei, Lindtner, S., Chu, J.S., Paz, J.T., Vogt, D., Paredes, M.F., Rubenstein, J.L., 2020. Maf and Mafb control mouse pallial interneuron fate and maturation through neuropsychiatric disease gene regulation. eLife 9, e54903. 10.7554/eLife.54903

28. Pai, E.L.-L., Vogt, D., Clemente-Perez, A., McKinsey, G.L., Cho, F.S., Hu, J.S., Wimer, M., Paul, A., Fazel Darbandi, S., Pla, R., Nowakowski, T.J., Goodrich, L.V., Paz, J.T., Rubenstein, J.L.R., 2019. Mafb and c-Maf Have Prenatal Compensatory and Postnatal Antagonistic Roles in Cortical Interneuron Fate and Function. Cell Reports 26, 1157–1173.e5. 10.1016/j.celrep.2019.01.031

29. Pata, I., Studer, M., Doorninck, J.H.V., Briscoe, J., Kuuse, S., Engel, J.D., Grosveld, F., Karis, A., 1999. The transcription factor GATA3 is a downstream effector of *Hoxb1* specification in rhombomere 4. Development 126, 5523–5531. 10.1242/dev.126.23.5523

30. Payne, S.A., Joens, M.S., Chung, H., Skigen, N., Frank, A., Gattani, S., Vaughn, K., Schwed, A., Nester, M., Bhattacharyya, A., Iyer, G., Davis, B., Carlquist, J., Patel, H., Fitzpatrick, J.A.J., Rutherford, M.A., 2021. Maturation of Heterogeneity in Afferent Synapse Ultrastructure in the Mouse Cochlea. Front. Synaptic Neurosci. 13, 678575. 10.3389/fnsyn.2021.678575

31. Perkins, R.E., Morest, D.K., 1975. A study of cochlear innervation patterns in cats and rats with the Golgi method and Nomarski optics. J. Comp. Neurol. 163, 129–158. 10.1002/cne.901630202

32. Petitpré, C., Faure, L., Uhl, P., Fontanet, P., Filova, I., Pavlinkova, G., Adameyko, I., Hadjab, S., Lallemend, F., 2022. Single-cell RNA-sequencing analysis of the developing mouse inner ear identifies molecular logic of auditory neuron diversification. Nat Commun 13, 3878. 10.1038/s41467-022-31580-1

33. Petitpré, C., Wu, H., Sharma, A., Tokarska, A., Fontanet, P., Wang, Y., Helmbacher, F., Yackle, K., Silberberg, G., Hadjab, S., Lallemend, F., 2018. Neuronal heterogeneity and stereotyped connectivity in the auditory afferent system. Nat Commun 9, 3691. 10.1038/s41467-018-06033-3

34. Pogenberg, V., Consani Textor, L., Vanhille, L., Holton, S.J., Sieweke, M.H., Wilmanns, M., 2014. Design of a bZip Transcription Factor with Homo/Heterodimer-Induced DNA-Binding Preference. Structure 22, 466–477. 10.1016/j.str.2013.12.017

35. Rodríguez-Martínez, J.A., Reinke, A.W., Bhimsaria, D., Keating, A.E., Ansari, A.Z., 2017. Combinatorial bZIP dimers display complex DNA-binding specificity landscapes. eLife 6, e19272. 10.7554/eLife.19272

36. Ross, S.E., Mardinly, A.R., McCord, A.E., Zurawski, J., Cohen, S., Jung, C., Hu, L., Mok, S.I., Shah, A., Savner, E.M., Tolias, C., Corfas, R., Chen, S., Inquimbert, P., Xu, Y., McInnes, R.R., Rice, F.L., Corfas, G., Ma, Q., Woolf, C.J., Greenberg, M.E., 2010. Loss of inhibitory interneurons in the dorsal spinal cord and elevated itch in Bhlhb5 mutant mice. Neuron 65, 886–898. 10.1016/j.neuron.2010.02.025

37. Ryugo, D.K., 1992. The Auditory Nerve: Peripheral Innervation, Cell Body Morphology, and Central Projections, in: Webster, D.B., Popper, A.N., Fay, R.R. (Eds.), The Mammalian Auditory Pathway: Neuroanatomy, Springer Handbook of Auditory Research. Springer New York, New York, NY, pp. 23–65. 10.1007/978-1-4612-4416-5_2

38. Sanders, T.R., Kelley, M.W., 2022. Specification of neuronal subtypes in the spiral ganglion begins prior to birth in the mouse. Proc Natl Acad Sci U S A 119, e2203935119. 10.1073/pnas.2203935119

39. Sendin, G., Bulankina, A.V., Riedel, D., Moser, T., 2007. Maturation of Ribbon Synapses in Hair Cells Is Driven by Thyroid Hormone. Journal of Neuroscience 27, 3163–3173. 10.1523/JNEUROSCI.3974-06.2007

40. Sheng, M., Greenberg, M.E., 1990. The regulation and function of c-fos and other immediate early genes in the nervous system. Neuron 4, 477–485. 10.1016/0896-6273(90)90106-P

41. Shrestha, B.R., Chia, C., Wu, L., Kujawa, S.G., Liberman, M.C., Goodrich, L.V., 2018. Sensory Neuron Diversity in the Inner Ear Is Shaped by Activity. Cell 174, 1229–1246.e17. 10.1016/j.cell.2018.07.007

42. Shrestha, B.R., Goodrich, L.V., 2019. Wiring the Cochlea for Sound Perception, in: Kandler, K. (Ed.), The Oxford Handbook of the Auditory Brainstem. Oxford University Press, pp. xxx–36. 10.1093/oxfordhb/9780190849061.013.1

43. Shrestha, B.R., Wu, L., Goodrich, L.V., 2023. Runx1 controls auditory sensory neuron diversity in mice. Developmental Cell 58, 306–319.e5. 10.1016/j.devcel.2023.01.008

44. Siebald, C., Vincent, P.F.Y., Bottom, R.T., Sun, S., Reijntjes, D.O.J., Manca, M., Glowatzki, E., Müller, U., 2023. Molecular signatures define subtypes of auditory afferents with distinct peripheral projection patterns and physiological properties. Proc Natl Acad Sci U S A 120, e2217033120. 10.1073/pnas.2217033120

45. Sousa, E., Flames, N., 2022. Transcriptional regulation of neuronal identity. Eur J of Neuroscience 55, 645–660. 10.1111/ejn.15551

46. Spoendlin, H., 1985. Anatomy of Cochlear Innervation. American Journal of Otolaryngology 6, 453–467. 10.1016/S0196-0709(85)80026-0

47. Stuart, T., Butler, A., Hoffman, P., Hafemeister, C., Papalexi, E., Mauck, W.M., Hao, Y., Stoeckius, M., Smibert, P., Satija, R., 2019. Comprehensive Integration of Single-Cell Data. Cell 177, 1888–1902.e21. 10.1016/j.cell.2019.05.031

48. Suda, N., Itoh, T., Nakato, R., Shirakawa, D., Bando, M., Katou, Y., Kataoka, K., Shirahige, K., Tickle, C., Tanaka, M., 2014. Dimeric combinations of MafB, cFos and cJun control the apoptosis-survival balance in limb morphogenesis. Development 141, 2885–2894. 10.1242/dev.099150

49. Sun, S., Babola, T., Pregernig, G., So, K.S., Nguyen, M., Su, S.-S.M., Palermo, A.T., Bergles, D.E., Burns, J.C., Müller, U., 2018. Hair Cell Mechanotransduction Regulates Spontaneous Activity and Spiral Ganglion Subtype Specification in the Auditory System. Cell 174, 1247–1263.e15. 10.1016/j.cell.2018.07.008

50. Wende, H., Lechner, S.G., Cheret, C., Bourane, S., Kolanczyk, M.E., Pattyn, A., Reuter, K., Munier, F.L., Carroll, P., Lewin, G.R., Birchmeier, C., 2012. The Transcription Factor c-Maf Controls Touch Receptor Development and Function. Science 335, 1373–1376. 10.1126/science.1214314

51. Yang, Y., Cvekl, A., 2016. Large Maf Transcription Factors: Cousins of AP-1 Proteins and Important Regulators of Cellular Differentiation. EJBM 23, 2. 10.23861/EJBM20072347

52. Yin, Y., Liberman, L.D., Maison, S.F., Liberman, M.C., 2014. Olivocochlear Innervation Maintains the Normal Modiolar-Pillar and Habenular-Cuticular Gradients in Cochlear Synaptic Morphology. JARO 15, 571–583. 10.1007/s10162-014-0462-z

53. Yu, W.-M., Appler, J.M., Kim, Y.-H., Nishitani, A.M., Holt, J.R., Goodrich, L.V., 2013. A Gata3– Mafb transcriptional network directs post-synaptic differentiation in synapses specialized for hearing. eLife 2, e01341. 10.7554/eLife.01341

54. Yu, W.-M., Goodrich, L.V., 2014. Morphological and physiological development of auditory synapses. Hearing Research 311, 3–16. 10.1016/j.heares.2014.01.007

55. Zhang, K.D., Coate, T.M., 2017. Recent advances in the development and function of type II spiral ganglion neurons in the mammalian inner ear. Seminars in Cell & Developmental Biology 65, 80–87. 10.1016/j.semcdb.2016.09.017

