## Supplementary figures and images for "Combinatorial transcriptional regulation establishes subtype-appropriate synaptic properties in auditory neurons"

### Supplemental Figure 1

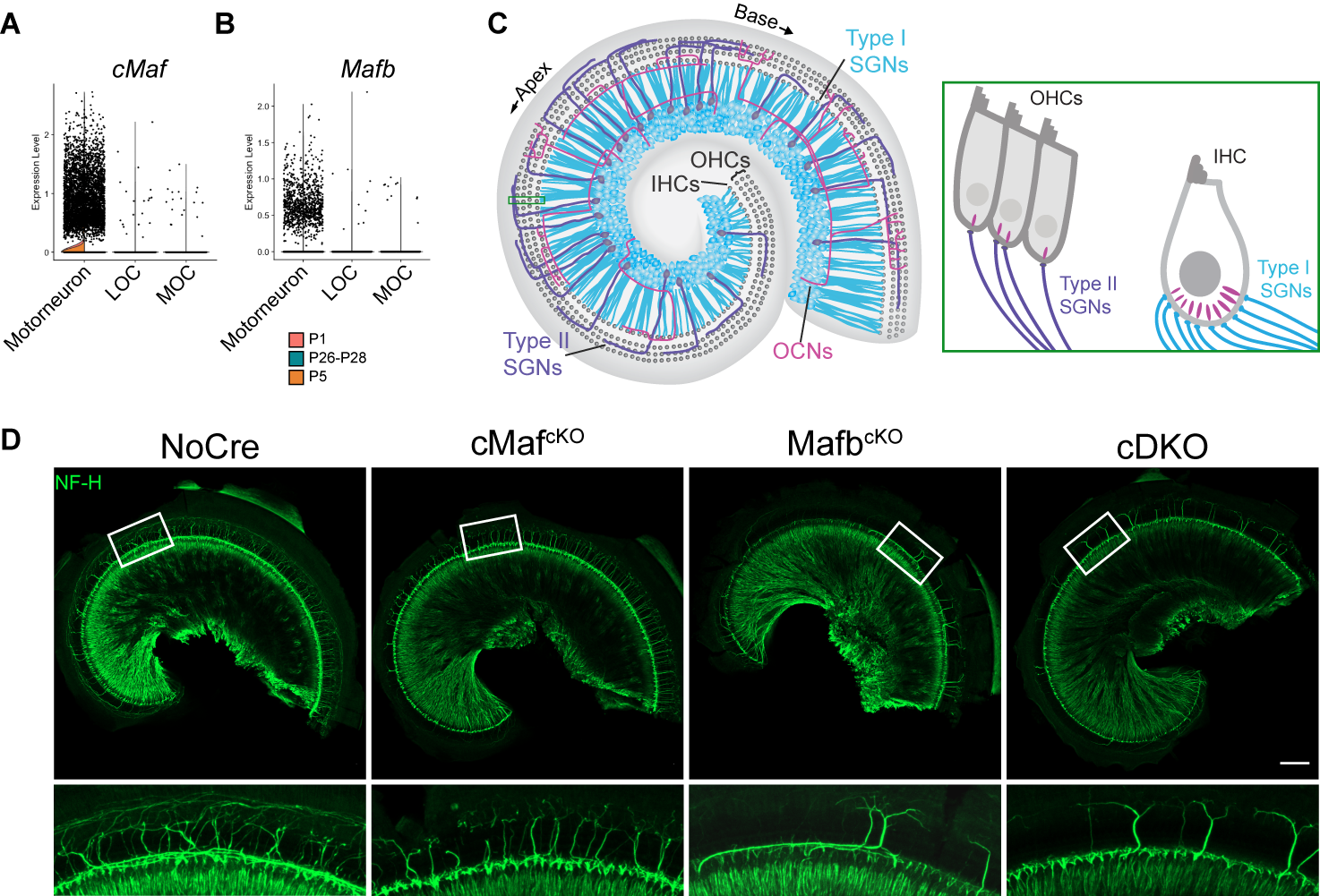

### Supplemental Figure 2

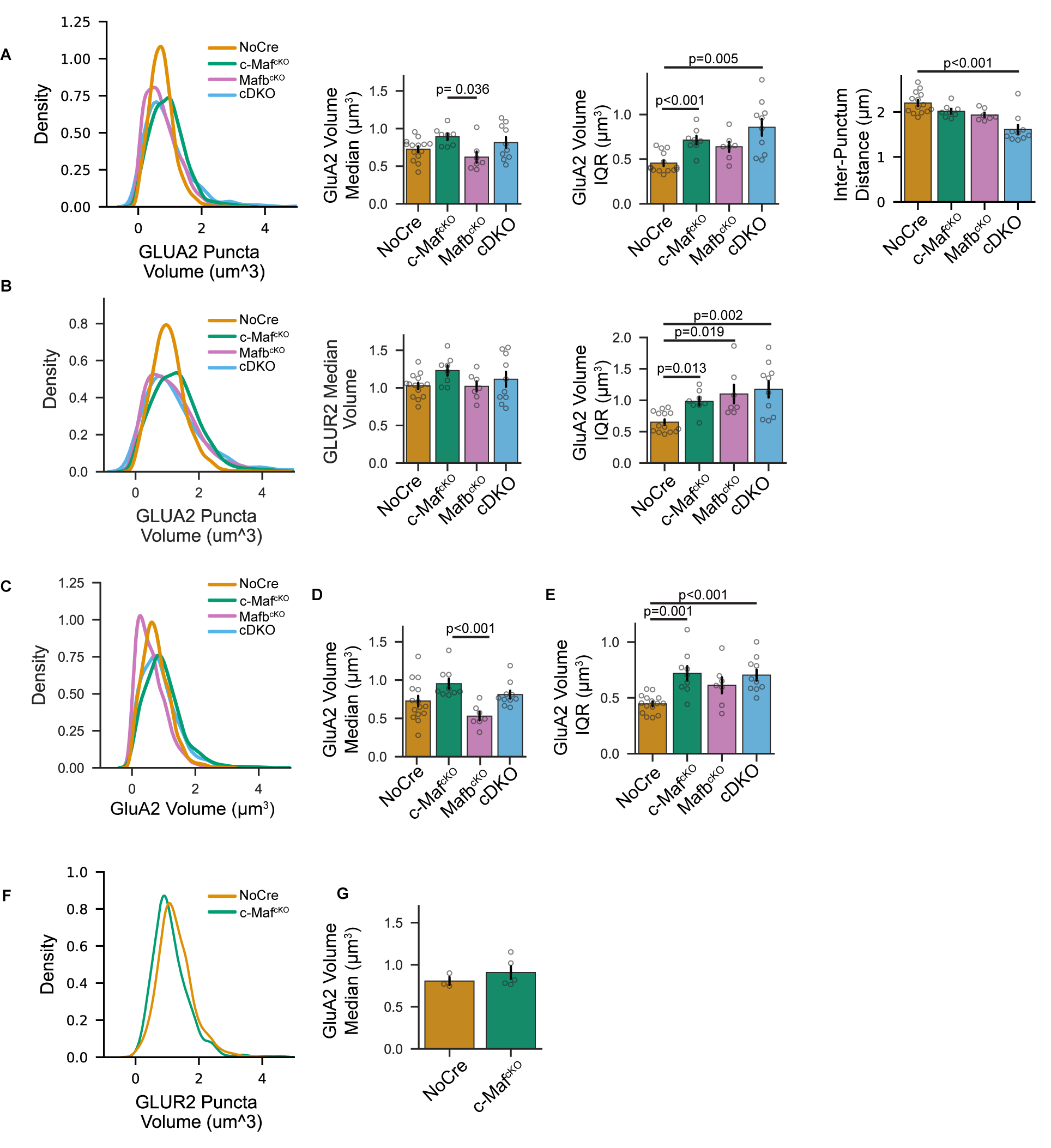

### Supplemental Figure 3

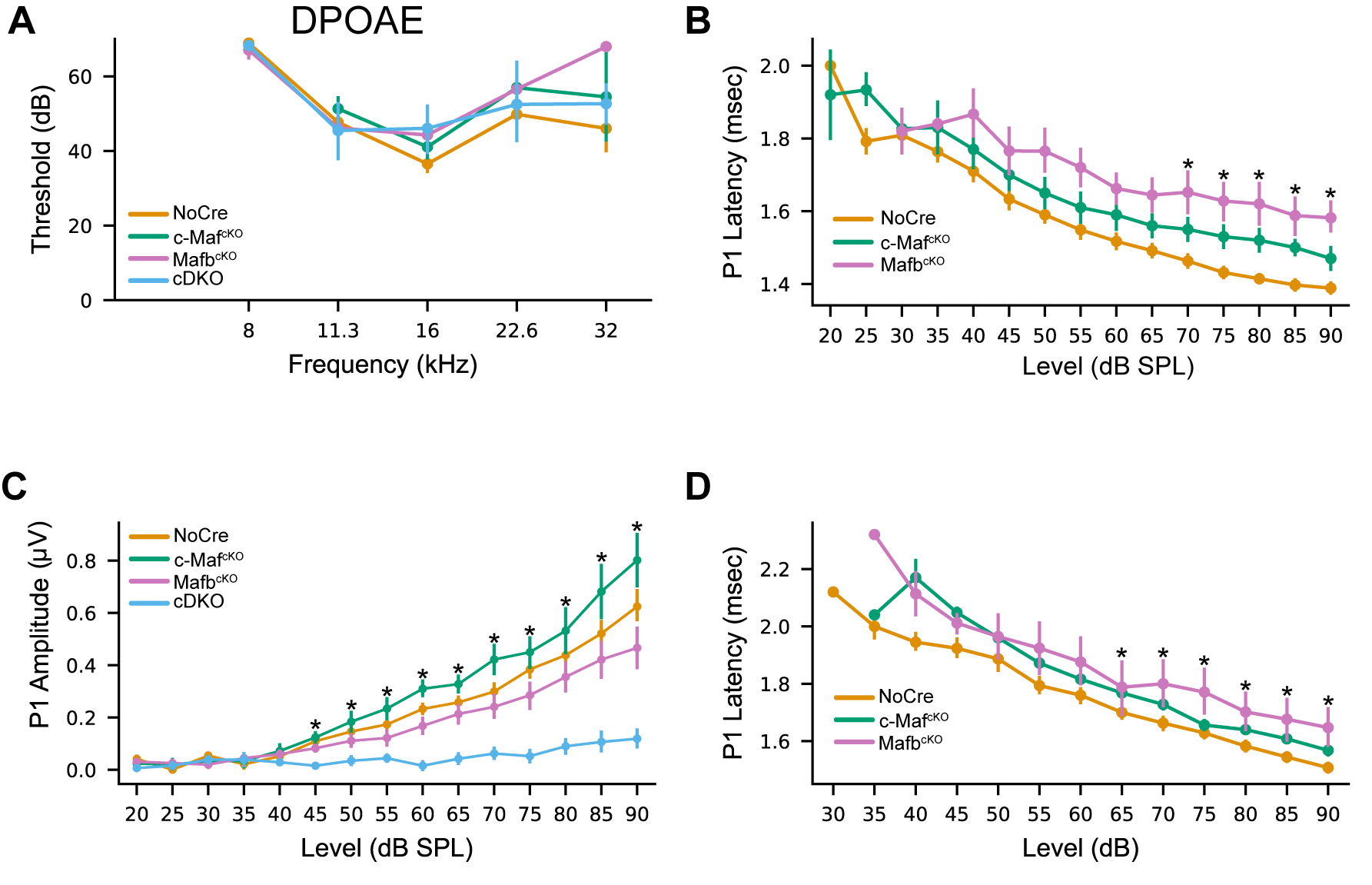

### Supplemental Figure 5

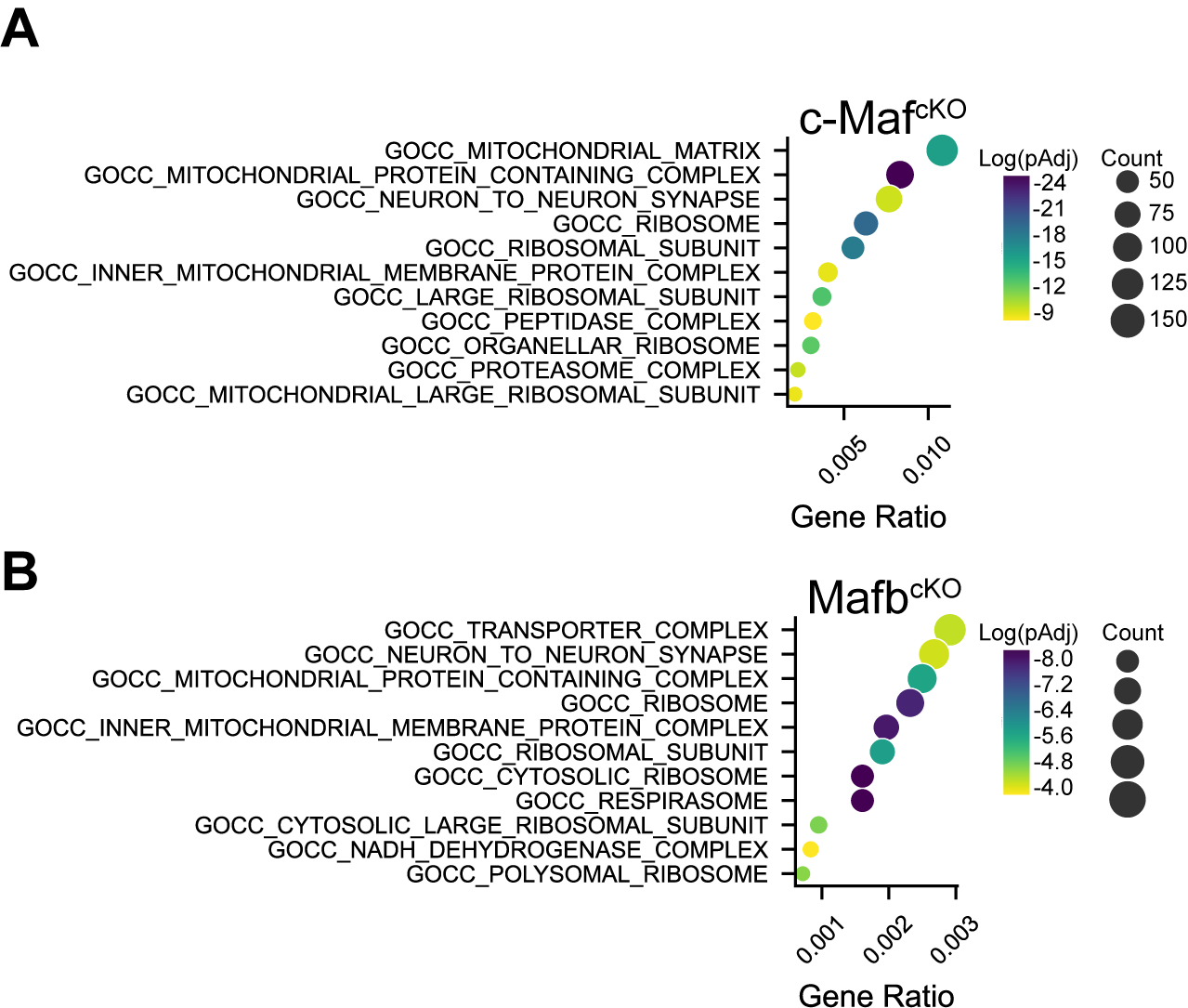

### Supplemental Figure 7

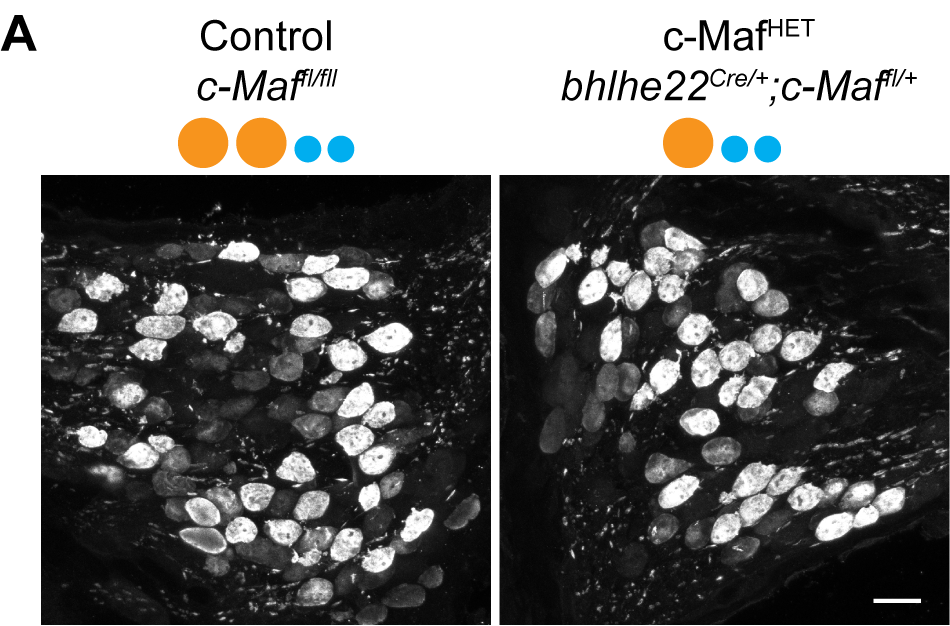
