## Supplemental Table 2 for "Combinatorial transcriptional regulation establishes subtype-appropriate synaptic properties in auditory neurons"

| **ID** | **Description** | **GeneRatio** | **BgRatio** | **pvalue** | **p.adjust** | **qvalue** | **geneID** | **Count** |
| --- | --- | --- | --- | --- | --- | --- | --- | --- |
| **GOCC_CYTOSOLIC_RIBOSOME** | GOCC_CYTOSOLIC_RIBOSOME | 19/377 | 95/16819 | 2.77E-13 | 1.7E-09 | 1.55E-09 | Rps2/Rpl39/Rpl35/Rpl37/Rps21/Rps29/Rps28/Rpl21/Rpl37a/Rpl38/Rpl36/Rps26/Rps27/Rps15a/Rpl36a/Rps17/Rpl31/Isg15/Apod | 19 |
| **GOCC_RIBOSOME** | GOCC_RIBOSOME | 24/377 | 201/16819 | 2.45E-11 | 7.21E-08 | 6.57E-08 | Rps2/Rpl39/Rpl35/Rpl37/Rps21/Rps29/Rps28/Rpl21/Rpl37a/Rpl38/Rpl36/Rpl22l1/Rps26/Rps27/Rps15a/Mt3/Rpl36a/Rps17/Mrps5/Dap3/Rpl31/Dhx9/Isg15/Apod | 24 |
| **GOMF_STRUCTURAL_CONSTITUENT_OF_RIBOSOME** | GOMF_STRUCTURAL_CONSTITUENT_OF_RIBOSOME | 21/377 | 154/16819 | 3.53E-11 | 7.21E-08 | 6.57E-08 | Rps2/Rpl39/Rpl35/Rpl37/Rps21/Rps29/Rps28/Rpl21/Rpl37a/Rpl38/Rpl36/Rpl22l1/Rps26/Rps27/Rps15a/Rpl36a/Rps17/Mrps5/Dap3/Rpl31/Isg15 | 21 |
| **GOBP_CYTOPLASMIC_TRANSLATION** | GOBP_CYTOPLASMIC_TRANSLATION | 20/377 | 143/16819 | 6.54E-11 | 1E-07 | 9.12E-08 | Rps2/Rpl39/Rpl35/Rpl37/Rps21/Rps29/Rps28/Rpl21/Rpl37a/Rpl38/Rpl36/Rpl22l1/Rps26/Rps27/Rps15a/Rpl36a/Rps17/Rpl31/Dhx9/Cpeb3 | 20 |
| **GOCC_POLYSOMAL_RIBOSOME** | GOCC_POLYSOMAL_RIBOSOME | 10/377 | 30/16819 | 5.73E-10 | 7.01E-07 | 6.39E-07 | Rpl39/Rps21/Rps29/Rps28/Rpl38/Rpl36/Rps26/Rpl36a/Rpl31/Dhx9 | 10 |
| **GOCC_NEURON_TO_NEURON_SYNAPSE** | GOCC_NEURON_TO_NEURON_SYNAPSE | 29/377 | 344/16819 | 1E-09 | 1.02E-06 | 9.3E-07 | Kcnd2/Sipa1l1/Clstn2/Cacna1c/Rpl38/Cabp1/Rps27/Lrrc4c/Mt3/Plekha5/Kpna1/Nrcam/Pkp4/Epha7/Tmem108/Calb1/Actr2/Ppp1r9a/Cryab/Cpeb3/Mpdz/Pde4b/Ptprd/Dcc/Add3/Drp2/Insyn1/Dlgap2/Slc8a1 | 29 |
| **GOCC_RIBOSOMAL_SUBUNIT** | GOCC_RIBOSOMAL_SUBUNIT | 20/377 | 171/16819 | 1.67E-09 | 1.46E-06 | 1.33E-06 | Rps2/Rpl39/Rpl35/Rpl37/Rps21/Rps29/Rps28/Rpl21/Rpl37a/Rpl38/Rpl36/Rps26/Rps27/Rps15a/Rpl36a/Rps17/Mrps5/Dap3/Rpl31/Isg15 | 20 |
| **GOBP_AXON_DEVELOPMENT** | GOBP_AXON_DEVELOPMENT | 33/377 | 474/16819 | 8.76E-09 | 6.7E-06 | 6.11E-06 | Scn1b/Ndn/Alcam/Spp1/Ncam2/Sipa1l1/Adnp/Lrrc4c/Mt3/Tnr/Rab21/Auts2/Nfib/S100b/Nrcam/Epha7/Stk25/Nrxn3/Sema3a/Plp1/Sema5a/Lgi1/Pak1/Cdk5r2/Pgrmc1/Dcc/Crtac1/Nrp2/Nrep/Lama2/Sema6a/Robo2/Apod | 33 |
| **HP_ABNORMALITY_OF_THE_THENAR_EMINENCE** | HP_ABNORMALITY_OF_THE_THENAR_EMINENCE | 9/377 | 30/16819 | 1.23E-08 | 8.33E-06 | 7.6E-06 | Rpl35/Rps29/Rps28/Rps26/Rps27/Rps15a/Rps17/Rpl31/Fgf10 | 9 |
| **HP_ERYTHROID_HYPOPLASIA** | HP_ERYTHROID_HYPOPLASIA | 8/377 | 25/16819 | 4.59E-08 | 2.55E-05 | 2.33E-05 | Rpl35/Rps29/Rps28/Rps26/Rps27/Rps15a/Rps17/Rpl31 | 8 |
| **HP_PURE_RED_CELL_APLASIA** | HP_PURE_RED_CELL_APLASIA | 8/377 | 25/16819 | 4.59E-08 | 2.55E-05 | 2.33E-05 | Rpl35/Rps29/Rps28/Rps26/Rps27/Rps15a/Rps17/Rpl31 | 8 |
| **HP_MALIGNANT_GENITOURINARY_TRACT_TUMOR** | HP_MALIGNANT_GENITOURINARY_TRACT_TUMOR | 8/377 | 26/16819 | 6.49E-08 | 3.31E-05 | 3.02E-05 | Rpl35/Rps29/Rps28/Rps26/Rps27/Rps15a/Rps17/Rpl31 | 8 |
| **HP_ABSENT_THUMB** | HP_ABSENT_THUMB | 10/377 | 49/16819 | 1.07E-07 | 5.05E-05 | 4.6E-05 | Rpl35/Rps29/Rps28/Rps26/Rps27/Rps15a/Rps17/Rpl31/Sf3b4/Fgf10 | 10 |
| **HP_INCREASED_MEAN_CORPUSCULAR_VOLUME** | HP_INCREASED_MEAN_CORPUSCULAR_VOLUME | 9/377 | 38/16819 | 1.19E-07 | 5.21E-05 | 4.75E-05 | Rpl35/Ankrd11/Rps29/Rps28/Rps26/Rps27/Rps15a/Rps17/Rpl31 | 9 |
| **HP_PARTIAL_DUPLICATION_OF_THE_PHALANX_OF_HAND** | HP_PARTIAL_DUPLICATION_OF_THE_PHALANX_OF_HAND | 9/377 | 39/16819 | 1.52E-07 | 6.2E-05 | 5.65E-05 | Rpl35/Rps29/Rps28/Rps26/Rps27/Rps15a/Rps17/Rpl31/Fgf10 | 9 |
| **HP_PERSISTENCE_OF_HEMOGLOBIN_F** | HP_PERSISTENCE_OF_HEMOGLOBIN_F | 8/377 | 29/16819 | 1.68E-07 | 6.43E-05 | 5.86E-05 | Rpl35/Rps29/Rps28/Rps26/Rps27/Rps15a/Rps17/Rpl31 | 8 |
| **GOCC_CYTOSOLIC_SMALL_RIBOSOMAL_SUBUNIT** | GOCC_CYTOSOLIC_SMALL_RIBOSOMAL_SUBUNIT | 9/377 | 40/16819 | 1.92E-07 | 6.92E-05 | 6.3E-05 | Rps2/Rps21/Rps29/Rps28/Rps26/Rps27/Rps15a/Rps17/Isg15 | 9 |
| **GOCC_SMALL_RIBOSOMAL_SUBUNIT** | GOCC_SMALL_RIBOSOMAL_SUBUNIT | 11/377 | 66/16819 | 2.21E-07 | 7.24E-05 | 6.6E-05 | Rps2/Rps21/Rps29/Rps28/Rps26/Rps27/Rps15a/Rps17/Mrps5/Dap3/Isg15 | 11 |
| **HP_ADENOCARCINOMA_OF_THE_COLON** | HP_ADENOCARCINOMA_OF_THE_COLON | 8/377 | 30/16819 | 2.25E-07 | 7.24E-05 | 6.6E-05 | Rpl35/Rps29/Rps28/Rps26/Rps27/Rps15a/Rps17/Rpl31 | 8 |
| **HP_TRIPHALANGEAL_THUMB** | HP_TRIPHALANGEAL_THUMB | 11/377 | 67/16819 | 2.59E-07 | 7.92E-05 | 7.22E-05 | Rpl35/Mafb/Rps29/Rps28/Rps26/Rps27/Rps15a/Rps17/Rpl31/Sf3b4/Fgf10 | 11 |
| **HP_THROMBOCYTOSIS** | HP_THROMBOCYTOSIS | 10/377 | 55/16819 | 3.39E-07 | 9.87E-05 | 8.99E-05 | Rpl35/Rps29/Rps28/Rps26/Rps27/Rps15a/Cd55/Rps17/Rpl31/Hmgcl | 10 |
| **HP_ABNORMAL_ERYTHROCYTE_ENZYME_LEVEL** | HP_ABNORMAL_ERYTHROCYTE_ENZYME_LEVEL | 8/377 | 32/16819 | 3.89E-07 | 0.000108116 | 9.86E-05 | Rpl35/Rps29/Rps28/Rps26/Rps27/Rps15a/Rps17/Rpl31 | 8 |
| **HP_OSTEOSARCOMA** | HP_OSTEOSARCOMA | 8/377 | 33/16819 | 5.03E-07 | 0.000128281 | 0.000116937 | Rpl35/Rps29/Rps28/Rps26/Rps27/Rps15a/Rps17/Rpl31 | 8 |
| **HP_RETICULOCYTOPENIA** | HP_RETICULOCYTOPENIA | 8/377 | 33/16819 | 5.03E-07 | 0.000128281 | 0.000116937 | Rpl35/Rps29/Rps28/Rps26/Rps27/Rps15a/Rps17/Rpl31 | 8 |
| **HP_NORMOCHROMIC_ANEMIA** | HP_NORMOCHROMIC_ANEMIA | 8/377 | 34/16819 | 6.45E-07 | 0.000157916 | 0.000143951 | Rpl35/Rps29/Rps28/Rps26/Rps27/Rps15a/Rps17/Rpl31 | 8 |
| **HP_APLASIA_HYPOPLASIA_OF_THE_EAR** | HP_APLASIA_HYPOPLASIA_OF_THE_EAR | 16/377 | 162/16819 | 7.55E-07 | 0.000177613 | 0.000161907 | Maf/Rpl35/Plcb4/Rps29/Rps28/Rtl1/Rps26/Rps27/Rps15a/Adnp/Rps17/Rpl31/Sf3b4/Fgf10/Fkrp/Pak1 | 16 |
| **HP_ABNORMAL_MEAN_CORPUSCULAR_VOLUME** | HP_ABNORMAL_MEAN_CORPUSCULAR_VOLUME | 9/377 | 48/16819 | 1.01E-06 | 0.000227964 | 0.000207805 | Rpl35/Ankrd11/Rps29/Rps28/Rps26/Rps27/Rps15a/Rps17/Rpl31 | 9 |
| **GOBP_REGULATION_OF_NEURON_PROJECTION_DEVELOPMENT** | GOBP_REGULATION_OF_NEURON_PROJECTION_DEVELOPMENT | 27/377 | 423/16819 | 1.14E-06 | 0.00024879 | 0.00022679 | Scn1b/Spp1/Sipa1l1/Sez6/Adnp/Lrrc4c/Mt3/Tnr/Rab21/Zfp804a/Ndrg4/Nrcam/Kdm1a/Epha7/Stk25/Actr2/Sema3a/Sema5a/Il1rapl1/Pak1/Camk1d/Ptprd/Id1/Dcc/Rgs2/Sema6a/Robo2 | 27 |
| **GOCC_POLYSOME** | GOCC_POLYSOME | 10/377 | 63/16819 | 1.26E-06 | 0.00026617 | 0.000242633 | Rpl39/Rps21/Rps29/Rps28/Rpl38/Rpl36/Rps26/Rpl36a/Rpl31/Dhx9 | 10 |
| **HP_ADENOCARCINOMA_OF_THE_INTESTINES** | HP_ADENOCARCINOMA_OF_THE_INTESTINES | 8/377 | 38/16819 | 1.61E-06 | 0.000327738 | 0.000298756 | Rpl35/Rps29/Rps28/Rps26/Rps27/Rps15a/Rps17/Rpl31 | 8 |
| **HP_ABNORMAL_SCAPULA_MORPHOLOGY** | HP_ABNORMAL_SCAPULA_MORPHOLOGY | 15/377 | 153/16819 | 1.84E-06 | 0.000362813 | 0.000330729 | Rpl35/Ttn/Rps29/Rps28/Rtl1/Rps26/Rps27/Rps15a/Dnm1l/Rps17/Rpl31/Sf3b4/Gfpt1/Tk2/Neb | 15 |
| **HP_CLEFT_SOFT_PALATE** | HP_CLEFT_SOFT_PALATE | 16/377 | 174/16819 | 1.96E-06 | 0.000374565 | 0.000341442 | Snrpn/Ubb/Rpl35/Ttn/Plcb4/Rps29/Rps28/Rtl1/Rps26/Rps27/Rps15a/Sms/Rps17/Rpl31/Fgf10/Fkrp | 16 |
| **HP_SPRENGEL_ANOMALY** | HP_SPRENGEL_ANOMALY | 9/377 | 52/16819 | 2.04E-06 | 0.000378056 | 0.000344624 | Rpl35/Rps29/Rps28/Rps26/Rps27/Rps15a/Rps17/Rpl31/Sf3b4 | 9 |
| **GOBP_DEVELOPMENTAL_CELL_GROWTH** | GOBP_DEVELOPMENTAL_CELL_GROWTH | 18/377 | 219/16819 | 2.3E-06 | 0.000413299 | 0.000376751 | Ndn/Alcam/Spp1/Sorbs2/Adnp/Mt3/Tnr/Rab21/Auts2/Nrcam/Epha7/Tmem108/Sema3a/Sema5a/Dcc/Rgs2/Nrp2/Sema6a | 18 |
| **HP_DUPLICATION_OF_THUMB_PHALANX** | HP_DUPLICATION_OF_THUMB_PHALANX | 9/377 | 53/16819 | 2.41E-06 | 0.000420963 | 0.000383737 | Rpl35/Rps29/Rps28/Rps26/Rps27/Rps15a/Rps17/Rpl31/Fgf10 | 9 |
| **GOCC_CYTOSOLIC_LARGE_RIBOSOMAL_SUBUNIT** | GOCC_CYTOSOLIC_LARGE_RIBOSOMAL_SUBUNIT | 9/377 | 54/16819 | 2.83E-06 | 0.000479603 | 0.000437192 | Rpl39/Rpl35/Rpl37/Rpl21/Rpl37a/Rpl38/Rpl36/Rpl36a/Rpl31 | 9 |
| **HP_ABNORMAL_NUMBER_OF_ERYTHROID_PRECURSORS** | HP_ABNORMAL_NUMBER_OF_ERYTHROID_PRECURSORS | 8/377 | 41/16819 | 2.96E-06 | 0.000479603 | 0.000437192 | Rpl35/Rps29/Rps28/Rps26/Rps27/Rps15a/Rps17/Rpl31 | 8 |
| **HP_APLASIA_OF_THE_FINGERS** | HP_APLASIA_OF_THE_FINGERS | 10/377 | 69/16819 | 2.98E-06 | 0.000479603 | 0.000437192 | Rpl35/Rps29/Rps28/Rps26/Rps27/Rps15a/Rps17/Rpl31/Sf3b4/Fgf10 | 10 |
| **HP_GASTROINTESTINAL_CARCINOMA** | HP_GASTROINTESTINAL_CARCINOMA | 10/377 | 71/16819 | 3.89E-06 | 0.000609817 | 0.000555891 | Rpl35/Rps29/Rps28/Rps26/Rps27/Rps15a/Rps17/Rpl31/Dlc1/Dcc | 10 |
| **GOBP_DEVELOPMENTAL_GROWTH_INVOLVED_IN_MORPHOGENESIS** | GOBP_DEVELOPMENTAL_GROWTH_INVOLVED_IN_MORPHOGENESIS | 18/377 | 233/16819 | 5.51E-06 | 0.000842172 | 0.000767698 | Ndn/Alcam/Spp1/Adnp/Mt3/Tnr/Rab21/Auts2/Nrcam/Epha7/Tmem108/Fgf10/Sema3a/Sema5a/Dcc/Nrp2/Sema6a/Fgf1 | 18 |
| **HP_WEBBED_NECK** | HP_WEBBED_NECK | 12/377 | 110/16819 | 6.59E-06 | 0.000977661 | 0.000891206 | Rpl35/Ankrd11/Mafb/Rps29/Rps28/Rps26/Rps27/Rps15a/Sms/Hras/Rps17/Rpl31 | 12 |
| **GOBP_AXONAL_FASCICULATION** | GOBP_AXONAL_FASCICULATION | 6/377 | 22/16819 | 6.71E-06 | 0.000977661 | 0.000891206 | Ndn/Ncam2/Nrcam/Sema3a/Sema5a/Crtac1 | 6 |
| **HP_APLASIA_HYPOPLASIA_OF_THE_THUMB** | HP_APLASIA_HYPOPLASIA_OF_THE_THUMB | 13/377 | 130/16819 | 7.2E-06 | 0.001019557 | 0.000929397 | Rpl35/Mafb/Rps29/Rps28/Rtl1/Rps26/Rps27/Rps15a/Rps17/Kdm1a/Rpl31/Sf3b4/Fgf10 | 13 |
| **HP_NONIMMUNE_HYDROPS_FETALIS** | HP_NONIMMUNE_HYDROPS_FETALIS | 8/377 | 46/16819 | 7.33E-06 | 0.001019557 | 0.000929397 | Rpl35/Rps29/Rps28/Rps26/Rps27/Rps15a/Rps17/Rpl31 | 8 |
| **HP_HYPOVENTILATION** | HP_HYPOVENTILATION | 8/377 | 47/16819 | 8.66E-06 | 0.001178141 | 0.001073958 | Snrpn/Ndn/Ttn/Pura/Dmd/Fkrp/Lama2/Neb | 8 |
| **GOBP_REGULATION_OF_AXONOGENESIS** | GOBP_REGULATION_OF_AXONOGENESIS | 14/377 | 154/16819 | 9.77E-06 | 0.001273201 | 0.001160611 | Spp1/Sipa1l1/Adnp/Lrrc4c/Mt3/Tnr/Epha7/Stk25/Sema3a/Sema5a/Pak1/Dcc/Sema6a/Robo2 | 14 |
| **HP_ABNORMAL_SOFT_PALATE_MORPHOLOGY** | HP_ABNORMAL_SOFT_PALATE_MORPHOLOGY | 17/377 | 220/16819 | 9.99E-06 | 0.001273201 | 0.001160611 | Snrpn/Ubb/Rpl35/Ttn/Plcb4/Rps29/Rps28/Rtl1/Rps26/Rps27/Rps15a/Sms/Rps17/Rpl31/Sf3b4/Fgf10/Fkrp | 17 |
| **HP_ABNORMALITY_OF_THE_MUSCULATURE_OF_THE_UPPER_LIMBS** | HP_ABNORMALITY_OF_THE_MUSCULATURE_OF_THE_UPPER_LIMBS | 17/377 | 220/16819 | 9.99E-06 | 0.001273201 | 0.001160611 | Rpl35/Ttn/Rps29/Rps28/Rps26/Rps27/Rps15a/Dnm1l/Rps17/Rpl31/Fgf10/Gfpt1/Adssl1/Tk2/Fkrp/Lama2/Neb | 17 |
| **HP_ABNORMALITY_OF_THE_MUSCULATURE_OF_THE_HAND** | HP_ABNORMALITY_OF_THE_MUSCULATURE_OF_THE_HAND | 11/377 | 97/16819 | 1.1E-05 | 0.001373292 | 0.001251851 | Rpl35/Rps29/Rps28/Rps26/Rps27/Rps15a/Rps17/Rpl31/Fgf10/Tk2/Neb | 11 |
| **GOBP_CELL_GROWTH** | GOBP_CELL_GROWTH | 27/377 | 480/16819 | 1.19E-05 | 0.00141722 | 0.001291895 | Rasgrp2/Ndn/Alcam/Spp1/Egln2/Sorbs2/Adnp/Mt3/Tnr/Rab21/Csnk2a1/Auts2/Phb/Nrcam/Eno1/Epha7/Tmem108/Sema3a/Tead1/Cryab/Sema5a/Lgi1/Dcc/Rgs2/Nrp2/Sema6a/Igfbp5 | 27 |
| **GOBP_NEURON_RECOGNITION** | GOBP_NEURON_RECOGNITION | 8/377 | 49/16819 | 1.19E-05 | 0.00141722 | 0.001291895 | Ndn/Ncam2/Nrcam/Sema3a/Sema5a/Crtac1/Ntm/Robo2 | 8 |
| **HP_MACROCYTIC_ANEMIA** | HP_MACROCYTIC_ANEMIA | 9/377 | 64/16819 | 1.2E-05 | 0.00141722 | 0.001291895 | Rpl35/Rps29/Rps28/Rps26/Rps27/Rps15a/Dnm1l/Rps17/Rpl31 | 9 |
| **HP_DEVELOPMENTAL_GLAUCOMA** | HP_DEVELOPMENTAL_GLAUCOMA | 9/377 | 65/16819 | 1.37E-05 | 0.001582445 | 0.001442509 | Rpl35/Rps29/Rps28/Rps26/Rps27/Rps15a/Rps17/Rpl31/Fkrp | 9 |
| **GOBP_AXON_EXTENSION** | GOBP_AXON_EXTENSION | 12/377 | 121/16819 | 1.76E-05 | 0.001993191 | 0.001816932 | Ndn/Alcam/Adnp/Mt3/Tnr/Rab21/Auts2/Nrcam/Sema3a/Sema5a/Nrp2/Sema6a | 12 |
| **HP_SHORT_THUMB** | HP_SHORT_THUMB | 11/377 | 104/16819 | 2.14E-05 | 0.002385621 | 0.00217466 | Rpl35/Rps29/Rps28/Rtl1/Rps26/Rps27/Rps15a/Rps17/Kdm1a/Rpl31/Fgf10 | 11 |
| **GOBP_NEURON_PROJECTION_GUIDANCE** | GOBP_NEURON_PROJECTION_GUIDANCE | 17/377 | 234/16819 | 2.23E-05 | 0.002436101 | 0.002220676 | Scn1b/Alcam/Tnr/Nfib/Nrcam/Epha7/Nrxn3/Sema3a/Sema5a/Lgi1/Cdk5r2/Pgrmc1/Dcc/Nrp2/Lama2/Sema6a/Robo2 | 17 |
| **HP_ABNORMAL_HEMOGLOBIN** | HP_ABNORMAL_HEMOGLOBIN | 8/377 | 54/16819 | 2.5E-05 | 0.002668265 | 0.00243231 | Rpl35/Rps29/Rps28/Rps26/Rps27/Rps15a/Rps17/Rpl31 | 8 |
| **HP_PALLOR** | HP_PALLOR | 13/377 | 146/16819 | 2.53E-05 | 0.002668265 | 0.00243231 | Scn1b/Rpl35/Rps29/Scn9a/Rps28/Rps26/Rps27/Rps15a/Rps17/Rpl31/Hmgcl/Cdh23/Klhl7 | 13 |
| **HP_DEVELOPMENTAL_CATARACT** | HP_DEVELOPMENTAL_CATARACT | 12/377 | 127/16819 | 2.87E-05 | 0.00297167 | 0.002708884 | Maf/mt-Nd6/Rpl35/Rps29/Rps28/Rps26/Rps27/Rps15a/Rps17/Rpl31/Vps4a/Cryab | 12 |
| **GOBP_REGULATION_OF_EXTENT_OF_CELL_GROWTH** | GOBP_REGULATION_OF_EXTENT_OF_CELL_GROWTH | 11/377 | 108/16819 | 3.06E-05 | 0.003073648 | 0.002801844 | Spp1/Adnp/Mt3/Tnr/Rab21/Nrcam/Epha7/Sema3a/Sema5a/Dcc/Sema6a | 11 |
| **GOCC_GLUTAMATERGIC_SYNAPSE** | GOCC_GLUTAMATERGIC_SYNAPSE | 20/377 | 315/16819 | 3.06E-05 | 0.003073648 | 0.002801844 | Kcnd2/Clstn2/Lrrc4c/Plekha5/Hras/Tnr/Kpna1/Pura/Fxyd6/Nrcam/Epha7/Calb1/Lgi1/Il1rapl1/Ptprd/Drp2/Nrp2/Bcan/Dlgap2/Cdh8 | 20 |
| **GOBP_REGULATION_OF_CELL_SIZE** | GOBP_REGULATION_OF_CELL_SIZE | 14/377 | 180/16819 | 5.62E-05 | 0.005548068 | 0.00505745 | Spp1/Adnp/Mt3/Tnr/Slc12a2/Rab21/Nrcam/Kdm1a/Epha7/Sema3a/Ano6/Sema5a/Dcc/Sema6a | 14 |
| **GOCC_TRANSPORTER_COMPLEX** | GOCC_TRANSPORTER_COMPLEX | 22/377 | 386/16819 | 6.35E-05 | 0.006170436 | 0.005624782 | mt-Nd6/Scn1b/mt-Nd4l/Kcnab1/Scn9a/Kcnd2/Cacna1c/mt-Nd4/Ndufc2/Hcn2/mt-Nd3/Uqcr10/Ano6/Kcns3/Atp10a/Best3/Pde4b/Cacna1e/Kcnip4/Dpp6/Cachd1/Kcnc2 | 22 |
| **GOBP_NEURAL_NUCLEUS_DEVELOPMENT** | GOBP_NEURAL_NUCLEUS_DEVELOPMENT | 8/377 | 62/16819 | 6.95E-05 | 0.006647422 | 0.006059588 | Ckb/Basp1/Nfib/Kirrel3/Plp1/Cdk5r2/Padi2/Kcnc2 | 8 |
| **HP_NARROW_PALM** | HP_NARROW_PALM | 4/377 | 11/16819 | 7.24E-05 | 0.006813191 | 0.006210698 | Snrpn/Ndn/Sms/Auts2 | 4 |
| **HP_SMALL_FOR_GESTATIONAL_AGE** | HP_SMALL_FOR_GESTATIONAL_AGE | 16/377 | 234/16819 | 8E-05 | 0.007412624 | 0.006757123 | Snrpn/Ndn/Rpl35/Rps29/Rps28/Rtl1/Rps26/Rps27/Rps15a/Auts2/Rps17/Dmd/Rpl31/Il1rapl1/Wdr73/Stambp | 16 |
| **GOBP_NEGATIVE_REGULATION_OF_AXONOGENESIS** | GOBP_NEGATIVE_REGULATION_OF_AXONOGENESIS | 8/377 | 64/16819 | 8.75E-05 | 0.007995039 | 0.007288035 | Spp1/Mt3/Tnr/Epha7/Sema3a/Sema5a/Dcc/Sema6a | 8 |
| **HP_GENERALIZED_ONSET_SEIZURE** | HP_GENERALIZED_ONSET_SEIZURE | 20/377 | 343/16819 | 0.000100065 | 0.009004349 | 0.008208092 | mt-Nd6/Scn1b/Scn9a/mt-Nd4/Dnm1l/Sms/Slc12a2/Pcyt2/Pura/Hcn2/mt-Nd3/Dmd/Kirrel3/Trpm3/Lnpk/Plp1/Tk2/Lgi1/Il1rapl1/Lama2 | 20 |
| **HP_ACUTE_MYELOID_LEUKEMIA** | HP_ACUTE_MYELOID_LEUKEMIA | 8/377 | 66/16819 | 0.000109262 | 0.009689506 | 0.00883266 | Rpl35/Rps29/Rps28/Rps26/Rps27/Rps15a/Rps17/Rpl31 | 8 |
| **GOCC_INNER_MITOCHONDRIAL_MEMBRANE_PROTEIN_COMPLEX** | GOCC_INNER_MITOCHONDRIAL_MEMBRANE_PROTEIN_COMPLEX | 12/377 | 146/16819 | 0.000112125 | 0.009763258 | 0.00889989 | mt-Nd6/mt-Atp8/mt-Nd4l/Atp5md/mt-Nd4/Ndufc2/mt-Nd3/Phb/Cox7c/Atp5mpl/Uqcr10/Romo1 | 12 |
| **GOBP_REGULATION_OF_NERVOUS_SYSTEM_DEVELOPMENT** | GOBP_REGULATION_OF_NERVOUS_SYSTEM_DEVELOPMENT | 23/377 | 430/16819 | 0.000113285 | 0.009763258 | 0.00889989 | Spp1/Clstn2/Adnp/Mt3/Tnr/Rab21/Kdm1a/Rxrg/Epha7/Stk25/Actr2/Sema3a/Sema5a/Il1rapl1/Id4/Ptprd/Id1/Dcc/Tenm4/Eif2ak3/Sema6a/Hey2/Robo2 | 23 |
| **GOBP_REGULATION_OF_CELLULAR_COMPONENT_SIZE** | GOBP_REGULATION_OF_CELLULAR_COMPONENT_SIZE | 20/377 | 349/16819 | 0.000126503 | 0.010751022 | 0.009800306 | Spp1/Adnp/Mt3/Tnr/Slc12a2/Rab21/Dstn/Nrcam/Kdm1a/Epha7/Arfgef1/Sema3a/Ano6/Pdxp/Sema5a/Ush1c/Dcc/Add3/Sema6a/Neb | 20 |
| **GOBP_NEURON_PROJECTION_EXTENSION** | GOBP_NEURON_PROJECTION_EXTENSION | 13/377 | 172/16819 | 0.000137519 | 0.011527119 | 0.010507772 | Ndn/Alcam/Adnp/Mt3/Tnr/Rab21/Auts2/Nrcam/Tmem108/Sema3a/Sema5a/Nrp2/Sema6a | 13 |
| **GOBP_REGULATION_OF_NEUROGENESIS** | GOBP_REGULATION_OF_NEUROGENESIS | 20/377 | 352/16819 | 0.000141903 | 0.011640441 | 0.010611073 | Spp1/Adnp/Mt3/Tnr/Rab21/Kdm1a/Epha7/Stk25/Actr2/Sema3a/Sema5a/Il1rapl1/Id4/Ptprd/Id1/Dcc/Tenm4/Sema6a/Hey2/Robo2 | 20 |
| **GOBP_ATP_SYNTHESIS_COUPLED_ELECTRON_TRANSPORT** | GOBP_ATP_SYNTHESIS_COUPLED_ELECTRON_TRANSPORT | 9/377 | 87/16819 | 0.000142676 | 0.011640441 | 0.010611073 | mt-Nd6/mt-Nd4l/Ghitm/mt-Nd4/Ndufc2/Cycs/mt-Nd3/Cox7c/Uqcr10 | 9 |
| **HP_NEOPLASM_OF_THE_LARGE_INTESTINE** | HP_NEOPLASM_OF_THE_LARGE_INTESTINE | 10/377 | 107/16819 | 0.000144847 | 0.011662108 | 0.010630824 | Rpl35/Rps29/Rps28/Rps26/Rps27/Rps15a/Rps17/Rpl31/Dlc1/Dcc | 10 |
| **GOBP_NEGATIVE_REGULATION_OF_SMOOTH_MUSCLE_CELL_PROLIFERATION** | GOBP_NEGATIVE_REGULATION_OF_SMOOTH_MUSCLE_CELL_PROLIFERATION | 7/377 | 52/16819 | 0.000150909 | 0.011992394 | 0.010931903 | Prkg1/Mef2c/Tafa5/Ndrg4/Igfbp5/Ogn/Apod | 7 |
| **GOBP_NEGATIVE_REGULATION_OF_SMOOTH_MUSCLE_CELL_MIGRATION** | GOBP_NEGATIVE_REGULATION_OF_SMOOTH_MUSCLE_CELL_MIGRATION | 5/377 | 24/16819 | 0.000164804 | 0.012928678 | 0.01178539 | Prkg1/Mef2c/Tafa5/Ndrg4/Igfbp5 | 5 |
| **HP_ABNORMALITY_OF_THE_VOICE** | HP_ABNORMALITY_OF_THE_VOICE | 24/377 | 477/16819 | 0.000203753 | 0.015781795 | 0.014386205 | mt-Nd6/Snrpn/Ubb/Ndn/Ttn/Plcb4/Rtl1/mt-Nd4/Dync1i2/Sms/Hras/Slc12a2/mt-Nd3/Gfpt1/Sema3a/Cryab/Sema5a/Tk2/Il1rapl1/Ttc19/Fkrp/Ghr/Dcc/Lama2 | 24 |
| **HP_LEUKOPENIA** | HP_LEUKOPENIA | 10/377 | 113/16819 | 0.000227333 | 0.017388133 | 0.015850494 | Rpl35/Rps29/Rps28/Spp1/Rps26/Rps27/Rps15a/Rps17/Rpl31/Hmgcl | 10 |
| **GOBP_REGULATION_OF_ANATOMICAL_STRUCTURE_SIZE** | GOBP_REGULATION_OF_ANATOMICAL_STRUCTURE_SIZE | 24/377 | 483/16819 | 0.00024511 | 0.018516412 | 0.016878999 | Prkg1/Spp1/Adnp/Mt3/Tnr/Slc12a2/Rab21/Dstn/Nrcam/Kdm1a/Epha7/Arfgef1/Sema3a/Ano6/Pdxp/Sema5a/Zdhhc21/Ush1c/Dcc/Add3/Rgs2/Sema6a/Slc8a1/Neb | 24 |
| **GOCC_SYNAPTIC_MEMBRANE** | GOCC_SYNAPTIC_MEMBRANE | 20/377 | 370/16819 | 0.000274025 | 0.020309035 | 0.0185131 | Rph3a/Kcnd2/Clstn2/Cacna1c/Lrrc4c/Fxyd6/Nrcam/Epha7/Dmd/Tmem240/Cryab/Il1rapl1/Itsn1/Ptprd/Dcc/Utrn/Nrp2/Slc8a1/Cdh8/Kcnc2 | 20 |
| **GOBP_NEURON_CELL_CELL_ADHESION** | GOBP_NEURON_CELL_CELL_ADHESION | 4/377 | 15/16819 | 0.000278797 | 0.020309035 | 0.0185131 | Ncam2/Astn2/Tnr/Nrxn3 | 4 |
| **HP_MUSCLE_FIBER_SPLITTING** | HP_MUSCLE_FIBER_SPLITTING | 4/377 | 15/16819 | 0.000278797 | 0.020309035 | 0.0185131 | Ttn/Adssl1/Cryab/Neb | 4 |
| **GOMF_STRUCTURAL_CONSTITUENT_OF_MUSCLE** | GOMF_STRUCTURAL_CONSTITUENT_OF_MUSCLE | 6/377 | 41/16819 | 0.0002823 | 0.02032231 | 0.018525202 | Ttn/Myl6b/Sorbs2/Synm/Dmd/Neb | 6 |
| **GOBP_REGULATION_OF_SYNAPSE_STRUCTURE_OR_ACTIVITY** | GOBP_REGULATION_OF_SYNAPSE_STRUCTURE_OR_ACTIVITY | 14/377 | 210/16819 | 0.00028683 | 0.020408321 | 0.018603607 | Mef2c/Sipa1l1/Clstn2/Adnp/Zfp804a/Nrcam/Epha7/Actr2/Il1rapl1/Ptprd/Pgrmc1/Nrp2/Robo2/Cdh8 | 14 |
| **HP_ABNORMAL_RETICULOCYTE_MORPHOLOGY** | HP_ABNORMAL_RETICULOCYTE_MORPHOLOGY | 8/377 | 76/16819 | 0.000295373 | 0.020774547 | 0.018937447 | Rpl35/Rps29/Rps28/Rps26/Rps27/Rps15a/Rps17/Rpl31 | 8 |
| **GOCC_RESPIRASOME** | GOCC_RESPIRASOME | 9/377 | 96/16819 | 0.000302316 | 0.02102127 | 0.019162352 | mt-Nd6/mt-Nd4l/mt-Nd4/Ndufc2/Cycs/mt-Nd3/Cox7c/Uqcr10/Ttc19 | 9 |
| **GOBP_NEURON_MIGRATION** | GOBP_NEURON_MIGRATION | 12/377 | 163/16819 | 0.000314625 | 0.021631369 | 0.0197185 | Prkg1/Ndn/Mef2c/Astn2/Auts2/Nrcam/Kirrel3/Sema3a/Cdk5r2/Dcc/Nrp2/Sema6a | 12 |
| **GOBP_REGULATION_OF_DENDRITE_DEVELOPMENT** | GOBP_REGULATION_OF_DENDRITE_DEVELOPMENT | 9/377 | 97/16819 | 0.000326772 | 0.022216882 | 0.020252236 | Sipa1l1/Sez6/Rab21/Actr2/Il1rapl1/Camk1d/Ptprd/Id1/Dcc | 9 |
| **GOBP_NEGATIVE_REGULATION_OF_NERVOUS_SYSTEM_DEVELOPMENT** | GOBP_NEGATIVE_REGULATION_OF_NERVOUS_SYSTEM_DEVELOPMENT | 11/377 | 141/16819 | 0.000338467 | 0.022759142 | 0.020746543 | Spp1/Mt3/Tnr/Epha7/Sema3a/Sema5a/Id4/Id1/Dcc/Sema6a/Robo2 | 11 |
| **HP_NEOPLASM_OF_THE_COLON** | HP_NEOPLASM_OF_THE_COLON | 8/377 | 78/16819 | 0.000353315 | 0.023499299 | 0.021421248 | Rpl35/Rps29/Rps28/Rps26/Rps27/Rps15a/Rps17/Rpl31 | 8 |
| **HP_RAGGED_RED_MUSCLE_FIBERS** | HP_RAGGED_RED_MUSCLE_FIBERS | 6/377 | 43/16819 | 0.000368607 | 0.024252755 | 0.022108076 | mt-Nd6/mt-Atp8/mt-Nd4/mt-Nd3/Gfpt1/Tk2 | 6 |
| **HP_ABNORMALITY_OF_THE_MUSCULATURE_OF_THE_LIMBS** | HP_ABNORMALITY_OF_THE_MUSCULATURE_OF_THE_LIMBS | 24/377 | 498/16819 | 0.000382597 | 0.024905406 | 0.022703013 | Rpl35/Ttn/Rps29/Rps28/Celf2/Rps26/Rps27/Rps15a/Dnm1l/Ndufc2/Rps17/Kdm1a/Dmd/Rpl31/Fgf10/Gfpt1/Adssl1/Plp1/Cryab/Ddhd1/Tk2/Fkrp/Lama2/Neb | 24 |
| **HP_HORSESHOE_KIDNEY** | HP_HORSESHOE_KIDNEY | 9/377 | 100/16819 | 0.000410137 | 0.026417125 | 0.02408105 | Rpl35/Rps29/Rps28/Rps26/Rps27/Rps15a/Hras/Rps17/Rpl31 | 9 |
| **GOBP_SYNAPSE_ORGANIZATION** | GOBP_SYNAPSE_ORGANIZATION | 21/377 | 412/16819 | 0.000425394 | 0.027114465 | 0.024716724 | Mef2c/Sipa1l1/Clstn2/Sez6/Adnp/Lrrc4c/Tnr/Zfp804a/Nrcam/Epha7/Kirrel3/Tmem108/Actr2/Il1rapl1/Ptprd/Pgrmc1/Drp2/Nrp2/Bcan/Robo2/Cdh8 | 21 |
| **HP_TRIANGULAR_SHAPED_PHALANGES_OF_THE_HAND** | HP_TRIANGULAR_SHAPED_PHALANGES_OF_THE_HAND | 14/377 | 219/16819 | 0.000438815 | 0.027681523 | 0.025233637 | Rpl35/Mafb/Rps29/Rps28/Rtl1/Rps26/Rps27/Rps15a/Adnp/Slc12a2/Rps17/Rpl31/Sf3b4/Fgf10 | 14 |
| **HP_GENERALIZED_ONSET_MOTOR_SEIZURE** | HP_GENERALIZED_ONSET_MOTOR_SEIZURE | 13/377 | 195/16819 | 0.000468892 | 0.029277036 | 0.026688057 | mt-Nd6/Scn1b/Scn9a/mt-Nd4/Dnm1l/Sms/Pcyt2/Pura/Hcn2/mt-Nd3/Kirrel3/Trpm3/Lnpk | 13 |
| **GOBP_REGULATION_OF_POSTSYNAPSE_ORGANIZATION** | GOBP_REGULATION_OF_POSTSYNAPSE_ORGANIZATION | 8/377 | 82/16819 | 0.000497013 | 0.030394051 | 0.027706294 | Sipa1l1/Zfp804a/Nrcam/Epha7/Actr2/Il1rapl1/Ptprd/Nrp2 | 8 |
| **HP_POOR_MOTOR_COORDINATION** | HP_POOR_MOTOR_COORDINATION | 7/377 | 63/16819 | 0.000505484 | 0.030394051 | 0.027706294 | Snrpn/Scn1b/Ndn/Scn9a/Rtl1/Fkrp/Dcc | 7 |
| **GOBP_NEGATIVE_REGULATION_OF_DEVELOPMENTAL_GROWTH** | GOBP_NEGATIVE_REGULATION_OF_DEVELOPMENTAL_GROWTH | 9/377 | 103/16819 | 0.000510273 | 0.030394051 | 0.027706294 | Spp1/Mt3/Tnr/Epha7/Sema3a/Sema5a/Dcc/Rgs2/Sema6a | 9 |
| **HP_RENAL_AGENESIS** | HP_RENAL_AGENESIS | 12/377 | 172/16819 | 0.000512375 | 0.030394051 | 0.027706294 | Rpl35/Rps29/Rps28/Rps26/Rps27/Rps15a/Rps17/Rpl31/Sf3b4/Fgf10/Sema3a/Dcc | 12 |
| **GOBP_REGULATION_OF_CELL_JUNCTION_ASSEMBLY** | GOBP_REGULATION_OF_CELL_JUNCTION_ASSEMBLY | 13/377 | 197/16819 | 0.000516585 | 0.030394051 | 0.027706294 | Mef2c/Clstn2/Slk/Adnp/Epha7/Map4k4/Dapk3/Il1rapl1/Ptprd/Dlc1/S100a10/Robo2/Apod | 13 |
| **GOMF_VOLTAGE_GATED_CHANNEL_ACTIVITY** | GOMF_VOLTAGE_GATED_CHANNEL_ACTIVITY | 13/377 | 197/16819 | 0.000516585 | 0.030394051 | 0.027706294 | Scn1b/Kcnab1/Scn9a/Kcnd2/Cacna1c/Hcn2/Tmem37/Ano6/Kcns3/Cacna1e/Kcnip4/Cachd1/Kcnc2 | 13 |
| **GOCC_NEURON_SPINE** | GOCC_NEURON_SPINE | 12/377 | 173/16819 | 0.000539702 | 0.031451766 | 0.028670476 | Rph3a/Kcnd2/Sipa1l1/Sez6/Mt3/Zfp804a/Calb1/Ppp1r9a/Cryab/Pde4b/Palmd/Slc8a1 | 12 |
| **HP_APLASIA_HYPOPLASIA_OF_FINGERS** | HP_APLASIA_HYPOPLASIA_OF_FINGERS | 18/377 | 334/16819 | 0.000567673 | 0.032769736 | 0.029871897 | Scn1b/Rpl35/Mafb/Rps29/Rps28/Rtl1/Rps26/Rps27/Rps15a/Rps17/Kdm1a/Dmd/Rpl31/Sf3b4/Fgf10/Il1rapl1/Stambp/Eif2ak3 | 18 |
| **HP_ABNORMALITY_OF_THE_CERVICAL_SPINE** | HP_ABNORMALITY_OF_THE_CERVICAL_SPINE | 20/377 | 393/16819 | 0.000591373 | 0.033803722 | 0.030814447 | Snrpn/Rpl35/Ankrd11/Mafb/Rps29/Rps28/Rtl1/Rps26/Rps27/Rps15a/Hras/Rps17/Rpl31/Sf3b4/Sema5a/Fkrp/Dpyd/Ripply2/Dcc/Neb | 20 |
| **GOBP_NEGATIVE_CHEMOTAXIS** | GOBP_NEGATIVE_CHEMOTAXIS | 6/377 | 47/16819 | 0.000602158 | 0.033803722 | 0.030814447 | Epha7/Sema3a/Sema5a/Nrp2/Sema6a/Robo2 | 6 |
| **HP_POOR_FINE_MOTOR_COORDINATION** | HP_POOR_FINE_MOTOR_COORDINATION | 6/377 | 47/16819 | 0.000602158 | 0.033803722 | 0.030814447 | Snrpn/Scn1b/Ndn/Scn9a/Rtl1/Dcc | 6 |
| **GOBP_REGULATION_OF_ION_TRANSMEMBRANE_TRANSPORT** | GOBP_REGULATION_OF_ION_TRANSMEMBRANE_TRANSPORT | 22/377 | 455/16819 | 0.000636212 | 0.03539076 | 0.032261143 | Scn1b/Kcnab1/Mef2c/Scn9a/Kcnd2/Cacna1c/Fxyd6/Dmd/Cox17/Cd63/Tmem37/Plp1/Ano6/Kcns3/Pde4b/Cacna1e/Utrn/Rgs2/Kcnip4/Dpp6/Slc8a1/Kcnc2 | 22 |
| **GOBP_REGULATION_OF_CELL_DEVELOPMENT** | GOBP_REGULATION_OF_CELL_DEVELOPMENT | 23/377 | 487/16819 | 0.000665291 | 0.036674914 | 0.033431739 | Hdac9/Spp1/Adnp/Mt3/Tnr/Rab21/Kdm1a/Epha7/Stk25/Actr2/Sema3a/Sema5a/Zdhhc21/Il1rapl1/Id4/Ptprd/Id1/Dcc/S100a10/Tenm4/Sema6a/Hey2/Robo2 | 23 |
| **HP_DIFFICULTY_CLIMBING_STAIRS** | HP_DIFFICULTY_CLIMBING_STAIRS | 6/377 | 48/16819 | 0.000675368 | 0.036768926 | 0.033517438 | Ttn/Dmd/Gfpt1/Adssl1/Fkrp/Lama2 | 6 |
| **GOBP_REGULATION_OF_DEVELOPMENTAL_GROWTH** | GOBP_REGULATION_OF_DEVELOPMENTAL_GROWTH | 17/377 | 311/16819 | 0.000684626 | 0.036768926 | 0.033517438 | Mef2c/Spp1/Adnp/Basp1/Tnks2/Mt3/Tnr/Rab21/Nrcam/Epha7/Sema3a/Sema5a/Ghr/Dcc/Rgs2/Sema6a/Hey2 | 17 |
| **HP_INFANTILE_MUSCULAR_HYPOTONIA** | HP_INFANTILE_MUSCULAR_HYPOTONIA | 13/377 | 203/16819 | 0.000685023 | 0.036768926 | 0.033517438 | Maf/mt-Nd6/Snrpn/Scn1b/Scn9a/Rtl1/mt-Nd4/Adnp/Sms/mt-Nd3/Atp10a/Fkrp/Dpyd | 13 |
| **GOBP_RESPIRATORY_ELECTRON_TRANSPORT_CHAIN** | GOBP_RESPIRATORY_ELECTRON_TRANSPORT_CHAIN | 9/377 | 108/16819 | 0.000721226 | 0.037719516 | 0.034383966 | mt-Nd6/mt-Nd4l/Ghitm/mt-Nd4/Ndufc2/Cycs/mt-Nd3/Cox7c/Uqcr10 | 9 |
| **GOCC_LARGE_RIBOSOMAL_SUBUNIT** | GOCC_LARGE_RIBOSOMAL_SUBUNIT | 9/377 | 108/16819 | 0.000721226 | 0.037719516 | 0.034383966 | Rpl39/Rpl35/Rpl37/Rpl21/Rpl37a/Rpl38/Rpl36/Rpl36a/Rpl31 | 9 |
| **HP_PSYCHOSIS** | HP_PSYCHOSIS | 9/377 | 108/16819 | 0.000721226 | 0.037719516 | 0.034383966 | mt-Nd6/Snrpn/Ndn/Spp1/mt-Nd4/Tbc1d7/Ttc19/Cdh23/Pus3 | 9 |
| **GOCC_SARCOLEMMA** | GOCC_SARCOLEMMA | 10/377 | 131/16819 | 0.000740536 | 0.038401205 | 0.035005373 | Scn1b/Cacna1c/Synm/Dmd/Ppp3r1/Sgcz/Fkrp/Utrn/Lama2/Slc8a1 | 10 |
| **GOBP_AEROBIC_RESPIRATION** | GOBP_AEROBIC_RESPIRATION | 12/377 | 180/16819 | 0.000767474 | 0.039463657 | 0.035973873 | mt-Nd6/mt-Atp8/mt-Nd4l/Ghitm/mt-Nd4/Ndufc2/Bloc1s1/Cycs/mt-Nd3/Cox7c/Uqcr10/Ide | 12 |
| **GOBP_OXIDATIVE_PHOSPHORYLATION** | GOBP_OXIDATIVE_PHOSPHORYLATION | 10/377 | 132/16819 | 0.000785622 | 0.040060168 | 0.036517633 | mt-Nd6/mt-Atp8/mt-Nd4l/Ghitm/mt-Nd4/Ndufc2/Cycs/mt-Nd3/Cox7c/Uqcr10 | 10 |
| **GOMF_MRNA_3_UTR_BINDING** | GOMF_MRNA_3_UTR_BINDING | 8/377 | 88/16819 | 0.000798065 | 0.040358345 | 0.036789442 | Pcbp4/Rbms3/Celf2/Carhsp1/Hnrnpa0/Cpeb3/Nova1/Zfp36l1 | 8 |
| **HP_BILATERAL_PTOSIS** | HP_BILATERAL_PTOSIS | 6/377 | 50/16819 | 0.000842311 | 0.042069402 | 0.03834919 | Maf/Adnp/Dnm1l/Auts2/Kdm1a/Tk2 | 6 |
| **GOBP_NEGATIVE_REGULATION_OF_CELL_GROWTH** | GOBP_NEGATIVE_REGULATION_OF_CELL_GROWTH | 12/377 | 182/16819 | 0.000845651 | 0.042069402 | 0.03834919 | Spp1/Mt3/Tnr/Phb/Eno1/Epha7/Sema3a/Cryab/Sema5a/Dcc/Rgs2/Sema6a | 12 |
| **HP_LETHARGY** | HP_LETHARGY | 13/377 | 208/16819 | 0.00085872 | 0.042375067 | 0.038627825 | Rpl35/Rps29/Rps28/Atp5md/Rps26/Rps27/Rps15a/Rps17/mt-Nd3/Rpl31/Hmgcl/Dpyd/Cdh23 | 13 |
| **HP_ABNORMAL_SUBARACHNOID_SPACE_MORPHOLOGY** | HP_ABNORMAL_SUBARACHNOID_SPACE_MORPHOLOGY | 10/377 | 134/16819 | 0.000882581 | 0.043204086 | 0.039383534 | mt-Nd6/Snrpn/Prkg1/mt-Nd4/Dmd/Kirrel3/Trpm3/Il1rapl1/Dcc/Hey2 | 10 |
| **HP_NEOPLASM_OF_THE_SKELETAL_SYSTEM** | HP_NEOPLASM_OF_THE_SKELETAL_SYSTEM | 8/377 | 91/16819 | 0.000995755 | 0.048033093 | 0.04378551 | Rpl35/Rps29/Rps28/Rps26/Rps27/Rps15a/Rps17/Rpl31 | 8 |
| **NABA_CORE_MATRISOME** | NABA_CORE_MATRISOME | 15/377 | 266/16819 | 0.001020482 | 0.048033093 | 0.04378551 | Efemp1/Spp1/Tnr/Igfbp6/Hmcn1/Crim1/Lgi1/Col19a1/Col9a3/Bcan/Lama2/Igfbp5/Edil3/Ogn/Coch | 15 |
| **GOCC_CATION_CHANNEL_COMPLEX** | GOCC_CATION_CHANNEL_COMPLEX | 13/377 | 212/16819 | 0.001023026 | 0.048033093 | 0.04378551 | Scn1b/Kcnab1/Scn9a/Kcnd2/Cacna1c/Hcn2/Kcns3/Pde4b/Cacna1e/Kcnip4/Dpp6/Cachd1/Kcnc2 | 13 |
| **GOBP_REGULATION_OF_TRANS_SYNAPTIC_SIGNALING** | GOBP_REGULATION_OF_TRANS_SYNAPTIC_SIGNALING | 20/377 | 411/16819 | 0.001026683 | 0.048033093 | 0.04378551 | Mef2c/Sipa1l1/Clstn2/Car2/Adnp/Lrrc4c/Hras/Tnr/Slc12a2/S100b/Tmem108/Car7/Calb1/Ppp1r9a/Cpeb3/Lgi1/Ptprd/Dcc/Lama2/Dlgap2 | 20 |
| **GOBP_INSULIN_LIKE_GROWTH_FACTOR_RECEPTOR_SIGNALING_PATHWAY** | GOBP_INSULIN_LIKE_GROWTH_FACTOR_RECEPTOR_SIGNALING_PATHWAY | 5/377 | 35/16819 | 0.001028327 | 0.048033093 | 0.04378551 | Igfbp6/Crim1/Ghr/Eif2ak3/Igfbp5 | 5 |
| **GOBP_MEMBRANE_DEPOLARIZATION_DURING_ACTION_POTENTIAL** | GOBP_MEMBRANE_DEPOLARIZATION_DURING_ACTION_POTENTIAL | 5/377 | 35/16819 | 0.001028327 | 0.048033093 | 0.04378551 | Scn1b/Scn9a/Cacna1c/Hcn2/Slc8a1 | 5 |
