## Supplemental Table 1 for "Combinatorial transcriptional regulation establishes subtype-appropriate synaptic properties in auditory neurons"

| ABR Threshold post-hoc Dunn |  |  |  |  |  |  |
| --- | --- | --- | --- | --- | --- | --- |
| Threshold_8kHz | Kruskal with posthoc Dunn |  | **DKO** | **CMAFCKO** | **NOCRE** | **MAFBCKO** |
| KruskalResult(statistic=19.322111934627316, pvalue=0.00023451450641378163) |  | **DKO** | 1 | 0.00037 | 0.000073 | 0.004833 |
|  |  | **CMAFCKO** | 0.00037 | 1 | 0.514821 | 0.133246 |
|  |  | **NOCRE** | 0.000073 | 0.514821 | 1 | 0.220367 |
|  |  | **MAFBCKO** | 0.004833 | 0.133246 | 0.220367 | 1 |
| Threshold_16kHz | Kruskal with posthoc Dunn |  | **DKO** | **CMAFCKO** | **NOCRE** | **MAFBCKO** |
| KruskalResult(statistic=21.382743193637186, pvalue=8.766259814282283e-05) |  | **DKO** | 1 | 0.000165 | 0.000114 | 0.049639 |
|  |  | **CMAFCKO** | 0.000165 | 1 | 0.240438 | 0.011924 |
|  |  | **NOCRE** | 0.000114 | 0.240438 | 1 | 0.035543 |
|  |  | **MAFBCKO** | 0.049639 | 0.011924 | 0.035543 | 1 |
| Frequency: 32 kHz: |  |  |  |  |  |  |
| Stat: 5.206 P val: 0.157 |  |  |  |  |  |  |
| Frequency: 45 kHz: |  |  |  |  |  |  |
| Stat: 3.725 P val: 0.293 |  |  |  |  |  |  |
| Amplitude post-hoc Dunn |  |  |  |  |  |  |
| 16_90_P1AMP | Kruskal with posthoc Dunn |  | **NOCRE** | **dKO** | **CMAFCKO** | **MAFBCKO** |
| Kruskal(Stat: 23.319, pvalue=3.465288275835971e-05) |  | **NOCRE** | 1 | 0.000014 | 0.518054 | 0.057026 |
|  |  | **dKO** | 0.000014 | 1 | 0.000213 | 0.01513 |
|  |  | **CMAFCKO** | 0.518054 | 0.000213 | 1 | 0.052272 |
|  |  | **MAFBCKO** | 0.057026 | 0.01513 | 0.052272 | 1 |
| 16_85_P1AMP | Kruskal with post-hoc Dunn and bonferonni |  | **NOCRE** | **dKO** | **CMAFCKO** | **MAFBCKO** |
| Kruskal(Stat: 22.704, pvalue=4.6544621465723745e-05) |  | **NOCRE** | 1 | 0.000017 | 0.576526 | 0.041912 |
|  |  | **dKO** | 0.000017 | 1 | 0.000333 | 0.023241 |
|  |  | **CMAFCKO** | 0.576526 | 0.000333 | 1 | 0.051638 |
|  |  | **MAFBCKO** | 0.041912 | 0.023241 | 0.051638 | 1 |
| 16_80_P1AMP | Kruskal with posthoc Dunn |  | **NOCRE** | **dKO** | **CMAFCKO** | **MAFBCKO** |
| Kruskal(Stat: 22.962, pvalue=3.192078002118693e-05) |  | **NOCRE** | 1 | 0.000022 | 0.478381 | 0.034824 |
|  |  | **dKO** | 0.000022 | 1 | 0.000227 | 0.032063 |
|  |  | **CMAFCKO** | 0.478381 | 0.000227 | 1 | 0.031979 |
|  |  | **MAFBCKO** | 0.034824 | 0.032063 | 0.031979 | 1 |
| 16_75_P1AMP | Kruskal with posthoc Dunn |  | **NOCRE** | **dKO** | **CMAFCKO** | **MAFBCKO** |
| Kruskal(Stat: 23.49, pvalue=3.192078002118693e-05) |  | **NOCRE** | 1 | 0.00001 | 0.615771 | 0.046859 |
|  |  | **dKO** | 0.00001 | 1 | 0.000297 | 0.015411 |
|  |  | **CMAFCKO** | 0.615771 | 0.000297 | 1 | 0.063047 |
|  |  | **MAFBCKO** | 0.046859 | 0.015411 | 0.063047 | 1 |
| 16_70_P1AMP | Kruskal with posthoc Dunn |  | **NOCRE** | **dKO** | **CMAFCKO** | **MAFBCKO** |
| Kruskal(Stat: 23.571, pvalue=3.0691892781457156e-05) |  | **NOCRE** | 1 | 0.000011 | 0.578442 | 0.038789 |
|  |  | **dKO** | 0.000011 | 1 | 0.000263 | 0.020061 |
|  |  | **CMAFCKO** | 0.578442 | 0.000263 | 1 | 0.049355 |
|  |  | **MAFBCKO** | 0.038789 | 0.020061 | 0.049355 | 1 |
| 16_65_P1Amp | Kruskal with posthoc Dunn |  | **NOCRE** | **dKO** | **CMAFCKO** | **MAFBCKO** |
| Kruskal(Stat: 20.796, pvalue=0.00011606240622921905) |  | **NOCRE** | 1 | 0.000039 | 0.578358 | 0.05427 |
|  |  | **dKO** | 0.000039 | 1 | 0.000561 | 0.028807 |
|  |  | **CMAFCKO** | 0.578358 | 0.000561 | 1 | 0.061787 |
|  |  | **MAFBCKO** | 0.05427 | 0.028807 | 0.061787 | 1 |
| 16_60_P1Amp | Kruskal with posthoc Dunn |  | **NOCRE** | **dKO** | **CMAFCKO** | **MAFBCKO** |
| Kruskal(Stat:20.731, pvalue=0.00011974163729525886) |  | **NOCRE** | 1 | 0.000107 | 0.401171 | 0.0243 |
|  |  | **dKO** | 0.000107 | 1 | 0.00039 | 0.096218 |
|  |  | **CMAFCKO** | 0.401171 | 0.00039 | 1 | 0.017814 |
|  |  | **MAFBCKO** | 0.0243 | 0.096218 | 0.017814 | 1 |
| 16_55_P1Amp | Kruskal with posthoc Dunn |  | **NOCRE** | **dKO** | **CMAFCKO** | **MAFBCKO** |
| Kruskal(Stat: 20.045, pvalue=0.0001661209600006024) |  | **NOCRE** | 1 | 0.000097 | 0.431925 | 0.047036 |
|  |  | **dKO** | 0.000097 | 1 | 0.000444 | 0.054336 |
|  |  | **CMAFCKO** | 0.431925 | 0.000444 | 1 | 0.032878 |
|  |  | **MAFBCKO** | 0.047036 | 0.054336 | 0.032878 | 1 |
| 16_50_P1Amp | Kruskal with posthoc Dunn |  | **NOCRE** | **dKO** | **CMAFCKO** | **MAFBCKO** |
| Kruskal(21.294, pvalue=9.145856879761562e-05) |  | **NOCRE** | 1 | 0.000049 | 0.401119 | 0.088387 |
|  |  | **dKO** | 0.000049 | 1 | 0.000233 | 0.019619 |
|  |  | **CMAFCKO** | 0.401119 | 0.000233 | 1 | 0.046467 |
|  |  | **MAFBCKO** | 0.088387 | 0.019619 | 0.046467 | 1 |
| 16_45_P1Amp | Kruskal with posthoc Dunn |  | **NOCRE** | **dKO** | **CMAFCKO** | **MAFBCKO** |
| Kruskal(Stat: 17.367, pvalue=0.0005938602940773141) |  | **NOCRE** | 1 | 0.001379 | 0.164276 | 0.071941 |
|  |  | **dKO** | 0.001379 | 1 | 0.000335 | 0.153351 |
|  |  | **CMAFCKO** | 0.164276 | 0.000335 | 1 | 0.009534 |
|  |  | **MAFBCKO** | 0.071941 | 0.153351 | 0.009534 | 1 |
| 16_40_P1Amp | Kruskal with posthoc Dunn |  | **NOCRE** | **dKO** | **CMAFCKO** | **MAFBCKO** |
| Kruskal(Stat: 17.737, pvalue=0.0004984463775650707) |  | **NOCRE** | 1 | 0.000566 | 0.304073 | 0.044838 |
|  |  | **dKO** | 0.000566 | 1 | 0.000624 | 0.139207 |
|  |  | **CMAFCKO** | 0.304073 | 0.000624 | 1 | 0.017203 |
|  |  | **MAFBCKO** | 0.044838 | 0.139207 | 0.017203 | 1 |
| 16_35_P1Amp | Kruskal with posthoc Dunn |  | **NOCRE** | **dKO** | **CMAFCKO** | **MAFBCKO** |
| Kruskal(Stat: 15.344, pvalue=0.0015452763623379155) |  | **NOCRE** | 1 | 0.000824 | 0.545277 | 0.035636 |
|  |  | **dKO** | 0.000824 | 1 | 0.003193 | 0.195076 |
|  |  | **CMAFCKO** | 0.545277 | 0.003193 | 1 | 0.041632 |
|  |  | **MAFBCKO** | 0.035636 | 0.195076 | 0.041632 | 1 |
| 16_30_P1Amp | Kruskal with posthoc Dunn |  | **NOCRE** | **dKO** | **CMAFCKO** | **MAFBCKO** |
| Kruskal(Stat: 12.582, pvalue=0.005634508659930855) |  | **NOCRE** | 1 | 0.004587 | 0.229654 | 0.469733 |
|  |  | **dKO** | 0.004587 | 1 | 0.00164 | 0.040707 |
|  |  | **CMAFCKO** | 0.229654 | 0.00164 | 1 | 0.095856 |
|  |  | **MAFBCKO** | 0.469733 | 0.040707 | 0.095856 | 1 |
| 16_25_P1Amp | Kruskal | pvalue=0.14677591035838353 |  |  |  |  |
| 16_20_P1Amp | Kruskal | pvalue=0.8130613102888596 |  |  |  |  |
| 8_90_P1Amp | Kruskal with posthoc Dunn |  | **NOCRE** | **dKO** | **CMAFCKO** | **MAFBCKO** |
|  |  | **NOCRE** | 1 | 0.000076 | 0.32956 | 0.187395 |
|  |  | **dKO** | 0.000076 | 1 | 0.000065 | 0.008933 |
|  |  | **CMAFCKO** | 0.32956 | 0.000065 | 1 | 0.050865 |
|  |  | **MAFBCKO** | 0.187395 | 0.008933 | 0.050865 | 1 |
| 8_85_P1Amp | Kruskal with posthoc Dunn |  | **NOCRE** | **dKO** | **CMAFCKO** | **MAFBCKO** |
|  |  | **NOCRE** | 1 | 0.000095 | 0.356363 | 0.235574 |
|  |  | **dKO** | 0.000095 | 1 | 0.000094 | 0.007247 |
|  |  | **CMAFCKO** | 0.356363 | 0.000094 | 1 | 0.071735 |
|  |  | **MAFBCKO** | 0.235574 | 0.007247 | 0.071735 | 1 |
| 8_80_P1Amp | Kruskal with posthoc Dunn |  | **NOCRE** | **dKO** | **CMAFCKO** | **MAFBCKO** |
|  |  | **NOCRE** | 1 | 0.000042 | 0.51354 | 0.184382 |
|  |  | **dKO** | 0.000042 | 1 | 0.000179 | 0.006893 |
|  |  | **CMAFCKO** | 0.51354 | 0.000179 | 1 | 0.104662 |
|  |  | **MAFBCKO** | 0.184382 | 0.006893 | 0.104662 | 1 |
| 8_75_P1Amp | Kruskal with posthoc Dunn |  | **NOCRE** | **dKO** | **CMAFCKO** | **MAFBCKO** |
|  |  | **NOCRE** | 1 | 0.000034 | 0.575791 | 0.132496 |
|  |  | **dKO** | 0.000034 | 1 | 0.000217 | 0.009548 |
|  |  | **CMAFCKO** | 0.575791 | 0.000217 | 1 | 0.096063 |
|  |  | **MAFBCKO** | 0.132496 | 0.009548 | 0.096063 | 1 |
| 8_70_P1Amp | Kruskal with posthoc Dunn |  | **NOCRE** | **dKO** | **CMAFCKO** | **MAFBCKO** |
|  |  | **NOCRE** | 1 | 0.000214 | 0.181282 | 0.306813 |
|  |  | **dKO** | 0.000214 | 1 | 0.000044 | 0.009255 |
|  |  | **CMAFCKO** | 0.181282 | 0.000044 | 1 | 0.039909 |
|  |  | **MAFBCKO** | 0.306813 | 0.009255 | 0.039909 | 1 |
| 8_65_P1Amp | Kruskal with posthoc Dunn |  | **NOCRE** | **dKO** | **CMAFCKO** | **MAFBCKO** |
|  |  | **NOCRE** | 1 | 0.000121 | 0.262228 | 0.319316 |
|  |  | **dKO** | 0.000121 | 1 | 0.000066 | 0.00577 |
|  |  | **CMAFCKO** | 0.262228 | 0.000066 | 1 | 0.067718 |
|  |  | **MAFBCKO** | 0.319316 | 0.00577 | 0.067718 | 1 |
| 8_60_P1Amp | Kruskal with posthoc Dunn |  | **NOCRE** | **dKO** | **CMAFCKO** | **MAFBCKO** |
|  |  | **NOCRE** | 1 | 0.000041 | 0.223316 | 0.143964 |
|  |  | **dKO** | 0.000041 | 1 | 0.000019 | 0.009658 |
|  |  | **CMAFCKO** | 0.223316 | 0.000019 | 1 | 0.023341 |
|  |  | **MAFBCKO** | 0.143964 | 0.009658 | 0.023341 | 1 |
| 8_55_P1Amp | Kruskal with posthoc Dunn |  | **NOCRE** | **dKO** | **CMAFCKO** | **MAFBCKO** |
|  |  | **NOCRE** | 1 | 0.000657 | 0.301055 | 0.136722 |
|  |  | **dKO** | 0.000657 | 1 | 0.00035 | 0.056862 |
|  |  | **CMAFCKO** | 0.301055 | 0.00035 | 1 | 0.034807 |
|  |  | **MAFBCKO** | 0.136722 | 0.056862 | 0.034807 | 1 |
| 8_50_P1Amp | Kruskal with posthoc Dunn |  | **NOCRE** | **dKO** | **CMAFCKO** | **MAFBCKO** |
|  |  | **NOCRE** | 1 | 0.001043 | 0.490605 | 0.227433 |
|  |  | **dKO** | 0.001043 | 1 | 0.001603 | 0.04188 |
|  |  | **CMAFCKO** | 0.490605 | 0.001603 | 1 | 0.116994 |
|  |  | **MAFBCKO** | 0.227433 | 0.04188 | 0.116994 | 1 |
| 8_45_P1Amp | Kruskal with posthoc Dunn |  | **NOCRE** | **dKO** | **CMAFCKO** | **MAFBCKO** |
|  |  | **NOCRE** | 1 | 0.000128 | 0.623694 | 0.25547 |
|  |  | **dKO** | 0.000128 | 1 | 0.000692 | 0.008739 |
|  |  | **CMAFCKO** | 0.623694 | 0.000692 | 1 | 0.185705 |
|  |  | **MAFBCKO** | 0.25547 | 0.008739 | 0.185705 | 1 |
| 8_40_P1Amp | Kruskal with posthoc Dunn |  | **NOCRE** | **dKO** | **CMAFCKO** | **MAFBCKO** |
|  |  | **NOCRE** | 1 | 0.208936 | 0.410973 | 0.913981 |
|  |  | **dKO** | 0.208936 | 1 | 0.083556 | 0.196631 |
|  |  | **CMAFCKO** | 0.410973 | 0.083556 | 1 | 0.475546 |
|  |  | **MAFBCKO** | 0.913981 | 0.196631 | 0.475546 | 1 |
| Latency post-hoc Dunn |  |  |  |  |  |  |
| 16_90_P1Lat | Kruskal with posthoc Dunn |  | **NOCRE** | **CMAFCKO** | **MAFBCKO** |  |
| Kruskal: p=2.29E-04 |  | **NOCRE** | 1 | 0.101105 | 0.000046 |  |
|  |  | **CMAFCKO** | 0.101105 | 1 | 0.221992 |  |
|  |  | **MAFBCKO** | 0.000046 | 0.221992 | 1 |  |
| 16_85_P1lat | Kruskal with posthoc Dunn |  | **NOCRE** | **CMAFCKO** | **MAFBCKO** |  |
| Kruskal: p=0.001 |  | **NOCRE** | 1 | 0.02357 | 0.000559 |  |
|  |  | **CMAFCKO** | 0.02357 | 1 | 0.85505 |  |
|  |  | **MAFBCKO** | 0.000559 | 0.85505 | 1 |  |
| 16_80_P1Lat | Kruskal with posthoc Dunn |  | **NOCRE** | **CMAFCKO** | **MAFBCKO** |  |
| Kruskal: p=0.002 |  | **NOCRE** | 1 | 0.032732 | 0.000659 |  |
|  |  | **CMAFCKO** | 0.032732 | 1 | 0.781838 |  |
|  |  | **MAFBCKO** | 0.000659 | 0.781838 | 1 |  |
| 16_75_P1Lat | Kruskal with posthoc Dunn |  | **NOCRE** | **CMAFCKO** | **MAFBCKO** |  |
| Kruskal: p=0.004 |  | **NOCRE** | 1 | 0.034074 | 0.002145 |  |
|  |  | **CMAFCKO** | 0.034074 | 1 | 0.951864 |  |
|  |  | **MAFBCKO** | 0.002145 | 0.951864 | 1 |  |
| 16_70_P1Lat | Kruskal with posthoc Dunn |  | **NOCRE** | **CMAFCKO** | **MAFBCKO** |  |
| Kruskal: p= 0.019 |  | **NOCRE** | 1 | 0.073684 | 0.00904 |  |
|  |  | **CMAFCKO** | 0.073684 | 1 | 0.948382 |  |
|  |  | **MAFBCKO** | 0.00904 | 0.948382 | 1 |  |
| Frequency: 16 dB: 65 |  | pvalue=0.1940904724613508 |  |  |  |  |
| Stat: 3.279 P val: 0.194 |  |  | Kruskal |  |  |  |
| Frequency: 16 dB: 60 |  | pvalue=0.1887420539126795 |  |  |  |  |
| Stat: 3.335 P val: 0.189 |  |  | Kruskal |  |  |  |
| Frequency: 16 dB: 55 |  | pvalue=0.37891076678707053 |  |  |  |  |
| Stat: 1.941 P val: 0.379 |  |  | Kruskal |  |  |  |
| Frequency: 16 dB: 50 |  |  |  |  |  |  |
| Stat: 2.569 P val: 0.277 |  |  | Kruskal |  |  |  |
| Frequency: 16 dB: 45 |  |  |  |  |  |  |
| Stat: 1.497 P val: 0.473 |  |  | Kruskal |  |  |  |
| Frequency: 16 dB: 40 |  |  |  |  |  |  |
| Stat: 1.585 P val: 0.453 |  |  | Kruskal |  |  |  |
| Frequency: 16 dB: 35 |  | pvalue=0.03338786875203418 |  |  |  |  |
| Stat: 6.799 P val: 0.033 |  |  |  |  |  |  |
| Frequency: 16 dB: 30 |  |  | Kruskal |  |  |  |
| Stat: 6.743 P val: 0.034 |  |  |  |  |  |  |
| Frequency: 16 dB: 25 |  |  | Kruskal |  |  |  |
| Stat: 11.309 P val: 0.004 |  |  |  |  |  |  |
| Frequency: 16 dB: 20 |  |  | Kruskal |  |  |  |
| Stat: 4.965 P val: 0.084 |  |  |  |  |  |  |
| 8_90_P1Lat |  |  | **NOCRE** | **CMAFCKO** | **MAFBCKO** |  |
|  |  | **NOCRE** | 1 | 0.161979 | 0.000405 |  |
|  |  | **CMAFCKO** | 0.161979 | 1 | 0.186154 |  |
|  |  | **MAFBCKO** | 0.000405 | 0.186154 | 1 |  |
| 8_85_P1Lat |  |  | **NOCRE** | **CMAFCKO** | **MAFBCKO** |  |
|  |  | **NOCRE** | 1 | 0.096574 | 0.000252 |  |
|  |  | **CMAFCKO** | 0.096574 | 1 | 0.246514 |  |
|  |  | **MAFBCKO** | 0.000252 | 0.246514 | 1 |  |
| 8_80_P1Lat |  |  | **NOCRE** | **CMAFCKO** | **MAFBCKO** |  |
|  |  | **NOCRE** | 1 | 0.157931 | 0.000995 |  |
|  |  | **CMAFCKO** | 0.157931 | 1 | 0.273443 |  |
|  |  | **MAFBCKO** | 0.000995 | 0.273443 | 1 |  |
| 8_75_P1Lat |  |  | **NOCRE** | **CMAFCKO** | **MAFBCKO** |  |
|  |  | **NOCRE** | 1 | 0.549463 | 0.000918 |  |
|  |  | **CMAFCKO** | 0.549463 | 1 | 0.057729 |  |
|  |  | **MAFBCKO** | 0.000918 | 0.057729 | 1 |  |
| 8_70_P1Lat |  |  | **NOCRE** | **CMAFCKO** | **MAFBCKO** |  |
|  |  | **NOCRE** | 1 | 0.206899 | 0.001546 |  |
|  |  | **CMAFCKO** | 0.206899 | 1 | 0.251854 |  |
|  |  | **MAFBCKO** | 0.001546 | 0.251854 | 1 |  |
| 8_65_P1Lat |  |  | **NOCRE** | **CMAFCKO** | **MAFBCKO** |  |
|  |  | **NOCRE** | 1 | 0.196813 | 0.016501 |  |
|  |  | **CMAFCKO** | 0.196813 | 1 | 0.586197 |  |
|  |  | **MAFBCKO** | 0.016501 | 0.586197 | 1 |  |
| Frequency: 8 dB: 60 |  |  | Kruskal |  |  |  |
| Stat: 4.655 P val: 0.098 |  |  |  |  |  |  |
| Frequency: 8 dB: 55 |  |  | Kruskal |  |  |  |
| Stat: 4.919 P val: 0.085 |  |  |  |  |  |  |
| Frequency: 8 dB: 50 |  |  | Kruskal |  |  |  |
| Stat: 3.21 P val: 0.201 |  |  |  |  |  |  |
| Frequency: 8 dB: 45 |  |  | Kruskal |  |  |  |
| Stat: 5.711 P val: 0.058 |  |  |  |  |  |  |
| Frequency: 8 dB: 40 |  |  | Kruskal |  |  |  |
| Stat: 4.383 P val: 0.112 |  |  |  |  |  |  |
| Frequency: 8 dB: 35 |  |  | Kruskal |  |  |  |
| Stat: 0.857 P val: 0.651 |  |  |  |  |  |  |
| Frequency: 8 dB: 30 |  |  | Kruskal |  |  |  |
| Stat: 1.924 P val: 0.382 |  |  |  |  |  |  |
| Frequency: 8 dB: 25 |  |  | Kruskal |  |  |  |
| Stat: 2.054 P val: 0.358 |  |  |  |  |  |  |
| Frequency: 8 dB: 20 |  |  | Kruskal |  |  |  |
| Stat: 3.936 P val: 0.14 |  |  |  |  |  |  |
| DPOAE |  |  |  |  |  |  |
| Frequency: 8.0 kHz: |  | Kruskal |  |  |  |  |
| Stat: 0.533 P val: 0.766 |  |  |  |  |  |  |
| Frequency: 11.3 kHz: |  | Kruskal |  |  |  |  |
| Stat: 1.989 P val: 0.575 |  |  |  |  |  |  |
| Frequency: 16.0 kHz: |  | Kruskal |  |  |  |  |
| Stat: 5.928 P val: 0.115 |  |  |  |  |  |  |
| Frequency: 22.6 kHz: |  | Kruskal |  |  |  |  |
| Stat: 1.3 P val: 0.729 |  |  |  |  |  |  |
| Frequency: 32.0 kHz: |  | Kruskal |  |  |  |  |
| Stat: 5.054 P val: 0.168 |  |  |  |  |  |  |
